# Dinucleotide preferences underlie apparent codon preference reversals in the *Drosophila melanogaster* lineage

**DOI:** 10.1101/2024.10.10.617326

**Authors:** Haruka Yamashita, Tomotaka Matsumoto, Kent Kawashima, Hassan Sibroe Abdulla Daanaa, Ziheng Yang, Hiroshi Akashi

**Affiliations:** Evolutionary Genetics Laboratory, National Institute of Genetics, Mishima, Shizuoka 411-8540, Japan; Department of Genetics, The Graduate University for Advanced Studies, SOKENDAI, Mishima, Shizuoka 411-8540, Japan; Department of Genetics, Evolution and Environment, University College London, London WC1E 6BT, United Kingdom

## Abstract

We employ fine-scale population genetic analyses to reveal dynamics among interacting forces that act at synonymous sites and introns among closely related Drosophila species. Synonymous codon usage bias has proven to be well-suited for population genetic inference. Under major codon preference, translationally superior “major” codons confer fitness benefits relative to their less efficiently and/or accurately decoded synonymous counterparts. Our codon family and lineage-specific analyses expand on previous findings in the *Drosophila simulans* lineage; patterns in naturally occurring polymorphism demonstrate fixation biases toward GC-ending codons that are consistent in direction, but heterogeneous in magnitude, among synonymous families. These forces are generally stronger than fixation biases in intron sequences. In contrast, population genetic analyses reveal unexpected evidence of codon preference reversals in the *Drosophila melanogaster* lineage. Codon family-specific polymorphism patterns support reduced efficacy of natural selection in most synonymous families but indicate reversals of favored states in the four codon families encoded by NAY. Accelerated synonymous fixations in favor of NAT and greater differences for both allele frequencies and fixation rates among X-linked, relative to autosomal, loci bolster support for fitness effect reversals. The specificity of preference reversals to codons whose cognate tRNAs undergo wobble position queuosine modification is intriguing. However, our analyses reveal prevalent dinucleotide preferences for ApT over ApC that act in opposition to GC favoring forces in both coding and intron regions. We present evidence that changes in the relative efficacy of translational selection and dinucleotide preference underlie apparent codon preference reversals.

## Introduction

Codon usage bias is a well-developed system for studying evolutionary dynamics under nearly neutral genome evolution. A combination of sequence patterns and experimental findings support “major codon preference” (MCP), co-adaptation between synonymous codon usage and cognate tRNA abundances and their modifications to allow efficient use of the translational machinery (*i.e.*, greater elongation rates and reduced misincorporation) (reviewed in Andersson and Kurland 1991). A wide range of microbes, as well as multi-cellular eukaryotes, show elevated usage of particular codons within synonymous families in highly expressed genes (reviewed in Akashi 2001 and Duret 2002). Such codons are generally recognized by abundant and/or non-wobble-pairing cognate tRNAs and are referred to as “major” codons (Ikemura 1985). Biochemical studies support both elevated elongation rates (Varenne et al. 1984; Curran and Yarus 1989; Dana and Tuller 2014) and reduced misincorporations (Precup and Parker 1987; Kramer and Farabaugh 2007) at major codons relative to their minor counterparts. Putative major codons can be identified through characterizations of tRNA pools (*e.g.*, Ikemura 1981a; Ikemura 1982), experimental measures of translation rate (*e.g.*, Varenne *et al*. 1984), or by compositional trends among genes (*e.g.*, Grantham *et al*. 1981; Shields *et al*. 1988; Sharp and Devine 1989; Lloyd and Sharp 1992; Stenico *et al*. 1994; Akashi 1995; Kanaya *et al*. 1999).

The main features of MCP can be captured in relatively simple evolutionary models that can guide data analyses. Observed levels of codon usage bias are consistent with a balance among weak evolutionary forces including mutation, genetic drift and weak natural selection (Li 1987; Bulmer 1991). Under MCP, mutations from minor to major codons confer small fitness benefits and mutations in the opposite direction are deleterious with similar magnitudes; population genetic approaches can test the model by contrasting evolutionary patterns between these predicted fitness classes (Akashi 1995). In addition, because translational effects of synonymous codon usage should be similar among genes, data can be pooled among loci to enhance the statistical power to detect minute fixation biases (*i.e.*, forces that alter expected allele frequencies from generation to generation in a consistent direction). These features facilitate the study of evolutionary dynamics under a balance of weak forces and previous population genetic analyses have demonstrated directional forces that favor major codons over their synonymous counterparts (*e.g.*, Akashi 1995; Akashi and Schaeffer 1997; Kliman 1999; Sharp *et al*. 2010; Jackson *et al*. 2017). GC-biased gene conversion can also underlie base compositional biases (Marais 2003; Duret and Galtier 2009) and distinguishing between natural selection and non-selective fixation biases has proven challenging. In addition, a number of empirical studies support beneficial effects of translational pauses in regulating protein folding or membrane insertion (reviewed in Komar 2021). Major codons may be disadvantageous in some contexts, but such cases may be more strongly selected and relatively rare.

Here, we take advantage of genome-scale data from population samples of closely related Drosophila species to pursue evolutionary analyses at an increased resolution. We test fixation biases within individual synonymous families both among mutations segregating in populations and among fixations in ancestral lineages. Our analyses support MCP favoring predominantly G- and C-ending codons in *Drosophila simulans* but reveal appreciable heterogeneity in the magnitude of selective forces among synonymous families. In contrast, the *Drosophila melanogaster* lineage shows little evidence of fixation biases on synonymous mutations for most codon families. Population genetics theory predicts long-term stability of codon preferences under co-adaptation between codon usage and tRNA pools, but surprisingly, we found that a subset of synonymous families, those encoded by NAY codons, show strong evidence for recent codon preference reversals in the *D. melanogaster* lineage. These findings, together with greater contrasts in both the site frequency spectrum and fixation patterns at X-linked, relative to autosomal, loci make a compelling case for genome-wide shifts in NAY codon family adaptation. Intriguingly, all NAY codons, and no other codons, are recognized by cognate tRNAs that undergo a chemical (queuosine) modification within the anti-codon region. The chemical precursor for this modification, queine, is obtained through the environment; diet- or habitat-based shifts in major codon definition have been suggested for other Drosophila lineages (Powell *et al*. 2003; Zaborske *et al*. 2014) and seemed to be an attractive scenario to account for the patterns observed in this study.

However, NAT preference at NAY codons could also reflect dinucleotide ApT preference over ApC. We show patterns at synonymous codons as well as intron sites that support prevalent dinucleotide preference in coding and non-coding regions. ApT over ApC preference acts in opposition to GC fixation biases (including MCP) in both *D. melanogaster* and *D. simulans*. In the *D. melanogaster* lineage, reductions in the efficacy of GC fixation bias have shifted the balance of forces toward relatively stronger context-dependent T over C preference leading to *apparent* codon preference reversals at NAY codons. Although similar opposing forces act in *D. simulans*, the greater relative strength of GC favoring factors, including major codon preference, can mask the signals of ApT over ApC preference. Our population genetic analyses reveal a dynamic balance among fixation biases underlying codon and base composition evolution among closely related Drosophila species.

## Results and Discussion

We analyzed available population genomic DNA sequence data among closely related species from the *Drosophila melanogaster* subgroup. We constructed CDS and intron sequence alignments for 21 *D. simulans* lines from a Madagascar population (Rogers *et al*. 2014; Jackson *et al*. 2017) and 14 *D. melanogaster* lines from a Rwandan population (Pool *et al*. 2012). Reference sequences for *D. yakuba* and *D. erecta* (Drosophila 12 Genomes Consortium 2007), were used as outgroups (Fig. 1a). Classifying synonymous mutations by their predicted fitness effects, a key step for population genetic analyses of codon usage (Akashi 1995), requires inference of ancestral and derived states at polymorphic sites. The relatively short branch lengths in our tree allows reliable inference using a likelihood based method that incorporates both biased and fluctuating base composition (Matsumoto et al. 2015; Matsumoto and Akashi 2018; see Materials and Methods).

**Fig. 1.**
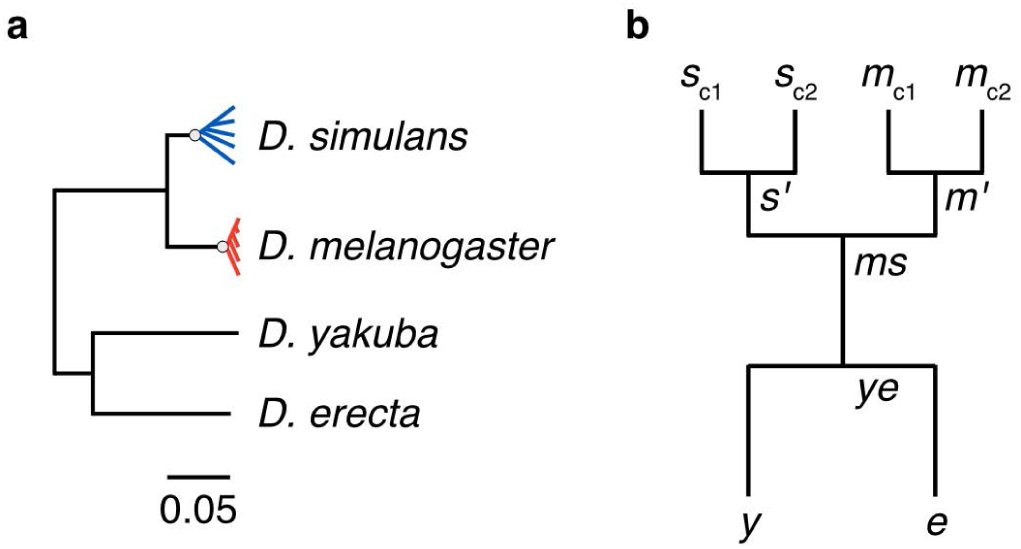
Gene tree of *Drosophila melanogaster* subgroup species employed in this study. Population genetic analyses employed data from *D. simulans* (*Dsim*), *D. melanogaster* (*Dmel*), and two outgroups, *D. yakuba* (*Dyak*) and *D. erecta* (*Dere*); only species included in our analyses are not shown. (a) Genetic distances among DNA sequences from the four species. Branch lengths are numbers of nucleotide differences per site at autosomal short introns. The within-species gene trees are rough depictions based on the strong and weaker excesses of rare polymorphisms (“star-like” trees) compared to neutral equilibrium expectations in *Dmel* and *Dsim*, respectively. Gray circles indicate approximate positions of the MRCA within each population sample. If a single allele is sampled from a population, a proportion of the changes inferred on the terminal branch will occur at polymorphic sites; the post-MRCA distances reflect the expected numbers of such changes assuming independence among observed polymorphisms. (b) Tree topology employed for ancestral inference. Within-species variation was assigned to two sequences (*m*_c1_ and *m*_c2_ for *Dmel*, *s*_c1_ and *s*_c2_ for *Dsim,* see Materials and Methods). Ancestral nodes for within-species sequences are indicated as *s’* for *Dsim* and *m’* for *Dmel*. Ancestral nodes for between-species sequences are labeled *ms* and *ye* for the pairs *Dmel* / *Dsim* and *Dyak* / *Dere*, respectively. This topology is assumed for ancestral inference (note that branch lengths are estimated in the process).

### Inferring fixation biases from D. simulans and D. melanogaster polymorphism

Directional forces impact allele frequencies of mutations segregating within populations. We can infer such forces by examining distributions of allele frequencies among variants segregating in a population, the site frequency spectrum (SFS). Direct comparisons of SFS between classes of mutations that are interspersed within DNA (Bulmer 1971; Sawyer *et al*. 1987) can be a robust statistical approach that is sensitive to weak forces (Akashi and Schaeffer 1997; Akashi 1999) including natural selection and GC-biased gene conversion (gBGC).

SFS comparisons support directional forces favoring G/C over A/T at both 2-fold and 4-fold redundant sites in *D. simulans* (Table S5). The results expand on previous findings that combined data from 2-fold and 4-fold synonymous families for smaller numbers of genes (Akashi and Schaeffer 1997; Kliman 1999) and that employed only 4-fold synonymous families in larger data sets (Jackson *et al*. 2017). These comparisons distinguish between mutational and fixation biases because the former affects the *numbers* of segregating mutations but not their frequencies within the population (assuming that mutation biases remain constant over the relevant time period). The results are also consistent with compositional trend analyses (Shields *et al*. 1988; Akashi 1995; Vicario *et al*. 2007) that supported G- and/or C-ending major codons in all synonymous families in *D. melanogaster* (compositional trends are similar in *D. simulans*).

Genome-scale data allow us to refine the SFS analyses to a resolution of individual synonymous families. The following analyses will initially focus on 2-fold redundant sites and the term “synonymous” will refer to this class unless noted otherwise. SFS analyses for 4-fold redundant sites in CDS are presented in Supplementary Results and Discussion, Fixation biases in 4-fold synonymous families. In *D. simulans*, each of the 10 synonymous families (*i.e.*, Phe, Asp, Asn, His, Tyr, Ser_2_, Cys, Gln, Glu, and Lys) shows SFS differences similar to the general pattern of elevated within-population sample frequencies of GC-increasing (W→S) compared with AT-increasing (S→W) mutations (Fig. 2a; Table S6). The results are consistent with MCP predictions, but SFS for W→S mutations within short introns (SI) also indicate a GC fixation bias (Table S3; this table also shows data for long introns, LI, >100bp) as noted in previous studies (Kliman 2014; Jackson *et al*. 2017; Jackson and Charlesworth 2021; Yıldırım and Vogl 2024). GC fixation biases that are common to synonymous and intron mutations can include biased gene conversion and/or fitness differences unrelated to MCP.

**Fig. 2.**
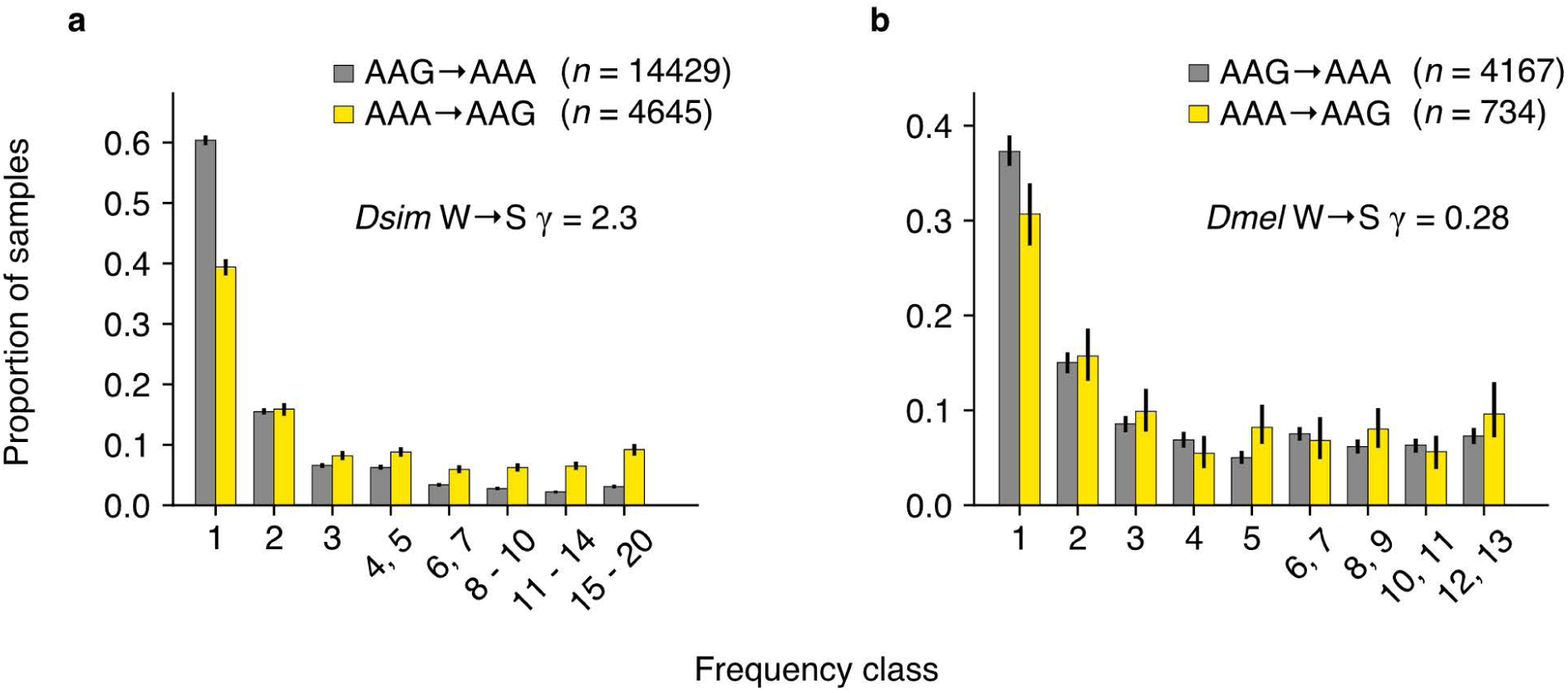
Site frequency spectra and fixation bias estimates: an example from Lysine codons in *D. simulans* and *D. melanogaster*. SFS for AAG→AAA (gray) is compared to that for AAA→AAG (yellow). (a) SFS for *D. simulans* (*Dsim*). (b) SFS for *D. melanogaster* (*Dmel*). See Tables S5 and S6 for the results of statistical tests for *Dsim* and *Dmel*, respectively. Counts (*n*) for the two polymorphism classes are shown (values are rounded to integers). Data from some frequency classes are pooled as indicated on the x-axis labels. “W→S γ” indicate maximum likelihood estimates of G- and C-favoring fixation biases using the approach of Glémin *et al*. (Glémin et al. 2015). SFS for intron GC-conservative changes are employed as a putatively neutral reference for the W→S γ estimation. Independent ancestral inference was performed for bootstrap replicates. Error bars indicate 95% CIs among 300 bootstrap replicates.

We employ summary statistics to capture the magnitude of differences in SFS comparisons in order to distinguish among underlying directional forces. Glémin *et al*. (Glémin *et al*. 2015) estimated a fixation-bias statistic (selection coefficient or conversion parameter scaled to population size) from SFS comparisons between forward and reverse mutations (see Materials and Methods). We employ “W→S γ” as a measure of the magnitude of SFS differences between W→S and S→W mutations. For some analyses, we employ a difference statistic (W→S aDAF skew, where aDAF refers to average derived allele frequency) as an alternative measure that does not require an assumed neutrally evolving control class of mutations. The two measures are strongly correlated (Fig. S1).

Directional forces favor G and/or C-ending codons across synonymous families in *D. simulans* (Fig. 3a) and the magnitude of GC fixation bias is heterogeneous among synonymous families. SI sequences also show GC fixation biases consistent with gBGC and/or natural selection. Larger W→S γ at synonymous sites than at SI sites supports additional directional forces favoring G and/or C-ending codons in coding regions (such as natural selection) as predicted under MCP. Kliman (Kliman 2014) found similar patterns among synonymous families as well as introns in polymorphism data from an autosome in *Drosophila pseudoobscura*.

**Fig. 3.**
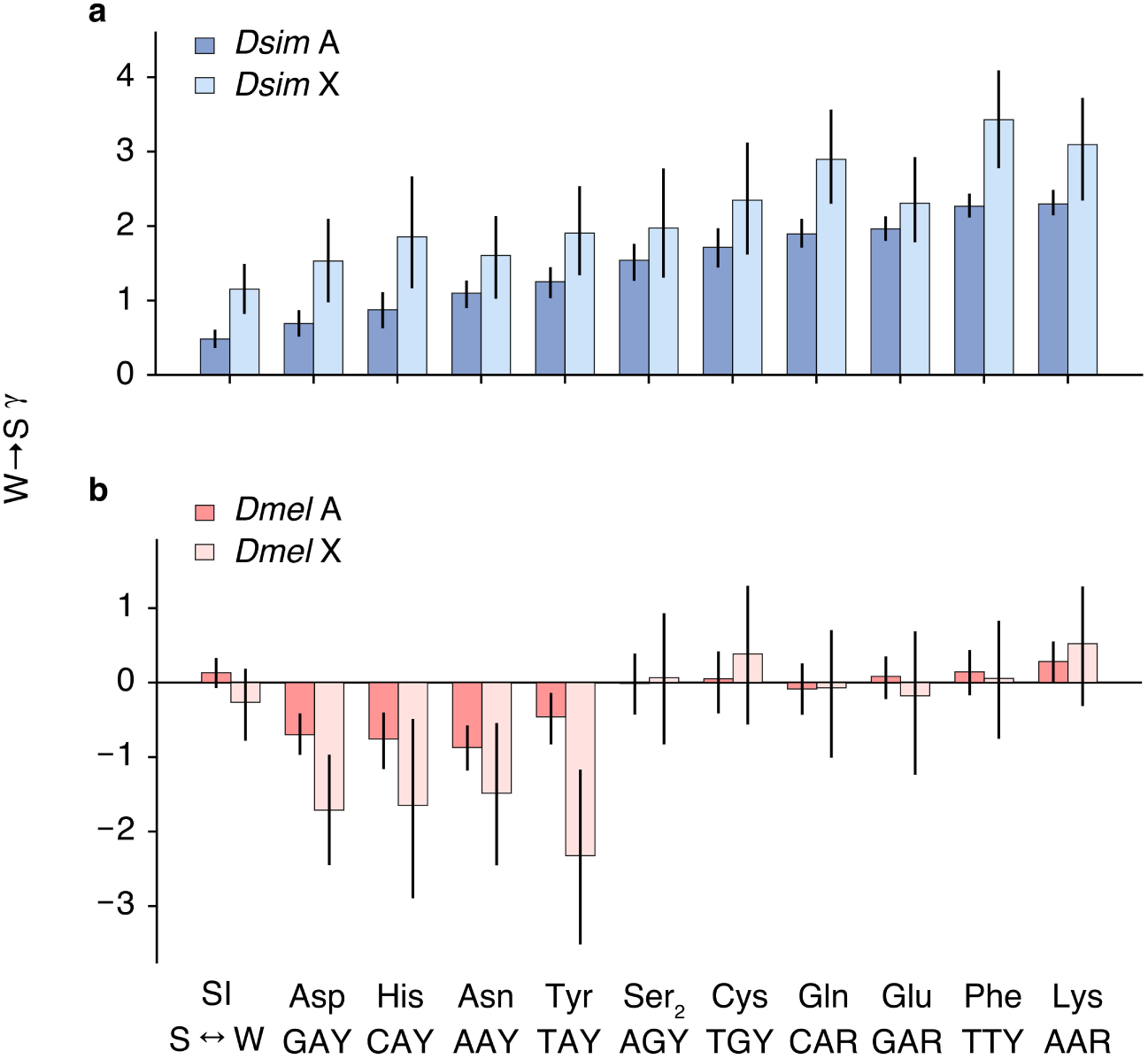
Synonymous family-specific fixation biases inferred from *D. simulans* and *D. melanogaster* site frequency spectra. Fixation biases for GC-altering changes, W→S γ, estimated using SFS data are shown for (a) *D. simulans* (*Dsim*) and (b) *D. melanogaster* (*Dmel*). Negative W→S γ values indicate fixation bias favoring AT. “SI” refers to short introns. Autosomal (darker colors; “A”) and X-linked (lighter colors; “X”) loci are analyzed separately. SFS for GC-conservative changes within short introns from autosomal and X-linked loci are employed as required neutral references for W→S γ estimation for the corresponding coding region data. Synonymous families are arranged in the order of γ values for *D. simulans* autosomal loci. W→S γ values are presented in Tables S5 and S6, for *D. simulans* and *D. melanogaster*, respectively. X-axis labels are shared between the top and bottom panels. Independent ancestral inference was performed for each bootstrap replicate. Error bars indicate 95% CIs among 300 bootstrap replicates.

The numbers of polymorphisms from genes on the X chromosome are sufficient for comparisons to autosomal loci (Table S6) and stronger fixation biases at X-linked loci are a conspicuous feature in *D. simulans* that holds across synonymous families (Fig. 3a; Wilcoxon signed-rank test *p* = 0.0051). Potential causes are difficult to resolve, however, and include greater efficacy of natural selection acting on partially recessive fitness effects (Charlesworth *et al*. 1987), GC elevation related to dosage compensation (Alekseyenko *et al*. 2012) or a greater impact of gBGC on the X chromosome.

In contrast to trends that generally conform to expectations under MCP in *D. simulans*, synonymous family-specific analyses reveal remarkable patterns in *D. melanogaster* polymorphism. Several synonymous families show SFS supporting fixation bias in the opposite direction to that observed in *D. simulans*: S→W mutations are segregating at higher frequencies than W→S mutations. The pattern holds for the four NAY synonymous families (codons encoding Asp, Asn, His, and Tyr) but is not found outside of these families (Fig. 3b; Table S7). The Lys synonymous family (AAR) shows SFS supporting GC fixation bias; W→S mutations segregate at higher frequencies than S→W mutations but the magnitude of fixation bias is small (Fig. 2b; Table S7). All other 2-fold synonymous families show indistinguishable SFS at both autosomal and X-linked loci (Table S7) consistent with previous analyses of pooled data (Akashi 1995).

*D. melanogaster* polymorphisms reveal differences in evolutionary processes for X-linked vs autosomal loci, an “X effect”, for NAY families. Fig. 3b shows consistent support, across the four NAY synonymous families, for stronger fixation biases favoring NAT over NAC at X-linked than at autosomal genes. Thus, *D. simulans* and *D. melanogaster* both show elevated fixation biases at X-linked genes, but in opposing directions, *i.e.*, favoring GC and AT, respectively. AT preference at NAY codons in *D. melanogaster* may reflect partially recessive fitness effects and/or other forces that override fixation biases toward GC (*i.e.*, gBGC or dosage compensation constraints). The cause(s) of X chromosome vs autosome differences in GC fixation biases in *D. simulans* are ambiguous as discussed above. However, the X effect for NAT fixation bias in *D. melanogaster* is difficult to explain in the absence of lineage-specific fitness benefits for NAT over NAC codons (interestingly, LI sites show an X effect in the opposite direction, GC preference. See Supplementary Results and Discussion, Fixation bias tests: mutation classes at intron sites). Mutation-driven scenarios require highly specified changes in mutational patterns (*i.e.*, specific to NAY codons, greater on the X chromosome, and timed near the MRCA of the species polymorphism) to explain NAY polymorphism patterns in *D. melanogaster*.

The analyses above support genome-wide differences in the direction of fixation biases at NAY codons in *D. melanogaster* and *D. simulans* but do not address the location of the preference shift within the evolutionary tree. Compositional trend analyses indicate NAC preference at all NAY families in species within the *melanogaster* group as well as in the more distantly related *obscura* group (Akashi and Schaeffer 1997; Vicario *et al*. 2007; see also below). SFS analyses further support NAC preference in *D. pseudoobscura* (Kliman 2014). The fixation bias patterns discussed above are most simply explained by a reversal of codon preference specific to NAY families in the *D. melanogaster* lineage after its split with *D. simulans*.

### Fixation biases in the ancestral lineages of D. simulans and D. melanogaster

The SFS analyses above focused on polymorphisms (*i.e.*, mutations segregating within populations) in *D. simulans* and *D. melanogaster* and revealed strong evidence for AT fixation biases specific to NAY codons in *D. melanogaster*. We also examined patterns deeper in the evolutionary histories of these species. Fixations on the *D. simulans* ancestral (*ms*-*s’*) and *D. melanogaster* ancestral (*ms*-*m’*) branches can reveal changes in mutation and/or fixation biases in the relatively short lineages prior to the *s’* and *m’* nodes (Fig. 1b). Previous analyses of pooled synonymous families showed weak AT content increases in the *D. simulans* ancestral lineage (Begun 2001; Kern and Begun 2005; Akashi *et al*. 2006; Singh *et al*. 2009; Jackson *et al*. 2017) and stronger AT content increases in the *D. melanogaster* ancestral lineage consistent with reduced GC fixation biases (Akashi 1995; Akashi 1996; Kern and Begun 2005; Akashi *et al*. 2006; Singh *et al*. 2009; Poh *et al*. 2012; Jackson *et al*. 2017).

We employ a summary statistic to capture the direction and extent of departures from GC content equilibrium: W→S fixation skew, (*a* − *b*) / (*a* + *b*), where *a* = *N*_W→S_, the number of W→S fixations and *b* = *N*_S→W_, the number of S→W fixations. Expected W→S fixation skew is zero at steady-state GC content and differs in sign, but is scaled symmetrically, for GC-increasing and GC-decreasing departures from equilibrium.

GC content decline is prevalent in the ancestral *D. simulans* lineage among mutation classes. However, W→S fixation skew is *positive* for our presumed best candidates for neutral evolution, SI with the lowest GC content (Fig. 4a; Fig. S4). We attribute GC elevation at putatively neutrally evolving sites to an increase in mutational pressure toward GC. From coding regions, we analyzed synonymous changes within NAY and “non-NAY” families (synonymous families encoding Phe, Cys, Ser_2_, Lys, Gln, and Glu) separately. For all three mutation classes (*i.e.*, SI, non-NAY and NAY), loss of GC intensifies with ancestral GC content (Fig. 4a; Fig. S6c). Such near-linear trends are consistent with simple MCP scenarios of non-stationary mutation ratio and/or fixation bias (Akashi *et al*. 2007). In particular, the *D. simulans* W→S fixation skew patterns appear roughly consistent with a combination of increase in GC mutation pressure and reduction in GC fixation bias (Fig. S6; note the contrasting patterns for LI, however).

**Fig. 4.**
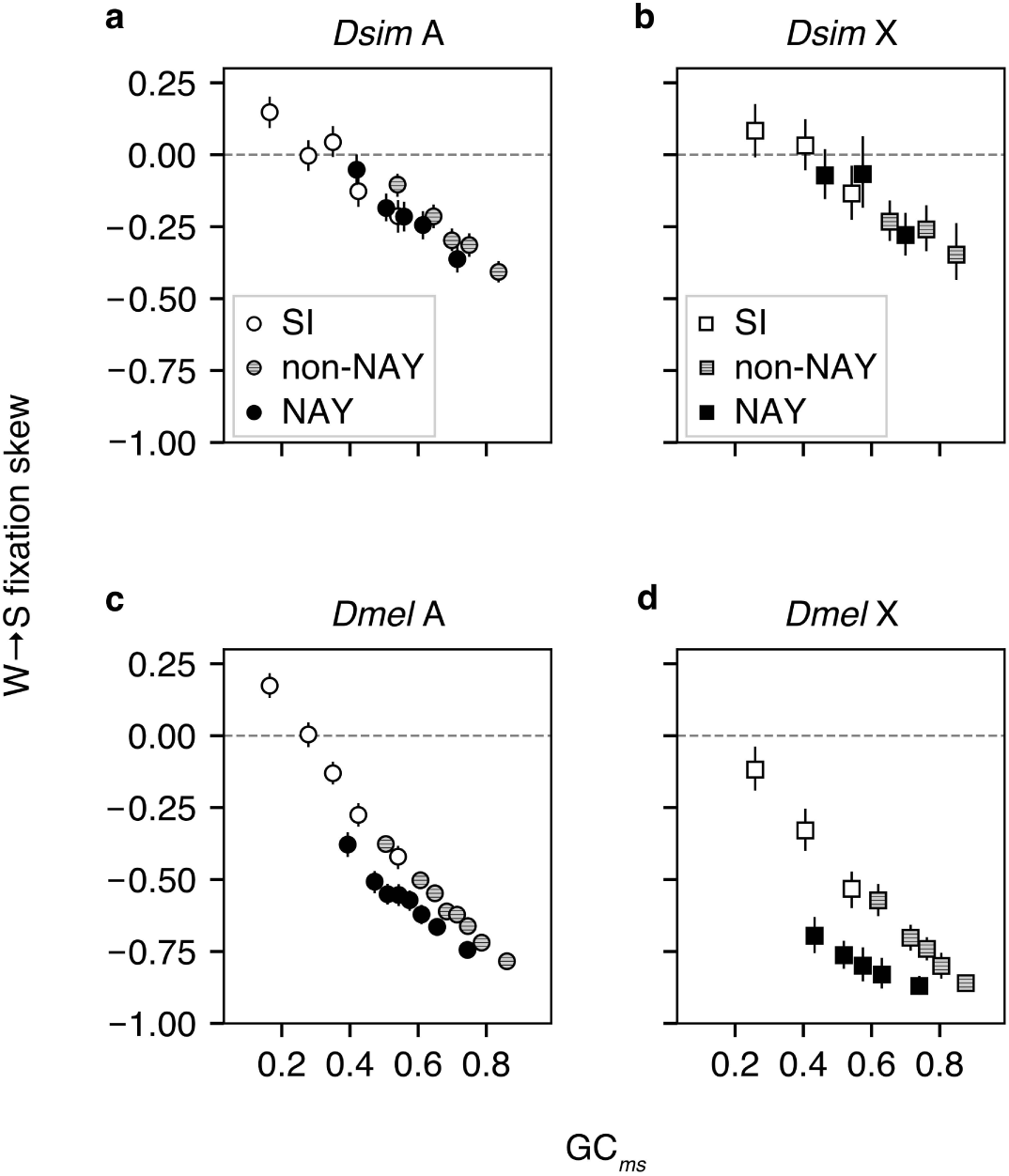
GC content changes in the *D. simulans* and *D. melanogaster* ancestral lineages. W→S fixation skew = (*N*_W_→_S_ − *N*_S_→_W_) / (*N*_W→S_ + *N*_S_→_W_), where *N*_W→S_ is W→S fixation count and *N*_S→W_ is S→W fixation count. This index measures the direction and magnitude of the departure from GC content equilibrium. *Changes* in GC content are plotted against GC content at *ms* node, GC*_ms_*. Fixations are inferred within internal branches: (a) and (b) *ms-s for D. simulans* fixation data and (c) and (d) *ms*-*m’* for *D. melanogaster* fixation data (Fig. 1b). (a) and (c) Autosomal and (b) and (d) X-linked loci are analyzed separately. The legends in (a) and (b) apply to (c) and (d), respectively. “SI” indicates short introns. Synonymous families are pooled into two classes: non-NAY (Gln, Glu, Lys, Phe, Cys, and Ser_2_) and NAY (Asp, Asn, His, and Tyr). For binning, introns or CDS are ranked by GC*_ms_* for the mutation class and assigned to bins with similar numbers of intron sites or codons, respectively. Statistical analyses are presented in Table S12. W→S fixation skews are compared between A vs X in Fig. S7. Error bars indicate 95% CIs among 1000 bootstrap replicates.

We tested whether GC content changes are consistent with shared parameter fluctuations among SI, non-NAY, and NAY mutation classes. Data were partitioned so that comparisons are conducted within similar ancestral GC-content ranges (Table S11). SI and NAY classes show indistinguishable W→S fixation skews in *D. simulans* (Table S11). Non-NAY codons show a weak signal toward less AT-biased fixation skews than SI and NAY codons (Table S11), but low fixation counts on the *ms-s’* branch (Fig. 1) prevent firm conclusions.

The ancestral lineage for *D. melanogaster* (Fig. 1) shows strong negative relationships between GC change and ancestral GC content for SI, non-NAY, and NAY classes. The slopes of these trends are steeper than in *D. simulans* (Fig. 4). Low-GC SI, our assumed neutrally evolving class, show an elevation of GC similar to the pattern in *D. simulans* (Fig. 4b) consistent with a shift in GC-elevating mutation bias that predates the *ms* node. Stronger negative W→S fixation skew slopes can be explained by larger fixation bias reductions (smaller *N_e_s*) in the ancestral *D. melanogaster* lineage (Akashi et al. 2007). Notably, NAY codons show accelerated AT content shifts compared to non-NAY codons within this lineage (Table S11). Extents of GC content change were statistically indistinguishable between SI sites and non-NAY codons (Table S11). Such patterns are predicted under a combination of fixation biases *favoring* S→W mutations at NAY codons *and* reduced GC fixation biases at SI sites and non-NAY codons.

X effects are a compelling feature in the SFS analyses (see above) and W→S fixation skew comparisons between X-linked vs. autosomal genes reveal similarly notable patterns. We focus on *D. melanogaster* because the low number of fixations on the *D. simulans* lineage limit the exploration of X effects (findings are discussed in Supplementary Results and Discussion). In *D. melanogaster*, non-NAY, NAY, and 4-fold synonymous classes as well as SI mutation classes show small, but statistically significant, elevations of AT gains at X-linked loci compared to autosomal loci (Fig. S7e-h and Table S12). Notably, NAY codons show significantly greater AT gains at X-linked, than at autosomal loci, than other mutation classes (Fig. S7h and Table S13). Enhanced efficacy of NAT codon preference at X-linked, relative to autosomal, loci for both polymorphism (post-*m’* node) and fixations (*ms*-*m’* nodes) in the *D. melanogaster* lineage provides a simple explanation for this pattern and is consistent with partially recessive fitness advantages for NAC to NAT mutations.

### tRNA modification: a potential factor in *D. melanogaster* preference reversals at NAY codons

Shared properties of NAY codons raise several possibilities for the biological cause(s) of the reversal of fitness benefits of NAT over NAC codons in the *D. melanogaster* lineage. These codons have common nucleotide composition at the 2nd and 3rd codon positions, show relatively weak fixation biases in *D. simulans* in SFS analyses (Fig. 3a), and are recognized by cognate tRNAs that undergo a particular chemical modification: queuosine (Q) modification at the anticodon 1st (wobble) position G nucleoside. Q modification is observed in the majority of eukaryotes (Harada and Nishimura 1972; White *et al*. 1973a; Kasai *et al*. 1975; Zallot *et al*. 2014) and is not known to occur for cognate tRNAs for other synonymous families (Fergus *et al*. 2015). Many eubacteria can synthesize the Q precursor, queine, but eukaryotes lack the required biochemical pathways and sequester the micronutrient from their diet and/or gut microbiota. Because queine levels are strongly dependent on diet in *Drosophila* (White *et al*. 1973b; Jacobson *et al*. 1981; Zaborske *et al*. 2014), lineage-specific modification levels (related to nutrient availability) have been proposed to explain codon preference differences (Chiari *et al*. 2010; Zaborske *et al*. 2014).

Fitness benefits of Q modification are clear from the retention of this pathway in a wide range of eukaryotes, but the functional basis of the benefit remains difficult to determine and may differ among taxa (reviewed in Fergus *et al*. 2015). Such benefits could lie outside of impacts on translational elongation/accuracy; for example, tRNA-modification affects rates of breakdown of isoacceptors into tRNA-derived small RNAs that function in gene regulation and stress response (Wang *et al*. 2018; Muthukumar *et al*. 2024). Experimental studies of protein synthesis support similar translation rates at NAC and NAT codons for unmodified cognate tRNAs and higher translational elongation rates at NAC codons for Q-modified tRNAs (Meier *et al*. 1985). Changes in Q modification can have measurable effects on translation rates at both cognate and non-cognate codons and tissue- and sex-specific effects in mice (Cirzi *et al*. 2023). Importantly, compositional trend analyses consistently support translational preference of NAC over NAT in organisms that employ Q modification [*Escherichia coli* (Kanaya *et al*. 1999), *Bacillus subtilis* (Kanaya *et al*. 1999), *Saccharomyces pombe* (Kanaya *et al*. 2001), and *Caenorhabditis elegans* (Stenico *et al*. 1994; Duret and Mouchiroud 1999)] as well as in those that lack the modification [*Saccharomyces cerevisiae* (Akashi 2003), *Candida albicans* (Lloyd and Sharp 1992) and *Arabidopsis thaliana* (Duret and Mouchiroud 1999; Wright *et al*. 2004)]. These patterns argue against a simple relationship between Q modification levels and major codon identity; in the limited number of established cases of loss of Q modification, translation selection appears to favor NAC codons.

We tested for signals for major codon reassignment (reversals in translational advantages) as a cause of NAT preference in *D. melanogaster*. Greater fitness benefits in highly translated genes are a hallmark feature of MCP (Ikemura 1981a; Ikemura 1981b). We expect elevated fixation biases for minor to major codon mutations among genes that show the strongest patterns of ancestral MCP. In *D. simulans*, both NAY and non-NAY codons show greater GC fixation biases in genes with higher major codon usage at the *ms* node (Fig. S20). If NAT preference in *D. melanogaster* is driven by translational selection, then we expect similarly larger absolute magnitudes of fixation bias in genes under stronger translational selection. We detected no such trend; magnitudes of fixation bias are indistinguishable between high and low major codon usage genes (Fig. S20). We consider potential advantages of NAT codon usage that fall outside of the standard MCP model.

### Testing ApT vs ApC dinucleotide preference

Context-dependent fixation biases can arise from dinucleotide preferences that act in opposition to MCP-related GC fixation biases. ApT vs ApC dinucleotide preferences could, in principle, account for NAY family-specific polymorphism and fixation patterns and may show less association with gene expression level than MCP. Interactions between dinucleotide preferences and translational selection have been proposed in previous studies (Nussinov 1981; Antezana and Kreitman 1999; Kokate et al. 2021). We take advantage of features within the genetic code to distinguish specific codon and dinucleotide preference scenarios. Leu is encoded by six codons and, T→C transitions at the first position are synonymous among YTR codons. ApT dinucleotide preference predicts context-dependent fixation biases toward TTR codons for Leu codons downstream of a 5’ A (A|TTR vs A|CTR where pipe symbols indicate codon boundaries). We discuss *D. melanogaster* patterns before comparisons to *D. simulans* in order to assess causes of NAT preference. Fig. 5a confirms the predicted signal within *D. melanogaster* polymorphism; SFS is skewed toward elevated frequencies for C→T compared to T→C transitions among A|TTR↔A|CTR synonymous polymorphism. The magnitude of the fixation bias (aDAF skew is strongly correlated with γ; Fig. S1) appears roughly similar to NAT preference at NAY codons. This pattern is consistent with ApT over ApC dinucleotide preference but could also simply reflect translational selection favoring TTR over CTR Leu codons. We distinguish these possibilities by examining context (5’ nearest neighbor) effects for synonymous changes at Leu first codon positions within *D. melanogaster* polymorphism. Differences in SFS for T→C and reverse changes are specific to the 5’ A context; B|TTR↔B|CTR synonymous polymorphism show similar aDAF (ambiguity character B denotes C, T, or G nucleotides; Fig. 5a). Permutation tests support context-dependent T→C fixation bias; W→S aDAF skew is lower for A|TTR→A|CTR than for B|TTR→B|CTR (*p* < 0.005, 10^5^ iterations). These patterns together support ApT over ApC dinucleotide preference and, because Leu cognate tRNAs do not undergo queuosine modification, call into question the tRNA modification driven codon preference reversal scenario.

**Fig. 5.**
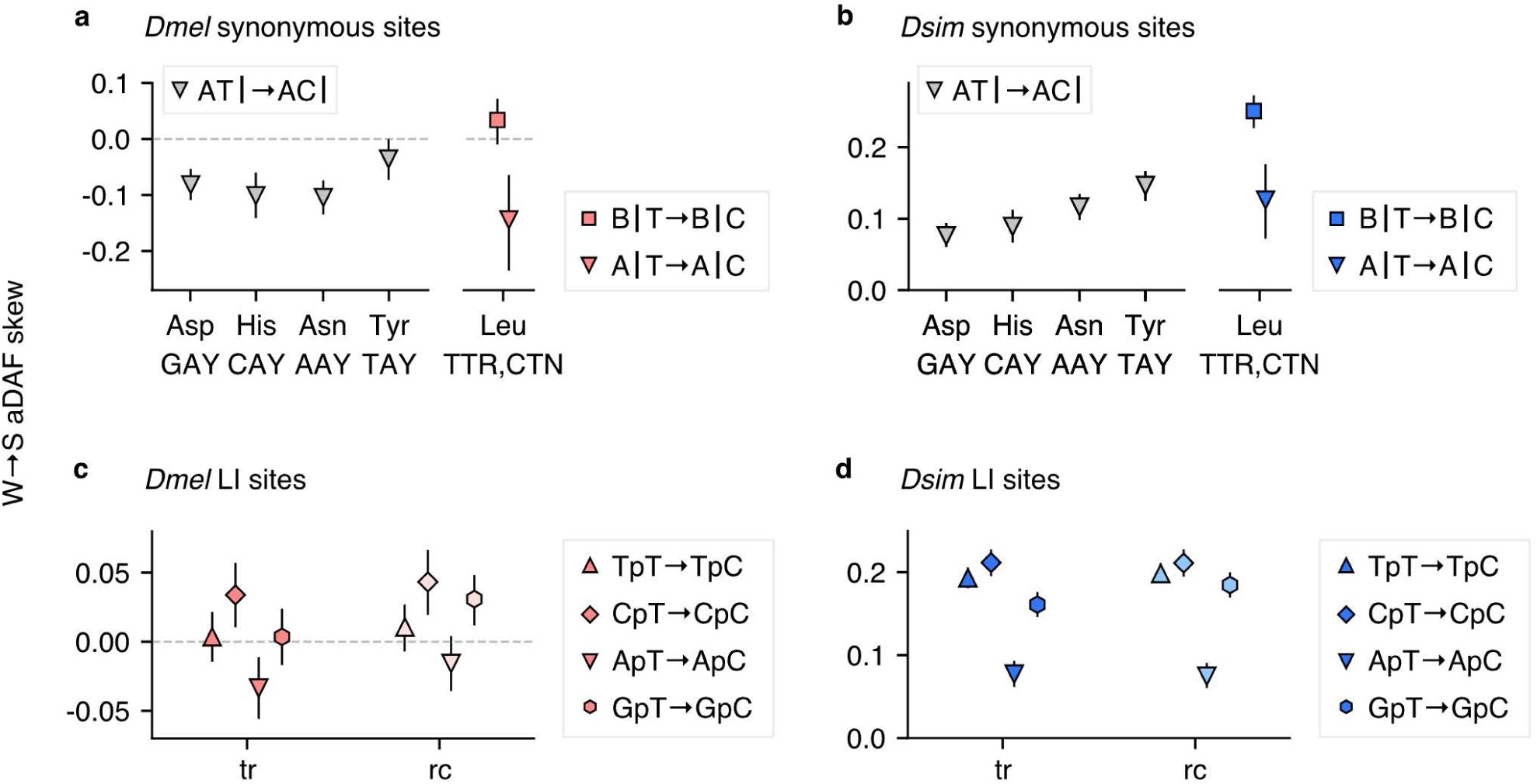
ApT vs ApC dinucleotide preference inferred from *D. melanogaster* and *D. simulans* site frequency spectra. W→S aDAF skew measures the direction and magnitude of SFS difference between “forward” and “reverse” context-dependent nucleotide changes. Arrows specify the forward direction. Positive W→S aDAF skew indicates higher frequencies for forward changes than reverse changes. W→S aDAF skews for ApT→ApC are compared to those for BpT→BpC (“B” indicates “non-A” or T, C, or G nucleotides). ApT→ApC and BpT→BpC changes at synonymous sites in (a) *D. melanogaster* (*Dmel*) and (b) *D. simulans* (*Dsim*). Pipes (“|”) indicate codon borders. ApT→ApC and other context-dependent nucleotide changes at long intron (LI) sites in (c) *Dmel* and (d) *Dsim*. Context-dependent nucleotide changes on the transcribed (tr) strand are indicated in the in-figure legends. Light colors indicate reverse complement (rc) changes (*e.g.*, ApA→GpA is the reverse complement of TpT→TpC). Error bars indicate 95% CIs among 1000 bootstrap replicates. Ancestral reconstructions are resampled in units of introns or CDS.

We examined first codon position synonymous changes at Leu codons to test for ApT vs ApC preference within *D. simulans* polymorphism. Fig. 5b illustrates context-dependent SFS differences among T→C and reverse synonymous changes at Leu first positions. SFS differences indicate a relatively strong GC fixation bias among B|TTR↔B|CTR synonymous polymorphism. However, the same nucleotide transition in a 5’ A context shows more similar SFS; permutation tests support lower W→S aDAF skew for A|TTR→A|CTR than for B|TTR→B|CTR (*p* < 0.0001, 10^5^ iterations). Viewed in isolation, this difference could be interpreted most simply as support for context-dependent weakening of first codon position C preference (*i.e.*, reduced MCP) at Leu codons. However, support for fixation biases favoring A|TTR over A|CTR in *D. melanogaster* (Fig. 5a) lead us to posit that this reduction in the magnitude of aDAF skew in *D. simulans* (Fig. 5b) reflects a common ApT over ApC preference acting in opposition to a stronger CTR-favoring force (presumably MCP) than in *D. melanogaster*. We employ similar reasoning in the analyses below.

We tested for dinucleotide preferences for ApT vs GpT, the reverse complement of ApT vs ApC within coding regions. We consider synonymous A↔G transitions in the context of 3’ neighboring T vs V (C, A, or G nucleotides). Overall trends toward aDAF skew differences expected under ApT over GpT preference are strong among 13 synonymous families (*D. melanogaster*, *p* < 0.0008; *D. simulans*, *p* < 0.0003; Fig. S8; Tables S14 and S15). We propose a scenario of MCP-related GC pressure balanced by an opposing ApT over ApC pressure at NAY and Leu codons in both *D. melanogaster* and *D. simulans*. Synonymous changes within a larger number of families support this dinucleotide preference on the non-transcribed strand. Reduced efficacy of MCP reveals the action of this dinucleotide preference at *D. melanogaster* NAY codons but nearest-neighbor dependent SFS in other synonymous families reveal similar forces, acting with different relative intensities, in *D. simulans*.

We expanded the data for analyses of ApT vs ApC preference by including polymorphisms within LI. Although constraints within LI are not well understood (Halligan *et al*. 2004), larger numbers of segregating sites in these data (Tables S16 and S17) allow us to examine fixation biases in specific contexts. In *D. melanogaster*, aDAF skew indicates T over C preference in a 5’ A context which is absent, or acting in the opposing direction, for other 5’ contexts (Fig. 5c; Table S16). Confidence intervals are large in the *D. melanogaster* data, but context-dependent SFS differences appear to be similar on transcribed and non-transcribed strands (Fig. 5c). Higher polymorphism levels in the *D. simulans* data help to reveal strong context-dependence in T→C fixation biases. BpT→BpC mutations are strongly favored but this W→S fixation bias is clearly reduced for ApT→ApC changes (Fig. 5d). These contrasting patterns of SFS are similar for reverse complement changes. SFS analyses of polymorphisms within LI support ApT over ApC dinucleotide preference in non-coding regions and confirm that evolutionary forces act similarly in the transcribed and non-transcribed strands.

These features of ApT over ApC preference may be important in considering the biological function(s) underlying dinucleotide preferences. Dinucleotides can impact biophysical properties of DNA such as bendability that can affect chromatin structure, DNA repair, and protein binding (Antezana and Kreitman 1999; reviewed in Sievers *et al*. 2023). Highly ApT-rich regions can be prone to chromosomal mutations (Irony-Tur Sinai *et al*. 2019). Dinucleotides can influence secondary structure formation in mRNAs (Workman and Krogh 1999) which can affect the translation process (Yang *et al*. 2014). However, the similarity of preferences in coding and non-coding regions and on transcribed and non-transcribed strands is more consistent with DNA constraints. Langley and coworkers (Langley *et al*. 2014) found enrichment of ApA, TpT, and GpC dinucleotides in *D. melanogaster* intergenic and intron regions. These dinucleotides occur in (roughly 10bp) periodic patterns consistent with nucleosome positioning constraints. Population genetic analyses support fixation biases underlying these patterns. However, ApT dinucleotides are not over-represented in their analyses (Langley *et al*. 2014) or in previous compositional analyses within coding regions (Antezana and Kreitman 1999; Gentles and Karlin 2001; but see Fedorov *et al*. 2002). The functional bases for ApT over ApC dinucleotide preferences remain unclear.

### Genome dynamics under near neutrality

Sensitivity of rates of evolution and levels of adaptation to effective population size are central features of nearly neutral evolution. *N_e_* variation on the time-scale of evolutionary processes (and corresponding modulations in selection intensity, *N_e_s*) are often invoked to explain patterns in genome variation (Ohta 1993; Moran 1996; Akashi et al. 2006). Our findings challenge whether *N_e_* fluctuations are sufficient to account for evolutionary dynamics across mutation classes in either the *D. melanogaster* or the *D. simulans* lineage.

In *D. melanogaster*, reduced historical *N_e_* could explain patterns of declining GC usage among classes of synonymous sites in coding regions (non-NAY and 4-fold synonymous families), SI, and LI (Fig. S6d). However, the same factor should also have attenuated the efficacy of selection for preferred dinucleotides. Changes in the relative fitness effects (*i.e*., an increase in selection coefficients favoring ApT vs ApC dinucleotides relative to those favoring major codons and intron GC-elevation) appears necessary to account for our observations.

In *D. simulans*, proportionate changes in *N_e_* are consistent with declines in GC among several classes of synonymous change (4-fold and NAY synonymous families; Fig. S6c) as well as SI. However, LI show a distinct pattern of GC content stability across GC classes in the ancestral lineage; this suggests an elevation in selection coefficients favoring GC at LI sites relative to similar forces at SI and synonymous sites. Our findings are difficult to explain in the absence of changes in the relative magnitudes of fitness effects among mutation classes in both focal evolutionary lineages. More generally, our findings indicate that pleiotropy plays an important role in synonymous site and non-coding DNA evolution and that small fitness effects may not reflect weak functional effects; in particular, smaller fixation biases at NAY codons compared to other synonymous families do not necessarily indicate smaller translational effects of synonymous changes in these families.

Our study demonstrates how increasing the resolution (including codon-family specificity and short time scales) of molecular evolutionary studies can reveal context effects and interactions among opposing forces in genome adaptation. Between-species contrasts of polymorphism and fixation patterns were critical for identifying dinucleotide preferences that we hope will motivate functional genomic analyses.

## Materials and Methods

Abbreviations are summarized in Table S1.

### DNA sequences

We employed available genome data for isofemale lines established from natural populations of *D. melanogaster* and *D. simulans*. Data from a Rwanda population of *D. melanogaster* (Pool *et al*. 2012) was downloaded from http://www.dpgp.org/dpgp2/DPGP2.html (last downloaded on 22nd July 2014). Sequence analyses support a relatively stable effective population size (Sprengelmeyer *et al*. 2020) and lower inversion frequencies (Lack *et al*. 2015; Sprengelmeyer *et al*. 2020) for this population compared to other populations for which population genomic data are available. Among 22 lines reported in the Pool *et al*. study, 14 lines were employed for this analysis (RG2, RG3, RG5, RG9, RG18N, RG19, RG22, RG24, RG25, RG28, RG32N, RG34, RG36 and RG38N).We excluded five lines that show evidence for a high proportion of admixture with European populations [RG10, RG11N, RG15, RG21N and RG35; (Pool *et al*. 2012)]. In addition, we excluded three lines that contain relatively high proportions of ambiguous nucleotides (RG4N, RG7 and RG33). We extracted predicted CDS and intron sequences from the genomic sequences for a total of 13,691 protein-coding genes using FlyBase r6.24 *D. melanogaster* genome annotations (ftp://ftp.flybase.net/genomes/Drosophila_melanogaster/dmel_r6.24_FB2018_05/; last downloaded on 12th July 2019; Supplementary Methods).

We reconstructed genome sequences for *D. simulans* within-species samples using available data (https://drive.google.com/drive/folders/0B4O-acc8EJwheS1HZ1hnWkpOQlE; last downloaded on 1st March 2017; Supplementary Methods). We analyzed the DNA variant information for 21 lines from a Madagascar population (Rogers *et al*. 2014; Jackson *et al*. 2017); 10 lines (MD06, MD105, MD106, MD15, MD199, MD221, MD233, MD251, MD63, and MD73) were sequenced by Rogers *et al*. (Rogers *et al*. 2014), and 11 lines (MD03, MD146, MD197, MD201, MD224, MD225, MD235, MD238, MD243, MD255, and MD72) were sequenced by Jackson *et al*. (Jackson *et al*. 2017). These sequences show high nucleotide diversity relative to data from other population samples (Dean and Ballard 2004). These 10 and 11 lines were sampled from the same localities in Madagascar by Rogers *et al*. (Rogers *et al*. 2014) and William Ballard, respectively, as described in Jackson *et al*. (Jackson *et al*. 2017). We extracted predicted CDS and intron sequences from the genome sequences for a total of 13,831 protein-coding genes using FlyBase r2.02 *D. simulans* genome annotations (ftp://ftp.flybase.net/genomes/Drosophila_simulans/dsim_r2.02_FB2017_04; last downloaded on 9th March 2020).

Genome sequences from species within the *D. melanogaster* subgroup, *D. yakuba* and *D. erecta* were employed as outgroup data. We extracted predicted CDS and intron sequences from the genome sequences using genome annotations for FlyBase r1.05 *D. yakuba* (ftp://ftp.flybase.net/genomes/Drosophila_yakuba/dyak_r1.05_FB2016_05/; last downloaded on 12th July 2019) and FlyBase r1.05 *D. erecta* (ftp://ftp.flybase.net/genomes/Drosophila_erecta/dere_r1.05_FB2016_05/; last downloaded on 12th July 2019).

We constructed CDS and intron sequence alignments including *D. melanogaster* and *D. simulans* within-species samples and outgroup sequences (see Supplementary Methods). Custom Python codes used in the alignment pipeline are available at https://github.com/nigevogen. The data set includes 10,122 CDS and 18,719 intron alignments for autosomal loci and 1,746 CDS and 2,705 intron alignments for X-linked loci which are available at https://zenodo.org/communities/evogen-akashi-lab/.

### Inference of polymorphic changes

We define “synonymous family” as a group of synonymous codons that can be interchanged in single nucleotide steps; serine coding codons were split into a 2-fold (AGY, referred to as Ser_2_) and a 4-fold (TCN, referred to as Ser_4_) family. We analyzed 10 synonymous families (Phe, Asp, Asn, His, Tyr, Ser_2_, Cys, Gln, Glu, and Lys) with 2-fold redundancy and six families (Ala, Gly, Val, Thr, Pro and Ser_4_) with 4-fold redundancy.

We estimated probabilities of nucleotides at ancestral nodes in the gene tree shown in Fig. 1b. Although the sequences examined are relatively closely related, ancestral inference under simple substitution models such as maximum parsimony can be unreliable when character states are biased and/or composition is changing within the gene tree (Collins *et al*. 1994; Matsumoto and Akashi 2018). In addition, our analyses require inference of ancestral and derived states at segregating sites in recombining regions where gene trees may differ among sites. In order to address these issues, we employed a likelihood-based approach that incorporates uncertainty in ancestral inference. We inferred ancestral nucleotides for both within- and between-species variation using a combination of BASEML (Yang 2007) and the Bifurcating Tree with Weighting (BTW) approach (Matsumoto and Akashi 2018). For BASEML ancestral inference, we employed the GTR-NH_b_ nucleotide substitution model (Tavaré 1986; Matsumoto *et al*. 2015) with a newly implemented option (available on BASEML in PAMLver4.9) that allows user-defined branches to share transition parameters. Here, we set parameters to be shared within (but not between) collapsed sequence pairs in *D. melanogaster* and *D. simulans* (terminal branches leading to *m_c1_*, *m_c2_* and *s_c1_*, *s_c2_*, respectively, in Fig. 1b). Ancestral inference was conducted separately for data from autosomal and X-linked loci. We refer to inferred changes on the *m’*-*m_c1_* and *m’*-*m_c2_* branches as polymorphisms in *D. melanogaster* and those on the *s’*-*s_c1_* and *s’*-*s_c2_* branches as polymorphisms in *D. simulans*. We treated probabilities of changes as counts for the numbers of polymorphic mutations for each of 12 mutation classes (Supplementary Methods). Ancestral inference for context-dependence analysis is also described in Supplementary Methods.

### Polymorphism analysis

We analyzed SFS for forward and reverse mutations between pairs of nucleotides (*e.g.*, A→G vs G→A). We use the term “forward” mutations for W→S changes (where W indicates A or T, and S indicates G or C) among GC-altering mutations and for T→A and C→G among GC-conservative mutations. Mutations in the opposite direction are termed “reverse” mutations (designations of the terms “forward” and “reverse” are arbitrary). Among the six possible pairs, four are GC-altering (*i.e.*, W↔S) and two are GC-conservative (*i.e.*, A↔T and G↔C). Mann-Whitney U (MWU) tests were employed to test for SFS differences among synonymous and intron mutations. Direct SFS comparisons between mutation classes that are physically interspersed within DNA (Bulmer 1971; Sawyer *et al*. 1987) attempt to control for effects of linked selection and demographic history in the inference of fixation biases. This approach can be employed to test weak selection models of synonymous codon usage bias (Akashi and Schaeffer 1997; Akashi 1999).

We estimated the magnitude of fixation biases using a maximum likelihood method (Glémin *et al*. 2015) that fits observed SFS to theoretical expectations (Fisher 1930; Wright 1938). SFS of putatively neutral mutations (here, intron GC-conservative mutations; see also Table S3) were employed to adjust for possible departures from steady-state SFS caused by demographic history and linked selection (Eyre-Walker *et al*. 2006). We employed the M1 model in the anavar software package (Muyle *et al*. 2011; Glémin *et al*. 2015) to estimate W→S γ, the fixation bias statistic. Positive and negative values of W→S γ indicate fixation biases that elevate and reduce GC content, respectively. This statistic is an estimate of the product of 4*N_e_* and either the selection coefficient (*s*), or the intensity of the conversion bias (*b*) in selection and biased gene conversion models, respectively. In our analyses, γ estimates are strongly correlated with simple difference statistics between average derived allele frequencies (aDAF skew; Fig. S1). We employ these indices as summary statistics for the magnitude of difference between SFS in comparisons across mutation classes and X-linked vs autosomal loci.

### Inference of fixations in coding regions

We inferred synonymous and replacement changes within the *D. simulans* ancestral (*ms-s’*) and *D. melanogaster* ancestral (*ms-m’*) lineages by combining an approach introduced in Akashi *et al*. (Akashi *et al*. 2006) with the method described above (BTW using the GTR-NH_b_ model). This approach reconstructs internal node codon configurations (INCC) by combining separate estimations of internal node nucleotide configurations (INNC) for three codon positions, assuming independent processes among positions (Akashi *et al*. 2006).

We employed the alignments described in Supplementary Methods. We inferred ancestral codons at internal nodes in the tree shown in Fig. 1b. A pair of ancestral and derived codons that differ at a single position will be referred to as a “codon change” (*e.g*., AAA at the *ms* node and AAG at the *m’* node at a given codon position in the aligned CDS is an AAA→AAG codon change in the *ms*-*m’* branch). The probability of INCC is used as a “count” of the change. For the case of codon pairs that differ at two or three positions, we calculated the relative probability for each possible minimal step path from ancestral and derived codon. These probabilities were weighted by the numbers of synonymous and nonsynonymous changes involved in a path (Nei and Gojobori 1986) using the nonsynonymous-to-synonymous substitution rate ratio calculated from codon pairs that differ at a single position (*d*_N_ / *d*_S_ = 0.1234 for autosomal loci; *d*_N_ / *d*_S_ = 0.1434 for X-linked loci). We used the Nei and Gojobori (Nei and Gojobori 1986) method to count the numbers of synonymous and nonsynonymous sites.

In our analyses, “fixations” refer to changes inferred on the *ms*-*s’* and *ms*-*m’* branches and are thus “fixed in the sample” and may include mutations segregating at high frequencies within the populations (these changes do not overlap with data for the SFS analysis). We also note that our approach of fixation counting does not consider multiple changes at a site within a branch. This can cause underestimation of fixation counts, but the effect is expected to be minor given the low divergence levels between *D. simulans* and *D. melanogaster* (Matsumoto *et al*. 2015; Matsumoto and Akashi 2018).

### W→S fixation skew analysis

To infer changes in fixation bias and/or mutation rate bias, we compared *d*_WS,SW_ among genes that differ in GC content at the *ms* node (GC*_ms_*). GC*_ms_* is the sum of probabilities for INNC with G or C nucleotides at the *ms* node divided by the number of “sites” (*i.e.*, the total probabilities for INNC) for introns and serves as a proxy for ancestral GC fixation bias. We used a similar approach to calculate GC*_ms_* at synonymous sites, the proportion of G- and/or C-ending codons at the *ms* node. GC*_ms_* are used for binning introns and CDS into non-overlapping bins with similar numbers of sites (nucleotide positions for intron and codons for CDS). We filtered introns and CDS that include fewer than 10 aligned sites. Ancestral reconstructions were resampled in units of intron and CDS. Statistical approaches to detect differences in W→S fixation skews between mutation classes are described in Supplementary Results and Discussion, GC content evolution in the *D. simulans* and *D. melanogaster* ancestral lineages.

## References

Akashi H. 1995. Inferring weak selection from patterns of polymorphism and divergence at “silent” sites in Drosophila DNA. Genetics. 139(2):1067–1076.

Akashi H. 1996. Molecular evolution between *Drosophila melanogaster* and *D. simulans*: reduced codon bias, faster rates of amino acid substitution, and larger proteins in *D. melanogaster*. Genetics. 144(3):1297–1307.

Akashi H. 1999. Inferring the fitness effects of DNA mutations from polymorphism and divergence data: statistical power to detect directional selection under stationarity and free recombination. Genetics. 151(1):221–238.

Akashi H. 2001. Gene expression and molecular evolution. Curr Opin Genet Dev. 11(6):660–666. doi:10.1016/s0959-437x(00)00250-1.

Akashi H. 2003. Translational selection and yeast proteome evolution. Genetics. 164(4):1291–1303.

Akashi H, Goel P, John A. 2007. Ancestral inference and the study of codon bias evolution: implications for molecular evolutionary analyses of the *Drosophila melanogaster* subgroup. Fay J, editor. PloS One. 2(10):e1065. doi:10.1371/journal.pone.0001065.

Akashi H, Ko W-Y, Piao S, John A, Goel P, Lin C-F, Vitins AP. 2006. Molecular evolution in the *Drosophila melanogaster* species subgroup: frequent parameter fluctuations on the timescale of molecular divergence. Genetics. 172(3):1711–1726. doi:10.1534/genetics.105.049676.

Akashi H, Schaeffer SW. 1997. Natural selection and the frequency distributions of “silent” DNA polymorphism in Drosophila. Genetics. 146(1):295–307.

Alekseyenko AA, Ho JWK, Peng S, Gelbart M, Tolstorukov MY, Plachetka A, Kharchenko PV, Jung YL, Gorchakov AA, Larschan E, et al. 2012. Sequence-specific targeting of dosage compensation in Drosophila favors an active chromatin context. PLoS Genet. 8(4):e1002646. doi:10.1371/journal.pgen.1002646.

Andersson GE, Kurland CG. 1991. An extreme codon preference strategy: codon reassignment. Mol Biol Evol. 8(4):530–544. doi:10.1093/oxfordjournals.molbev.a040666.

Antezana MA, Kreitman M. 1999. The Nonrandom Location of Synonymous Codons Suggests That Reading Frame-Independent Forces Have Patterned Codon Preferences. J Mol Evol. 49(1):36–43. doi:10.1007/pl00006532.

Begun DJ. 2001. The frequency distribution of nucleotide variation in *Drosophila simulans*. Mol Biol Evol. 18(7):1343–1352. doi:10.1093/oxfordjournals.molbev.a003918.

Bulmer M. 1991. The selection-mutation-drift theory of synonymous codon usage. Genetics. 129(3):897–907.

Bulmer MG. 1971. Protein polymorphism. Nature. 234(5329):410–411. doi:10.1038/234410b0.

Charlesworth B, Coyne JA, Barton NH. 1987. The Relative Rates of Evolution of Sex Chromosomes and Autosomes. Am Nat. 130(1):113–146. doi:10.1086/284701.

Chiari Y, Dion K, Colborn J, Parmakelis A, Powell JR. 2010. On the possible role of tRNA base modifications in the evolution of codon usage: queuosine and Drosophila. J Mol Evol. 70(4):339–345. doi:10.1007/s00239-010-9329-z.

Cirzi C, Dyckow J, Legrand C, Schott J, Guo W, Perez Hernandez D, Hisaoka M, Parlato R, Pitzer C, van der Hoeven F, et al. 2023. Queuosine-tRNA promotes sex-dependent learning and memory formation by maintaining codon-biased translation elongation speed. EMBO J. 42(19):e112507. doi:10.15252/embj.2022112507.

Collins TM, Wimberger PH, Naylor GJP. 1994. Compositional Bias, Character-State Bias, and Character-State Reconstruction Using Parsimony. Syst Biol. 43(4):482–496. doi:10.1093/sysbio/43.4.482.

Curran JF, Yarus M. 1989. Rates of aminoacyl-tRNA selection at 29 sense codons *in vivo*. J Mol Biol. 209(1):65–77. doi:10.1016/0022-2836(89)90170-8.

Dana A, Tuller T. 2014. The effect of tRNA levels on decoding times of mRNA codons. Nucleic Acids Res. 42(14):9171–9181. doi:10.1093/nar/gku646.

Dean MD, Ballard JWO. 2004. Linking phylogenetics with population genetics to reconstruct the geographic origin of a species. Mol Phylogenet Evol. 32(3):998–1009. doi:10.1016/j.ympev.2004.03.013.

Drosophila 12 Genomes Consortium. 2007. Evolution of genes and genomes on the Drosophila phylogeny. Nature. 450(7167):203–218. doi:10.1038/nature06341.

Duret L. 2002. Evolution of synonymous codon usage in metazoans. Curr Opin Genet Dev. 12(6):640–649. doi:10.1016/s0959-437x(02)00353-2.

Duret L, Galtier N. 2009. Biased gene conversion and the evolution of mammalian genomic landscapes. Annu Rev Genomics Hum Genet. 10:285–311. doi:10.1146/annurev-genom-082908-150001.

Duret L, Mouchiroud D. 1999. Expression pattern and, surprisingly, gene length shape codon usage in *Caenorhabditis*, *Drosophila*, and *Arabidopsis*. Proc Natl Acad Sci U S A. 96(8):4482–4487. doi:10.1073/pnas.96.8.4482.

Eyre-Walker A, Woolfit M, Phelps T. 2006. The distribution of fitness effects of new deleterious amino acid mutations in humans. Genetics. 173(2):891–900. doi:10.1534/genetics.106.057570.

Fedorov A, Saxonov S, Gilbert W. 2002. Regularities of context-dependent codon bias in eukaryotic genes. Nucleic Acids Res. 30(5):1192–1197. doi:10.1093/nar/30.5.1192.

Fergus C, Barnes D, Alqasem MA, Kelly VP. 2015. The queuine micronutrient: charting a course from microbe to man. Nutrients. 7(4):2897–2929. doi:10.3390/nu7042897.

Fisher RA. 1930. The Genetical Theory of Natural Selection. Clarendon Press, Oxford. https://www.biodiversitylibrary.org/item/27468.

Gentles AJ, Karlin S. 2001. Genome-scale compositional comparisons in eukaryotes. Genome Res. 11(4):540–546. doi:10.1101/gr.163101.

Glémin S, Arndt PF, Messer PW, Petrov D, Galtier N, Duret L. 2015. Quantification of GC-biased gene conversion in the human genome. Genome Res. 25(8):1215–1228. doi:10.1101/gr.185488.114.

Grantham R, Gautier C, Gouy M, Jacobzone M, Mercier R. 1981. Codon catalog usage is a genome strategy modulated for gene expressivity. Nucleic Acids Res. 9(1):r43–74. doi:10.1093/nar/9.1.213-b.

Halligan DL, Eyre-Walker A, Andolfatto P, Keightley PD. 2004. Patterns of Evolutionary Constraints in Intronic and Intergenic DNA of Drosophila. Genome Res. 14(2):273–279. doi:10.1101/gr.1329204.

Harada F, Nishimura S. 1972. Possible anticodon sequences of tRNAHis, tRNAAsn, and tRNAAsp from *Escherichia coli* B. Universal presence of nucleoside Q in the first position of the anticodons of these transfer ribonucleic acids. Biochemistry. 11(2):301–308. doi:10.1021/bi00752a024.

Ikemura T. 1981a. Correlation between the abundance of *Escherichia coli* transfer RNAs and the occurrence of the respective codons in its protein genes: A proposal for a synonymous codon choice that is optimal for the *E. coli* translational system. J Mol Biol. 151(3):389–409. doi:10.1016/0022-2836(81)90003-6.

Ikemura T. 1981b. Correlation between the abundance of *Escherichia coli* transfer RNAs and the occurrence of the respective codons in its protein genes. J Mol Biol. 146(1):1–21. doi:10.1016/0022-2836(81)90363-6.

Ikemura T. 1982. Correlation between the abundance of yeast transfer RNAs and the occurrence of the respective codons in protein genes: Differences in synonymous codon choice patterns of yeast and *Escherichia coli* with reference to the abundance of isoaccepting transfer RNAs. J Mol Biol. 158(4):573–597. doi:10.1016/0022-2836(82)90250-9.

Ikemura T. 1985. Codon usage and tRNA content in unicellular and multicellular organisms. Mol Biol Evol. 2(1):13–34. doi:10.1093/oxfordjournals.molbev.a040335.

Irony-Tur Sinai M, Salamon A, Stanleigh N, Goldberg T, Weiss A, Wang Y-H, Kerem B. 2019. AT-dinucleotide rich sequences drive fragile site formation. Nucleic Acids Res. 47(18):9685–9695. doi:10.1093/nar/gkz689.

Jackson B, Charlesworth B. 2021. Evidence for a force favoring GC over AT at short intronic sites in *Drosophila simulans* and *Drosophila melanogaster*. G3. 11(9):jkab240. doi:10.1093/g3journal/jkab240.

Jackson BC, Campos JL, Haddrill PR, Charlesworth B, Zeng K. 2017. Variation in the Intensity of Selection on Codon Bias over Time Causes Contrasting Patterns of Base Composition Evolution in Drosophila. Genome Biol Evol. 9(1):102–123. doi:10.1093/gbe/evw291.

Jacobson KB, Farkas WR, Katze JR. 1981. Presence of queuine in *Drosophila melanogaster*: correlation of free pool with queuosine content of tRNA and effect of mutations in pteridine metabolism. Nucleic Acids Res. 9(10):2351–2366. doi:10.1093/nar/9.10.2351.

Kanaya S, Yamada Y, Kinouchi M, Kudo Y, Ikemura T. 2001. Codon Usage and tRNA Genes in Eukaryotes: Correlation of Codon Usage Diversity with Translation Efficiency and with CG-Dinucleotide Usage as Assessed by Multivariate Analysis. J Mol Evol. 53(4–5):290–298. doi:10.1007/s002390010219.

Kanaya S, Yamada Y, Kudo Y, Ikemura T. 1999. Studies of codon usage and tRNA genes of 18 unicellular organisms and quantification of *Bacillus subtilis* tRNAs: gene expression level and species-specific diversity of codon usage based on multivariate analysis. Gene. 238(1):143–155. doi:10.1016/s0378-1119(99)00225-5.

Kasai H, Kuchino Y, Nihei K, Nishimura S. 1975. Distribution of the modified nucleoside Q and its derivatives in animal and plant transfer RNA’s. Nucleic Acids Res. 2(10):1931–1939. doi:10.1093/nar/2.10.1931.

Kern AD, Begun DJ. 2005. Patterns of polymorphism and divergence from noncoding sequences of *Drosophila melanogaster* and *D. simulans*: evidence for nonequilibrium processes. Mol Biol Evol. 22(1):51–62. doi:10.1093/molbev/msh269.

Kliman RM. 1999. Recent Selection on Synonymous Codon Usage in Drosophila. J Mol Evol. 49(3):343–351. doi:10.1007/pl00006557.

Kliman RM. 2014. Evidence that natural selection on codon usage in *Drosophila pseudoobscura* varies across codons. G3 Genes Genomes Genet. 4(4):681–692. doi:10.1534/g3.114.010488.

Kokate PP, Techtmann SM, Werner T. 2021. Codon usage bias and dinucleotide preference in 29 Drosophila species. G3 Bethesda Md. 11(8):jkab191. doi:10.1093/g3journal/jkab191.

Komar AA. 2021. A Code Within a Code: How Codons Fine-Tune Protein Folding in the Cell. Biochem Biokhimiia. 86(8):976–991. doi:10.1134/S0006297921080083.

Kramer EB, Farabaugh PJ. 2007. The frequency of translational misreading errors in *E. coli* is largely determined by tRNA competition. RNA. 13(1):87–96. doi:10.1261/rna.294907.

Lack JB, Cardeno CM, Crepeau MW, Taylor W, Corbett-Detig RB, Stevens KA, Langley CH, Pool JE. 2015. The Drosophila genome nexus: a population genomic resource of 623 *Drosophila melanogaster* genomes, including 197 from a single ancestral range population. Genetics. 199(4):1229–1241. doi:10.1534/genetics.115.174664.

Langley SA, Karpen GH, Langley CH. 2014. Nucleosomes shape DNA polymorphism and divergence. Pritchard JK, editor. PLoS Genet. 10(7):e1004457. doi:10.1371/journal.pgen.1004457.

Li W-H. 1987. Models of nearly neutral mutations with particular implications for nonrandom usage of synonymous codons. J Mol Evol. 24(4):337–345. doi:10.1007/bf02134132.

Lloyd AT, Sharp PM. 1992. Evolution of codon usage patterns: the extent and nature of divergence between *Candida albicans* and *Saccharomyces cerevisiae*. Nucleic Acids Res. 20(20):5289–5295. doi:10.1093/nar/20.20.5289.

Marais G. 2003. Biased gene conversion: implications for genome and sex evolution. Trends Genet TIG. 19(6):330–338. doi:10.1016/S0168-9525(03)00116-1.

Matsumoto T, Akashi H. 2018. Distinguishing Among Evolutionary Forces Acting on Genome-Wide Base Composition: Computer Simulation Analysis of Approximate Methods for Inferring Site Frequency Spectra of Derived Mutations in Recombining Regions. G3. 8(5):1755–1769. doi:10.1534/g3.117.300512.

Matsumoto T, Akashi H, Yang Z. 2015. Evaluation of Ancestral Sequence Reconstruction Methods to Infer Nonstationary Patterns of Nucleotide Substitution. Genetics. 200(3):873–890. doi:10.1534/genetics.115.177386.

Meier F, Suter B, Grosjean H, Keith G, Kubli E. 1985. Queuosine modification of the wobble base in tRNAHis influences “*in vivo*” decoding properties. EMBO J. 4(3):823–827. doi:10.1002/j.1460-2075.1985.tb03704.x.

Moran NA. 1996. Accelerated evolution and Muller’s rachet in endosymbiotic bacteria. Proc Natl Acad Sci U S A. 93(7):2873–2878. doi:10.1073/pnas.93.7.2873.

Muthukumar S, Li C-T, Liu R-J, Bellodi C. 2024. Roles and regulation of tRNA-derived small RNAs in animals. Nat Rev Mol Cell Biol. 25(5):359–378. doi:10.1038/s41580-023-00690-z.

Muyle A, Serres-Giardi L, Ressayre A, Escobar J, Glémin S. 2011. GC-biased gene conversion and selection affect GC content in the Oryza genus (rice). Mol Biol Evol. 28(9):2695–2706. doi:10.1093/molbev/msr104.

Nei M, Gojobori T. 1986. Simple methods for estimating the numbers of synonymous and nonsynonymous nucleotide substitutions. Mol Biol Evol. 3(5):418–426. doi:10.1093/oxfordjournals.molbev.a040410.

Nussinov R. 1981. Eukaryotic dinucleotide preference rules and their implications for degenerate codon usage. J Mol Biol. 149(1):125–131. doi:10.1016/0022-2836(81)90264-3.

Ohta T. 1993. An examination of the generation-time effect on molecular evolution. Proc Natl Acad Sci U S A. 90(22):10676–10680. doi:10.1073/pnas.90.22.10676.

Poh Y-P, Ting C-T, Fu H-W, Langley CH, Begun DJ. 2012. Population Genomic Analysis of Base Composition Evolution in *Drosophila melanogaster*. Genome Biol Evol. 4(12):1245–1255. doi:10.1093/gbe/evs097.

Pool JE, Corbett-Detig RB, Sugino RP, Stevens KA, Cardeno CM, Crepeau MW, Duchen P, Emerson JJ, Saelao P, Begun DJ, et al. 2012. Population Genomics of Sub-Saharan *Drosophila melanogaster*: African Diversity and Non-African Admixture. Malik HS, editor. PLoS Genet. 8(12):e1003080. doi:10.1371/journal.pgen.1003080.

Powell JR, Sezzi E, Moriyama EN, Gleason JM, Caccone A. 2003. Analysis of a Shift in Codon Usage in Drosophila. J Mol Evol. 57(1):S214–S225. doi:10.1007/s00239-003-0030-3.

Precup J, Parker J. 1987. Missense misreading of asparagine codons as a function of codon identity and context. J Biol Chem. 262(23):11351–11355. doi:10.1016/S0021-9258(18)60966-4.

Rogers RL, Cridland JM, Shao L, Hu TT, Andolfatto P, Thornton KR. 2014. Landscape of standing variation for tandem duplications in *Drosophila yakuba* and *Drosophila simulans*. Mol Biol Evol. 31(7):1750–1766. doi:10.1093/molbev/msu124.

Sawyer SA, Dykhuizen DE, Hartl DL. 1987. Confidence interval for the number of selectively neutral amino acid polymorphisms. Proc Natl Acad Sci. 84(17):6225–6228. doi:10.1073/pnas.84.17.6225.

Sharp PM, Devine KM. 1989. Codon usage and gene expression level in *Dictyosteiium discoidtum*: highly expressed genes do “prefer” optimal codons. Nucleic Acids Res. 17(13):5029–5040. doi:10.1093/nar/17.13.5029.

Sharp PM, Emery LR, Zeng K. 2010. Forces that influence the evolution of codon bias. Philos Trans R Soc Lond B Biol Sci. 365(1544):1203–1212. doi:10.1098/rstb.2009.0305.

Shields DC, Sharp PM, Higgins DG, Wright F. 1988. Silent sites in Drosophila genes are not neutral: evidence of selection among synonymous codons. Mol Biol Evol. 5(6):704–716. doi:10.1093/oxfordjournals.molbev.a040525.

Sievers A, Sauer L, Bisch M, Sprengel J, Hausmann M, Hildenbrand G. 2023. Moderation of Structural DNA Properties by Coupled Dinucleotide Contents in Eukaryotes. Genes. 14(3):755. doi:10.3390/genes14030755.

Singh ND, Arndt PF, Clark AG, Aquadro CF. 2009. Strong Evidence for Lineage and Sequence Specificity of Substitution Rates and Patterns in Drosophila. Mol Biol Evol. 26(7):1591–1605. doi:10.1093/molbev/msp071.

Sprengelmeyer QD, Mansourian S, Lange JD, Matute DR, Cooper BS, Jirle EV, Stensmyr MC, Pool JE. 2020. Recurrent Collection of *Drosophila melanogaster* from Wild African Environments and Genomic Insights into Species History. Mol Biol Evol. 37(3):627–638. doi:10.1093/molbev/msz271.

Stenico M, Lloyd AT, Sharp PM. 1994. Codon usage in *Caenorhabditis elegans*: delineation of translational selection and mutational biases. Nucleic Acids Res. 22(13):2437–2446. doi:10.1093/nar/22.13.2437.

Tavaré S. 1986. Some Probabilistic and Statistical Problems in the Analysis of DNA Sequences. In: Miura RM, editor. Some Mathematical Questions in Biology: DNA Sequence Analysis. Vol. 17. Lectures on Mathematics in the Life Sciences. p. 57–86.

Varenne S, Buc J, Lloubes R, Lazdunski C. 1984. Translation is a non-uniform process. Effect of tRNA availability on the rate of elongation of nascent polypeptide chains. J Mol Biol. 180(3):549–576. doi:10.1016/0022-2836(84)90027-5.

Vicario S, Moriyama EN, Powell JR. 2007. Codon usage in twelve species of Drosophila. BMC Evol Biol. 7(1):1–17. doi:10.1186/1471-2148-7-226.

Wang X, Matuszek Z, Huang Y, Parisien M, Dai Q, Clark W, Schwartz MH, Pan T. 2018. Queuosine modification protects cognate tRNAs against ribonuclease cleavage. RNA. 24(10):1305–1313. doi:10.1261/rna.067033.118.

White BN, Tener GM, Holden J, Suzuki DT. 1973a. Activity of a transfer RNA modifying enzyme during the development of Drosophila and its relationship to the su(s) locus. J Mol Biol. 74(4):635–651. doi:10.1016/0022-2836(73)90054-5.

White BN, Tener GM, Holden J, Suzuki DT. 1973b. Analysis of tRNAs during the development of Drosophila. Dev Biol. 33(1):185–195. doi:10.1016/0012-1606(73)90173-5.

Workman C, Krogh A. 1999. No evidence that mRNAs have lower folding free energies than random sequences with the same dinucleotide distribution. Nucleic Acids Res. 27(24):4816–4822. doi:10.1093/nar/27.24.4816.

Wright S. 1938. The Distribution of Gene Frequencies Under Irreversible Mutation. Proc Natl Acad Sci U S A. 24(7):253–259. doi:10.1073/pnas.24.7.253.

Wright SI, Yau CBK, Looseley M, Meyers BC. 2004. Effects of gene expression on molecular evolution in *Arabidopsis thaliana* and *Arabidopsis lyrata*. Mol Biol Evol. 21(9):1719–1726. doi:10.1093/molbev/msh191.

Yang J-R, Chen X, Zhang J. 2014. Codon-by-codon modulation of translational speed and accuracy via mRNA folding. PLoS Biol. 12(7):e1001910. doi:10.1371/journal.pbio.1001910.

Yang Z. 2007. PAML 4: phylogenetic analysis by maximum likelihood. Mol Biol Evol. 24(8):1586–1591. doi:10.1093/molbev/msm088.

Yıldırım B, Vogl C. 2024. The influence of GC-biased gene conversion on nonadaptive sequence evolution in short introns of *Drosophila melanogaster*. J Evol Biol. 37(4):383–400. 10.1093/jeb/voae015.

Zaborske JM, DuMont VLB, Wallace EWJ, Pan T, Aquadro CF, Drummond DA. 2014. A nutrient-driven tRNA modification alters translational fidelity and genome-wide protein coding across an animal genus. Malik HS, editor. PLoS Biol. 12(12):e1002015. doi:10.1371/journal.pbio.1002015.

Zallot R, Brochier-Armanet C, Gaston KW, Forouhar F, Limbach PA, Hunt JF, Crécy-Lagard V de. 2014. Plant, Animal, and Fungal Micronutrient Queuosine Is Salvaged by Members of the DUF2419 Protein Family. ACS Publ. 9(8):1812–1825. doi:10.1021/cb500278k.

