## Supplemental Material for "Dinucleotide preferences underlie apparent codon preference reversals in the *Drosophila melanogaster* lineage"

**Table S1. Abbreviations used in body text.**

| Abbreviation | Definition |
|---|---|
| $2f_{non\text{-}NAY}$ | 2-fold synonymous families excluding NAY codons (*i.e.*, Phe, Cys, $Ser_2$, Lys, Gln, and Glu) |
| $4f_5$ | 4-fold synonymous families excluding Gly codons (*i.e.*, Val, $Ser_4$, Pro, Thr, and Ala) |
| aDAF | Average (mean) derived allele frequency |
| BTW | Bifurcating tree with weighting method (Matsumoto and Akashi 2018) |
| $CUB_{Chi/L}$ | Codon usage bias statistic. $\chi^2$ (Chi) is calculated for the deviation of synonymous codon usage from an expectation (based on short intron GC content) and is divided by the number of codons (L). |
| $CUB_{Chi/L\_binexp}$ | $CUB_{Chi/L}$ that uses GC content of GC bin-specific short introns for calculating expected values |
| gBGC | GC-biased gene conversion |
| INCC | Internal node codon configurations. |
| INNC | Internal node nucleotide configurations. |
| LI | Long introns: lengths >100 bp in both *Dmel* and *Dsim* references |
| MCP | Major codon preference |
| MH | Mantel-Haenszel test |
| MWU | Mann-Whitney U test |
| NAY | Codons encoding Asp, Asn, His, or Tyr |
| Q | queuosine |
| SFS | Site frequency spectrum |
| skew | an index calculated as $(a - b) / (a + b)$. A value of zero indicates symmetry and the index is scaled identically for deviations in both directions. |
| SI | Short introns: lengths ≤100 bp in both *Dmel* and *Dsim* references |
| TNNC | Terminal node nucleotide configurations |
| WSR | Wilcoxon signed-rank test |
| W→S aDAF skew | Skew statistic where *a*=GC-increasing (W→S) and *b*=GC-decreasing (S→W) mutations. |
| W→S fixation skew | Skew statistic, where *a*=W→S fixation count and *b*=S→W fixation count. "$d_{up,pu}$" in Akashi *et al.* (2006). |
| W→S γ | Polymorphism-based estimate of GC fixation bias (Glémin *et al.* 2015, B in this paper). |

Note: Entries are listed in alphabetical order.

**Table S2. Species name abbreviations.**

| Abbreviation | Species name |
|---|---|
| *Dmel* | *D. melanogaster* |
| *Dsim* | *D. simulans* |
| *Dyak* | *D. yakuba* |
| *Dere* | *D. erecta* |
| *Dana* | *D. ananassae* |
| *Dpse* | *D. pseudoobscura* |
| *Dwil* | *D. willistoni* |
| *Dgri* | *D. grimshawi* |
| *Dmoj* | *D. mojavensis* |
| *Dvir* | *D. virilis* |

Note: Abbreviations for species names are used in figures and in the Supplemental Material.

### Supplementary Results and Discussion

#### *GC fixation biases on intron mutations*

##### SFS comparisons

We employed Mann-Whitney *U* (MWU) tests to compare site frequency spectrum (SFS) between forward and reverse mutations. Abbreviations used in main text and Supplemental Material are summarized in Table S1. We use the terms "forward" (W→S and Y→R changes) and "reverse" to indicate directions of mutations (labeling is arbitrary). The statistical power of this approach to detect weak fixation biases was examined by Akashi (1999). Because the counts in our SFS are not integers, all counts were scaled by a factor of 100 and the resulting test statistic was adjusted accordingly (scaled down by the same factor).

We employed aDAF skew as an index for the magnitude of difference between two SFS. This statistic compares mean derived allele frequencies between forward and reverse changes. aDAF skew values show a strong association with W→S $\gamma$ (Fig. S1), and serve as an alternative measure of the magnitude of SFS difference. aDAF skew does not estimate a population genetic parameter but this statistic has an important advantage in not requiring neutral reference data. Putatively neutral mutations, GC-conservative changes within short introns for this study, are a small fraction of the Drosophila genome.

**Fig. S1. Derived allele frequency skew is strongly correlated with population genetic model-based fixation bias estimates.**

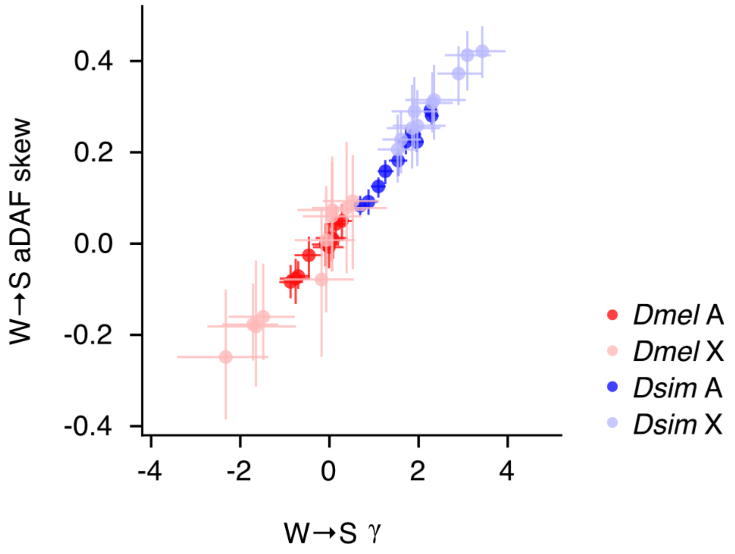

Comparisons of fixation bias statistics, W→S γ and aDAF skew, for mutations at 2-fold redundant synonymous sites and short intron (SI) sites. Each point is from a separate synonymous family. Intron GC-conservative mutations within chromosomal class and species were employed as a neutral reference for γ estimation (*e.g.*, Data from autosomal SI is employed for γ estimation for other mutation classes at autosomal loci for *Dmel*). Ancestral states were inferred separately for each bootstrap replicate. Error bars indicate 95% CIs among 300 bootstrap replicates.

**Fixation bias tests: mutation classes at intron sites**

Although nucleotide changes within short introns (SI changes) have been considered the best candidates for neutral evolution in Drosophila (Halligan and Keightley 2006; Parsch *et al.* 2010), a growing number of studies support directional forces on intron mutations (Kliman 2014; Jackson *et al.* 2017; Jackson and Charlesworth 2021; Yıldırım and Vogl 2024). Consistent with previous studies, our SFS analyses detected GC-favoring fixation bias acting on pooled SI changes in *Drosophila simulans* (*Dsim*, see Table S2 for others; Fig. 3). Furthermore, our results identified GC-favoring fixation biases acting on each of four GC-altering change pairs in *Dsim* at both autosomal and X-linked SI (Fig. S2; Table S3; autosomal T↔C, MWU test $p < 10^{-10}$; autosomal A↔G, MWU test $p < 10^{-5}$; autosomal T↔G, MWU test $p = 0.00015$; autosomal A↔C, MWU test $p = 0.0035$; X-linked T↔C, MWU test $p < 10^{-4}$; X-linked A↔G, MWU test $p < 10^{-5}$, X-linked T↔G, MWU test $p = 0.0012$, X-linked A↔C, MWU test $p < 10^{-4}$). Similar analyses reveal GC-favoring fixation biases within long introns (LI) at both autosomal and X-linked loci in *Dsim* (Fig. S2; Table S3; MWU test $p$ values were $< 10^{-10}$ for each GC-altering class at both autosomal and X-linked LI).

Because SI polymorphism data were insufficient for fixation bias inference in *D. melanogaster* (*Dmel*), we focused on LI changes. GC-fixation biases are strongly supported at both autosomal and X-linked LI in *Dmel* (except for T↔C at autosomal LI; Table S4); W→S are segregating at higher frequencies than S→W within each GC-altering change pair (autosomal T↔C, MWU test $p = 0.052$; autosomal A↔G, MWU test $p < 10^{-5}$; autosomal T↔G, MWU test $p < 10^{-10}$; autosomal A↔C, MWU test $p < 10^{-5}$; X-linked T↔C, MWU test $p = 0.00066$; X-linked A↔G, MWU test $p = 0.0024$, X-linked T↔G, MWU test $p = 0.00014$, X-linked A↔C, MWU test $p = 0.00049$).

We examined X-effects for GC-favoring fixation biases in *Dmel* using polymorphisms at LI sites. We employed a permutation approach to test the null hypothesis that the magnitudes of GC fixation biases at X-linked and autosomal loci are the same (see below). We employed W→S aDAF skew as a GC fixation bias estimate and examined differences in this statistic between X-linked and autosomal LI. The *p* values from the permutation approach were combined using Fisher's method (Fisher 1954). X-linked LI show significantly greater W→S aDAF skew than autosomal LI in *Dmel* (T→C $p = 0.001$, A→G $p = 0.047$, T→G $p = 0.22$, A→C $p = 0.13$, combined $p = 0.00067$), supporting X-effects for GC-favoring fixation biases at LI in *Dmel*.

Previous studies have employed GC-conservative mutations as an assumed neutral reference for Drosophila species (Maside *et al.* 2004; Galtier *et al.* 2006; Haddrill and Charlesworth 2008; Jackson *et al.* 2017; Jackson and Charlesworth 2021; Yıldırım and Vogl 2024) but have not explicitly tested this assumption. We found no evidence for fixation biases among GC-conservative mutations (G↔C and A↔T) at both SI and LI sites; SFS are statistically indistinguishable between forward and reverse changes within this mutation class in autosomal and X-linked introns in both species (Tables S3 and S4). We employ GC-conservative mutations at SI sites as a neutral reference in γ estimation. We pooled GC-conservative mutations into a single class.

**Fig. S2. SFS-based GC fixation biases at intron sites in *D. simulans* and in *D. melanogaster*.**

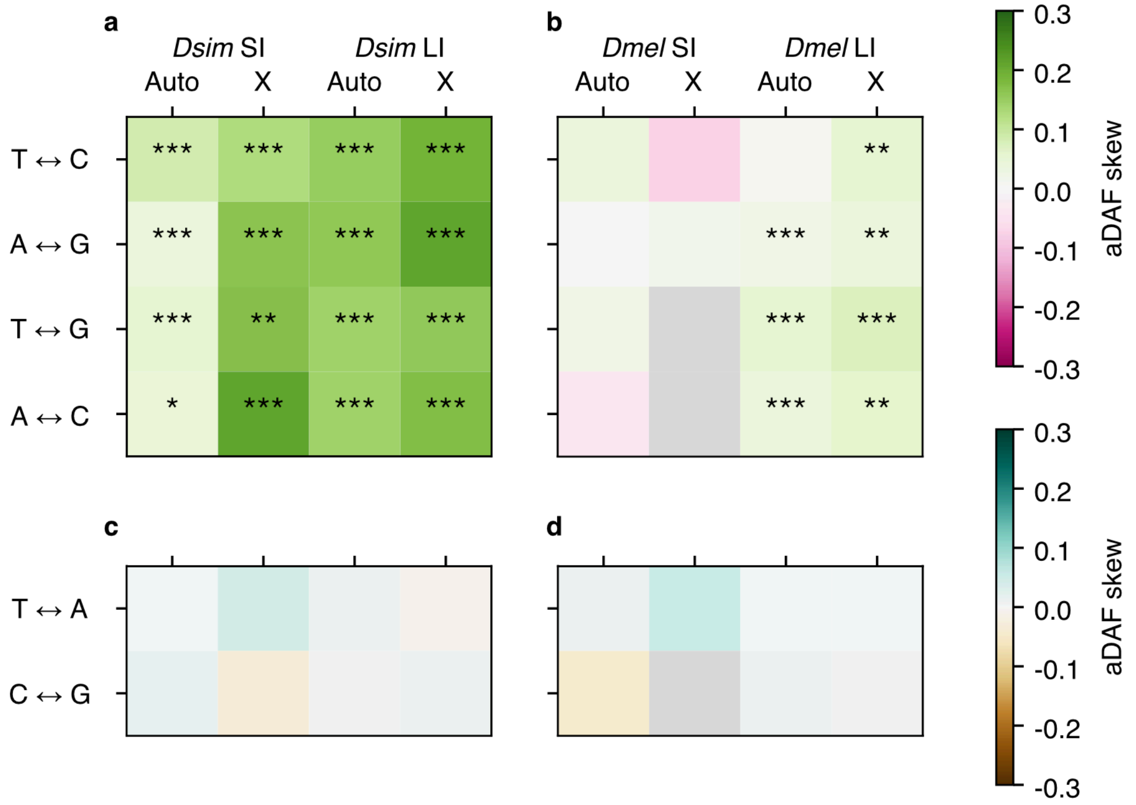

aDAF skew statistics are employed as summary statistics for magnitudes of SFS differences between forward and reverse mutations. “Forward” indicates “left” to “right” changes (*e.g.*, T→C for T↔C). Positive aDAF skew values indicate that forward mutations show higher DAF values compared with reverse mutations. (a) G/C-altering intron mutations in *D. simulans* (*Dsim*), (b) G/C-altering intron mutations in *D. melanogaster* (*Dmel*), (c) G/C-conservative intron mutations in *Dsim*, and (d) G/C-conservative intron mutations in *Dmel*. SI and LI indicate short introns and long introns, respectively. Gray shading indicates sample size < 200. Asterisks indicate statistical significance in SFS differences based on MWU tests (Tables S3 and S4). The sequential Bonferroni method (Holm 1979) was applied within each species and chromosome class to account for multiple tests. *, ** and *** indicate $p < 0.05$, < 0.01 and < 0.001, respectively.

**Table S3. SFS comparisons between forward and reverse changes at SI and LI sites in *D. simulans*.**

| | | SI | | | | | | LI | | | | |
|---|---|---|---|---|---|---|---|---|---|---|---|---|
| Chr [a] | Change [b] | # Forward [c] | # Reverse [c] | *z* [d] | | aDAF skew [e] | | # Forward [c] | # Reverse [c] | *z* [d] | | aDAF skew [e] |
| A | T→C | 5154.0 | 5675.0 | 7.42 | *** | 0.083 | | 81642.9 | 99273.1 | 51.79 | *** | 0.151 |
| | A→G | 4941.7 | 4865.3 | 5.45 | *** | 0.040 | | 78974.2 | 94786.8 | 55.09 | *** | 0.160 |
| | T→G | 2186.2 | 2723.8 | 3.79 | *** | 0.058 | | 33277.8 | 49802.2 | 33.00 | *** | 0.143 |
| | A→C | 2203.7 | 3313.3 | 2.92 | * | 0.044 | | 32954.3 | 49655.7 | 33.44 | *** | 0.143 |
| | T→A | 5253.7 | 5120.3 | -0.84 | | 0.002 | | 67515.9 | 67605.1 | 1.18 | | 0.010 |
| | C→G | 1521.0 | 1323.0 | -0.11 | | 0.018 | | 29255.7 | 28626.3 | -0.03 | | 0.006 |
| X | T→C | 675.8 | 762.2 | 4.09 | *** | 0.124 | | 12065.8 | 12487.2 | 24.96 | *** | 0.192 |
| | A→G | 627.3 | 549.7 | 4.69 | *** | 0.165 | | 11731.3 | 11798.7 | 25.92 | *** | 0.211 |
| | T→G | 281.0 | 310.0 | 3.23 | ** | 0.169 | | 4596.0 | 5880.0 | 13.44 | *** | 0.155 |
| | A→C | 262.0 | 357.0 | 4.27 | *** | 0.212 | | 4680.1 | 5903.9 | 14.48 | *** | 0.173 |
| | T→A | 476.3 | 460.7 | 1.67 | | 0.045 | | 8621.5 | 8535.5 | -0.07 | | -0.008 |
| | C→G | 236.4 | 172.6 | 0.91 | | -0.035 | | 3869.6 | 3654.4 | -0.15 | | 0.009 |

[a] Chromosome class: "A" for autosomal and "X" for X-linked loci. "SI" and "LI" indicate short and long introns, respectively.
[b] Nucleotide change in the "forward" direction indicated by the arrow. "Reverse" refers to the opposite direction.
[c] Numbers of polymorphic changes.
[d] MWU test statistics for SFS comparisons between forward and reverse changes. Positive *z* values indicate higher mean ranks in SFS of forward changes compared to SFS of reverse changes. The sequential Bonferroni method (Holm 1979) was employed in multiple test corrections within chromosome class within each species. *, ** and *** indicate $p < 0.05$, $< 0.01$ and $< 0.001$, respectively.
[e] Skew statistic that compares average derived allele frequency (aDAF) between forward and reverse changes, $(f - r) / (f + r)$, where $f$ is aDAF skew for forward and $r$ is aDAF skew for reverse changes. aDAF skew indicates the direction and the magnitude of differences in SFS locations between two mutation classes.

**Table S4. SFS comparisons between forward and reverse changes at SI and LI sites in *D. melanogaster*.**

| | | SI | | | | | LI | | | | |
|---|---|---|---|---|---|---|---|---|---|---|---|
| Chr [a] | Change [b] | # Forward [c] | # Reverse [c] | $z$ [d] | | aDAF skew [e] | # Forward [c] | # Reverse [c] | $z$ [d] | | aDAF skew [e] |
| A | T→C | 882.7 | 1382.3 | 1.40 | | 0.038 | 18454.9 | 34941.1 | 1.94 | | 0.009 |
| | A→G | 863.6 | 1214.4 | 0.12 | | 0.004 | 17891.2 | 33129.8 | 4.70 | *** | 0.023 |
| | T→G | 400.4 | 551.6 | 0.66 | | 0.027 | 7357.3 | 14625.7 | 7.02 | *** | 0.057 |
| | A→C | 386.6 | 697.4 | -1.77 | | -0.039 | 7430.6 | 14849.4 | 5.55 | *** | 0.040 |
| | T→A | 810.2 | 863.8 | 0.47 | | 0.013 | 15784.6 | 15825.4 | 0.10 | | 0.004 |
| | C→G | 357.7 | 317.3 | -1.48 | | -0.046 | 8972.2 | 8755.8 | 1.67 | | 0.009 |
| X | T→C | 169.4 | 359.6 | -2.13 | | -0.075 | 3469.1 | 6499.9 | 3.41 | ** | 0.051 |
| | A→G | 157.9 | 254.1 | 0.56 | | 0.013 | 3341.2 | 5955.8 | 3.04 | ** | 0.042 |
| | T→G | 69.2 | 117.8 | -0.21 | | -0.003 | 1410.5 | 2408.5 | 3.81 | *** | 0.071 |
| | A→C | 55.5 | 134.5 | -0.38 | | -0.044 | 1358.0 | 2497.0 | 3.49 | ** | 0.061 |
| | T→A | 157.4 | 148.6 | 1.15 | | 0.054 | 2833.1 | 2848.9 | 0.05 | | 0.004 |
| | C→G | 78.7 | 53.3 | -0.80 | | -0.013 | 1553.9 | 1471.1 | 0.81 | | 0.007 |

[a] Chromosome class: “A” for autosomal and “X” for X-linked loci. “SI” and “LI” indicate short and long introns, respectively.
[b] Nucleotide change in the “forward” direction indicated by the arrow. “Reverse” refers to the opposite direction.
[c] Numbers of polymorphic changes.
[d] MWU test statistics for SFS comparisons between forward and reverse changes. Positive $z$ values indicate higher mean ranks in SFS of forward changes compared to SFS of reverse changes. The sequential Bonferroni method (Holm 1979) was employed in multiple test corrections within chromosome class within each species. *, ** and *** indicate $p < 0.05$, $< 0.01$ and $< 0.001$, respectively.
[e] Skew statistic that compares average derived allele frequency (aDAF) between forward and reverse changes, $(f - r) / (f + r)$, where $f$ is aDAF skew for forward and $r$ is aDAF skew for reverse changes. aDAF skew indicates the direction and the magnitude of differences in SFS locations between two mutation classes.

**Heterogeneity in GC fixation biases among introns**

GC content associations between introns and synonymous sites ($CUB_{Chi/L}$ $2f_{non\text{-}NAY}$; Fig. S3) are consistent with previous reports (Kliman and Hey 1994; Bénitìere *et al.* 2024) and can be explained by sharing of evolutionary forces between these mutation classes. Such forces could include chromosomal region- and/or transcription-dependent mutation biases, GC-biased gene conversion (gBGC) and/or natural selection. GC fixation biases contribute to the variation in SI GC content in *Dsim* (Fig. S4; Jackson and Charlesworth 2021; Yıldırım and Vogl 2024). The GC content and reduced fixation biases for low GC introns are roughly consistent with predicted neutral equilibrium GC content of 20 ~ 32% based on nucleotide mutation rate estimates from mutation accumulation experiments and from naturally occurring polymorphism (Assaf *et al.* 2017) in *Dmel*. This support for GC fixation biases within SI does not distinguish effects of gBGC and natural selection. Resolving among these factors will be important because mutations in such regions are often used as a control class for interpreting evolutionary patterns within coding and regulatory regions.

**Fig. S3. Compositional associations: GC content at short introns and synonymous sites in distantly related Drosophila species.**

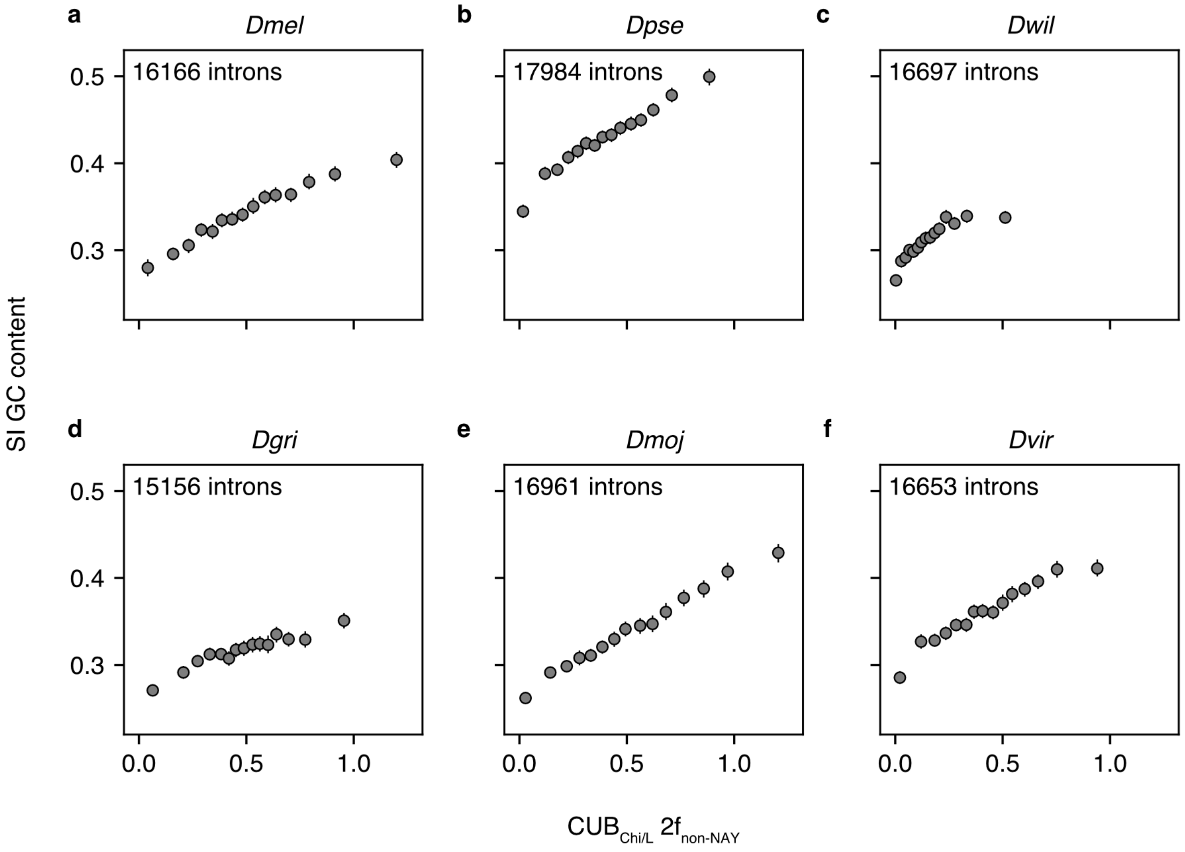

GC content of short intron (SI) sites are plotted against $CUB_{Chi/L}$ for $2f_{non\text{-}NAY}$ codons (*i.e.*, Gln, Glu, Lys, Phe, Cys, and $Ser_2$ families) for distantly related Drosophila species: (a) *D. melanogaster*, (b) *D. pseudoobscura*, (c) *D. willistoni*, (d) *D. grimshawi*, (e) *D. mojavensis*, (f) *D. virilis*. Autosomal loci are employed. Genes in which CDS has < 10 $2f_{non\text{-}NAY}$ codons are excluded. Genes are ranked by $CUB_{Chi/L}$ $2f_{non\text{-}NAY}$ and assigned to 15 bins with similar numbers of $2f_{non\text{-}NAY}$ codons. The expected GC content for $CUB_{Chi/L}$ calculation is GC content of autosomal SI. Intron sites are pooled within a bin to calculate GC content on y-axis. Error bars indicate 95% CIs among 1000 bootstrap replicates.

**Fig. S4. GC fixation bias and GC content among introns.**

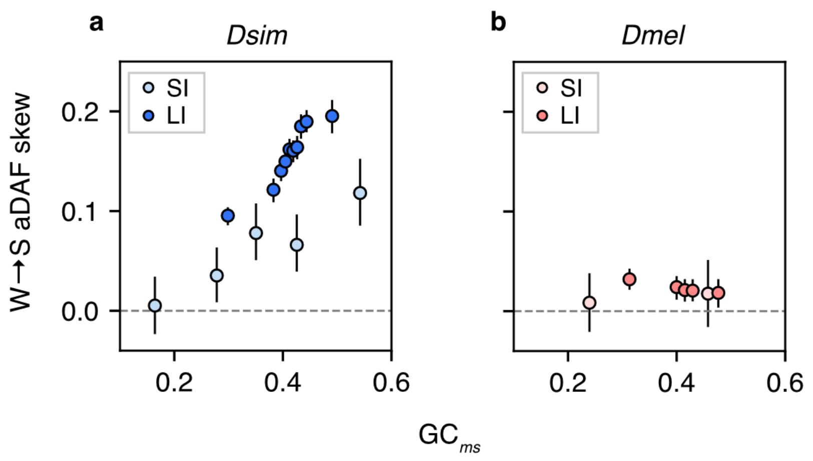

GC content at the *ms* node ($GC_{ms}$) is examined as a predictor of fixation bias, W→S aDAF skew, for short intron (SI) and long intron (LI) sites. We employed SFS from (a) *D. simulans* and (b) *D. melanogaster*. Introns are ordered by $GC_{ms}$ and classified into non-overlapping bins. The x-axis values are $GC_{ms}$ calculated from intron sites pooled within a bin. Introns with < 10 available sites are excluded. Autosomal loci are used (a total of 20,449 introns). Ancestral reconstructions were resampled in units of introns. Error bars indicate 95% CIs among 1000 bootstrap replicates.

#### *Fixation biases within 4-fold synonymous families*

SFS analysis for 2-fold synonymous families support consistent GC fixation biases in *Dsim* (Figs. 2a and 3a; Tables S5 and S6). We compared SFS for each pair of forward and reverse mutations at 4-fold redundant sites. In *Dsim*, all pairs of GC-altering mutations show GC fixation biases within five of six 4-fold synonymous families at autosomal loci and at X-linked loci (Fig. S5; Table S8). Magnitudes of fixation biases are larger at X-linked than at autosomal loci as we observed at 2-fold synonymous families (Figs. S5a and S5c; Wilcoxon signed-rank test $p = 0.00063$ among mutation classes that include 200 polymorphisms for each forward and reverse mutations). Tables S8 and S9 also show a number of cases of fixation biases acting on GC conservative mutations at 4-fold redundant sites. Although major codon preference is largely a GC-enhancing force in *Drosophila*, these results argue against the use of GC-conservative mutations in coding regions as a proxy for neutral mutations.

GC fixation biases are prevalent among synonymous sites in the *Dsim* genome with a notable exception at Gly (GGN) codons. Although GGW→GGC mutations segregate at higher frequencies than GGC→GGW, GGW→GGG segregate at lower frequencies than GGG→GGW at autosomal loci in *Dsim* (Fig. S5a; Table S8). GGC→GGG mutations also segregate at lower frequencies than GGG→GGC at autosomal loci in *Dsim* (Fig. S5a; Table S8). Similar SFS differences are found at X-linked loci for GGC↔GGG but not for GGA↔GGG and GGT↔GGG mutation pairs (Table S8).

In *Dmel*, GC fixation biases are weaker than in *Dsim* at 2-fold non-NAY codons and at most 4-fold synonymous codons (Fig. S5; Tables S7 and S9). Gly codons are also an exception in *Dmel*; GGW→GGG mutations are found at lower frequencies compared with mutations in the opposite direction at autosomal loci (Table S9). Fixation biases reduce GGG codon usage in both *Dsim* and

*Dmel*, consistent with patterns of GGG avoidance in compositional analyses for Gly codons (Shields *et al.* 1988).

Both the magnitude and direction of fixation biases vary among 4-fold synonymous families in *Dmel* and *Dsim* and we note some cases of AT preference. Such effects may have contributed to previous SFS analyses supporting overall AT preference at 4-fold redundant sites in *Dmel* (Poh *et al.* 2012). However, Poh *et al*. appear to pool data from North American and African populations in their SFS analysis. Our analyses, and those of Jackson *et al.* (2017), employed different population samples and a different ancestral inference approach from the Poh *et al*. (2012) and yielded different results for pooled 4-fold synonymous changes.

**Table S5. SFS comparisons between forward and reverse synonymous changes: pooled synonymous families within 2-fold and 4-fold redundancy cases.**

| Species [a] | Syn [b] | # W→S [c] | # S→W [c] | *z* [d] | |
|---|---|---|---|---|---|
| *Dsim* | 2-fold | 39447.5 | 103446.5 | 56.19 | *** |
| | 4-fold | 33847.6 | 105169.4 | 54.43 | *** |
| *Dmel* | 2-fold | 6346.2 | 30863.8 | -2.36 | * |
| | 4-fold | 6572.7 | 30582.3 | 7.01 | *** |

[a] "*Dsim*" and "*Dmel*" indicate *D. simulans* and *D. melanogaster*, respectively.
[b] Class of synonymous changes. "2-fold" and "4-fold" indicate 2-fold and 4-fold redundant sites in autosomal coding regions, respectively.
[c] Number of polymorphic changes.
[d] MWU test statistics from SFS comparisons between W→S and S→W changes. Positive *z* values indicate SFS of W→S changes skewed toward higher frequencies within a population compared with S→W changes. *, **, *** indicate $p < 0.05$, < 0.01, and 0.001, respectively.

**Table S6. SFS comparisons between GC-increasing and AT-increasing changes within synonymous families: 2-fold redundant sites in *D. simulans*.**

| | | A | | | | | | | X | | | | | |
|---|---|---|---|---|---|---|---|---|---|---|---|---|---|---|
| Syn [a] | Change [b] | GC (*ms*) [c] | # Forward [d] | # Reverse [d] | *z* [e] | | W→S $\gamma$ [f] | | GC (ms) [c] | # Forward [d] | # Reverse [d] | *z* [e] | | W→S $\gamma$ [f] |
| Phe | TTT→TTC | 0.655 | 4063.8 | 16559.2 | 26.95 | *** | 2.27 | | 0.713 | 515.1 | 1772.9 | 13.87 | *** | 3.43 |
| Tyr | TAT→TAC | 0.657 | 3367.2 | 7966.8 | 13.65 | *** | 1.25 | | 0.678 | 383.8 | 803.2 | 7.75 | *** | 1.91 |
| His | CAT→CAC | 0.637 | 2971.4 | 5914.6 | 8.44 | *** | 0.88 | | 0.649 | 416.5 | 641.5 | 6.21 | *** | 1.86 |
| Gln | CAA→CAG | 0.734 | 2898.9 | 12357.1 | 18.89 | *** | 1.89 | | 0.786 | 386.3 | 1365.7 | 12.37 | *** | 2.90 |
| Asn | AAT→AAC | 0.573 | 4995.3 | 8933.7 | 12.58 | *** | 1.10 | | 0.576 | 692.0 | 839.0 | 7.42 | *** | 1.61 |
| Lys | AAA→AAG | 0.732 | 4645.1 | 14428.9 | 29.53 | *** | 2.30 | | 0.800 | 463.0 | 1411.0 | 14.21 | *** | 3.09 |
| Asp | GAT→GAC | 0.480 | 6609.2 | 9070.8 | 8.36 | *** | 0.69 | | 0.507 | 864.3 | 926.7 | 7.96 | *** | 1.53 |
| Glu | GAA→GAG | 0.703 | 4537.4 | 16454.6 | 22.79 | *** | 1.96 | | 0.775 | 492.0 | 1809.0 | 11.32 | *** | 2.31 |
| Cys | TGT→TGC | 0.730 | 2315.3 | 5746.7 | 15.95 | *** | 1.71 | | 0.781 | 245.6 | 555.4 | 8.05 | *** | 2.35 |
| $Ser_2$ | AGT→AGC | 0.644 | 2983.7 | 6074.3 | 14.32 | *** | 1.54 | | 0.694 | 373.1 | 756.9 | 6.09 | *** | 1.97 |
| SI | W→S | 0.350 | 14485.6 | 16577.4 | 10.45 | *** | 0.48 | | 0.401 | 1846.0 | 1979.0 | 8.12 | *** | 1.15 |

[a] Synonymous family for CDS analysis. "SI" indicates short introns.
[b] Synonymous or intron change in the "forward" direction indicated by the arrows. "Reverse" refers to the opposite direction.
[c] The proportion of G or C at 3rd positions of codons or at intron sites. The *ms* node of Fig. 1b is employed.
[d] Number of polymorphic changes.
[e] MWU test statistics for SFS comparisons within synonymous families. Positive *z* values indicate SFS of GC-increasing mutations skewed toward higher values compared to SFS of AT-increasing mutations. Multiple test corrections were conducted within species for autosomal (A) and X-linked loci using the sequential Bonferroni method (Holm 1979). *, ** and *** indicate $p < 0.05$, $< 0.01$ and $< 0.001$, respectively.
[f] Maximum-likelihood based GC fixation bias estimates, W→S $\gamma$. Positive values indicate fixation biases to elevate GC content. SFS for GC-conservative changes at SI sites are used as a neutral reference for $\gamma$ estimation. See Fig. 3 for the comparison of W→S $\gamma$ among synonymous families.

**Table S7. SFS comparisons between GC-increasing and AT-increasing changes within synonymous families: 2-fold redundant sites in *D. melanogaster*.**

| | | A | | | | | X | | | | |
|---|---|---|---|---|---|---|---|---|---|---|---|
| Syn [a] | Change [b] | # Forward [c] | # Reverse [c] | *z* [d] | | W→S $\gamma$ [e] | # Forward [c] | # Reverse [c] | *z* [d] | | W→S $\gamma$ [e] |
| Phe | TTT→TTC | 696.1 | 4655.9 | 2.32 | | 0.14 | 91.9 | 1037.1 | 0.43 | | 0.06 |
| Tyr | TAT→TAC | 524.4 | 2601.6 | -1.31 | | -0.46 | 69.1 | 566.9 | -3.50 | ** | -2.32 |
| His | CAT→CAC | 453.2 | 1973.8 | -2.94 | * | -0.76 | 63.2 | 480.8 | -2.69 | | -1.65 |
| Gln | CAA→CAG | 480.5 | 3641.5 | -0.76 | | -0.09 | 73.5 | 881.5 | 0.45 | | -0.07 |
| Asn | AAT→AAC | 789.8 | 2854.2 | -4.75 | *** | -0.87 | 105.1 | 647.9 | -2.13 | | -1.49 |
| Lys | AAA→AAG | 733.5 | 4166.5 | 3.37 | ** | 0.28 | 96.2 | 886.8 | 1.37 | | 0.52 |
| Asp | GAT→GAC | 1004.3 | 2884.7 | -3.97 | *** | -0.70 | 176.3 | 661.7 | -3.91 | ** | -1.71 |
| Glu | GAA→GAG | 746.8 | 4746.2 | 0.28 | | 0.08 | 75.6 | 1062.4 | -1.92 | | -0.18 |
| Cys | TGT→TGC | 422.0 | 1667.0 | 0.23 | | 0.05 | 52.1 | 352.9 | 1.74 | | 0.39 |
| $Ser_2$ | AGT→AGC | 529.8 | 1638.2 | 0.26 | | -0.01 | 77.8 | 369.2 | 0.82 | | 0.06 |
| SI | W→S | 2533.4 | 3845.6 | 0.50 | | 0.13 | 452.1 | 865.9 | -1.30 | | -0.27 |

[a] Synonymous family for CDS analysis. “SI” indicates short introns.
[b] Synonymous or intron change in the “forward” direction indicated by the arrows. “Reverse” refers to the opposite direction.
[c] Number of polymorphic changes.
[d] MWU test statistics for SFS comparisons within synonymous families. Positive *z* values indicate SFS of GC-increasing mutations skewed toward higher values compared to SFS of AT-increasing mutations. Multiple test corrections were conducted within species for autosomal (A) and X-linked loci using the sequential Bonferroni method (Holm 1979). *, ** and *** indicate $p < 0.05$, $< 0.01$ and $< 0.001$, respectively.
[e] Maximum-likelihood based fixation bias estimates, W→S $\gamma$. Positive values indicate fixation biases in direction to elevate GC content. SFS for intron GC-conservative changes are used as a neutral reference for $\gamma$ estimation. See Fig. 3 for the comparison of W→S $\gamma$ among synonymous families.

**Fig. S5. Fixation biases at 4-fold redundant sites in coding regions.**

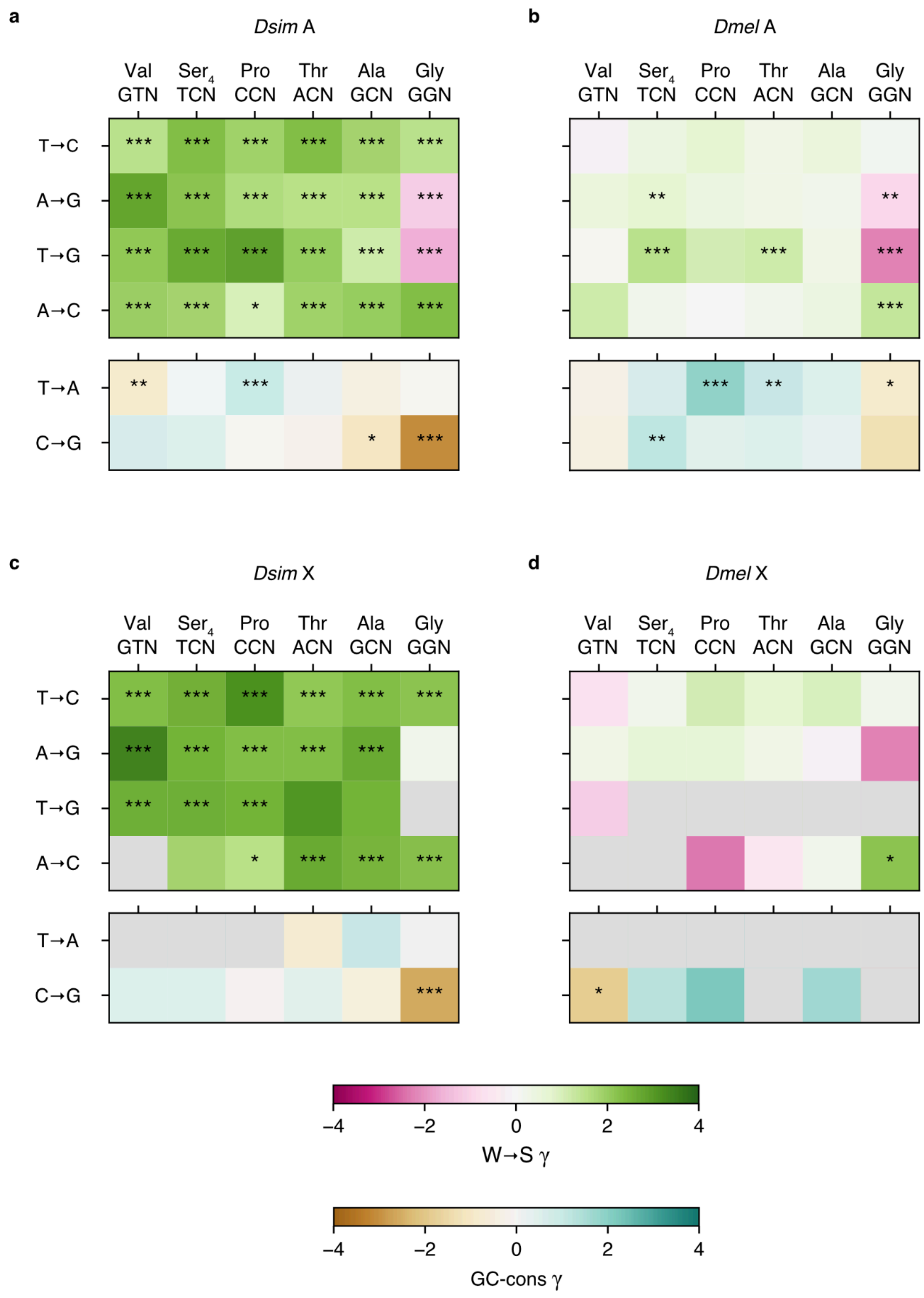

Fixation bias estimates, γ, for pairs of "forward" and "reverse" mutations in *D. simulans* (*Dsim*; a: autosomal, c: X-linked) and in *D. melanogaster* (*Dmel*; b: autosomal, d: X-linked). Forward direction is indicated by arrows and reverse is the opposite direction. Positive γ indicate fixation biases favoring forward mutations. SFS for GC-conservative mutations within short introns from autosomal and X-linked loci were employed as neutral references for γ estimation for the corresponding coding region data. GC-altering changes (the top table within a panel) and GC-conservative changes (the bottom table within a panel) are shown in different color scales: magenta-green and brown-blue green, respectively. Gray shading indicates cases with sample sizes < 200. Statistical results are from MWU tests (Tables S8 and S9 for data for *D. simulans* and *D. melanogaster*, respectively). The sequential Bonferroni method (Holm 1979) was applied in multiple test corrections within each panel. *, ** and *** indicate $p < 0.05$, $< 0.01$ and $< 0.001$, respectively

**Table S8. SFS comparisons between forward and reverse changes within 4-fold synonymous families in *D. simulans*.**

| | | A | | | | | X | | | | |
|---|---|---|---|---|---|---|---|---|---|---|---|
| Syn [a] | Change [b] | # Forward [c] | # Reverse [c] | $z$ [d] | | $\gamma$ [e] | # Forward [c] | # Reverse [c] | $z$ [d] | | $\gamma$ [e] |
| Val | GTT→GTC | 1452.8 | 4286.2 | 9.76 | *** | 1.53 | 164.9 | 505.1 | 5.40 | *** | 2.32 |
| | GTA→GTG | 1403.9 | 8413.1 | 18.72 | *** | 2.80 | 151.6 | 904.4 | 9.91 | *** | 3.43 |
| | GTT→GTG | 997.2 | 4929.8 | 12.37 | *** | 2.13 | 102.4 | 568.6 | 6.06 | *** | 2.63 |
| | GTA→GTC | 294.2 | 1524.8 | 6.63 | *** | 1.96 | 35.5 | 153.5 | 2.68 | | 3.06 |
| | GTT→GTA | 1149.8 | 1062.2 | -3.75 | ** | -0.88 | 113.1 | 82.9 | -1.86 | | -0.88 |
| | GTC→GTG | 1479.8 | 2370.2 | 2.56 | | 0.62 | 198.3 | 283.7 | 0.65 | | 0.52 |
| Ser4 | TCT→TCC | 1527.5 | 6309.5 | 16.84 | *** | 2.31 | 125.3 | 614.7 | 5.12 | *** | 2.59 |
| | TCA→TCG | 1304.4 | 4918.6 | 13.49 | *** | 2.20 | 158.3 | 643.7 | 7.29 | *** | 2.55 |
| | TCT→TCG | 522.2 | 2292.8 | 9.97 | *** | 2.72 | 49.6 | 242.4 | 4.31 | *** | 2.61 |
| | TCA→TCC | 684.0 | 2752.0 | 8.08 | *** | 1.82 | 65.9 | 264.1 | 2.20 | | 1.82 |
| | TCT→TCA | 760.6 | 762.4 | 0.77 | | 0.04 | 53.3 | 52.7 | 2.29 | | 1.39 |
| | TCC→TCG | 1945.8 | 1816.2 | 2.39 | | 0.48 | 257.2 | 259.8 | 1.75 | | 0.50 |
| Pro | CCT→CCC | 1677.5 | 5566.5 | 13.71 | *** | 1.92 | 141.9 | 580.1 | 5.50 | *** | 3.17 |
| | CCA→CCG | 2473.0 | 5946.0 | 13.17 | *** | 1.67 | 344.2 | 845.8 | 7.62 | *** | 2.33 |
| | CCT→CCG | 646.9 | 1878.1 | 11.42 | *** | 2.86 | 53.0 | 253.0 | 4.49 | *** | 2.54 |
| | CCA→CCC | 1207.2 | 3339.8 | 2.85 | * | 1.01 | 151.9 | 324.1 | 3.38 | * | 1.57 |
| | CCT→CCA | 1190.0 | 1355.0 | 4.96 | *** | 1.00 | 77.2 | 116.8 | 2.72 | | 2.50 |
| | CCC→CCG | 1941.2 | 1343.8 | 0.88 | | -0.03 | 284.3 | 211.7 | -0.41 | | -0.10 |

(Continue to the next page)

**Table S8.** (Continue from the last page)

| | | A | | | | | X | | | | |
|---|---|---|---|---|---|---|---|---|---|---|---|
| Syn [a] | Change [b] | # Forward [c] | # Reverse [c] | $z$ [d] | | $\gamma$ [e] | # Forward [c] | # Reverse [c] | $z$ [d] | | $\gamma$ [e] |
| Thr | ACT→ACC | 1633.7 | 6000.3 | 17.69 | *** | 2.31 | 156.8 | 652.2 | 6.82 | *** | 2.16 |
| | ACA→ACG | 1581.8 | 4813.2 | 11.49 | *** | 1.56 | 184.0 | 620.0 | 7.39 | *** | 2.33 |
| | ACT→ACG | 698.4 | 2148.6 | 8.09 | *** | 2.01 | 53.1 | 233.9 | 2.84 | | 3.11 |
| | ACA→ACC | 863.8 | 3645.2 | 9.01 | *** | 1.89 | 75.4 | 354.6 | 6.68 | *** | 2.68 |
| | ACT→ACA | 1576.4 | 1219.6 | 0.75 | | 0.19 | 107.3 | 126.7 | -1.95 | | -0.72 |
| | ACC→ACG | 1742.5 | 1098.5 | 1.08 | | -0.13 | 216.7 | 148.3 | 0.65 | | 0.42 |
| Ala | GCT→GCC | 2398.2 | 9659.8 | 16.33 | *** | 1.83 | 239.0 | 1163.0 | 8.90 | *** | 2.31 |
| | GCA→GCG | 1757.1 | 4519.9 | 10.05 | *** | 1.57 | 214.9 | 581.1 | 5.95 | *** | 2.73 |
| | GCT→GCG | 900.8 | 1959.2 | 5.46 | *** | 1.19 | 77.7 | 250.3 | 2.09 | | 2.54 |
| | GCA→GCC | 1077.6 | 4860.4 | 12.11 | *** | 2.02 | 135.8 | 546.2 | 6.18 | *** | 2.49 |
| | GCT→GCA | 1910.5 | 1665.5 | -1.27 | | -0.35 | 147.9 | 173.1 | 1.14 | | 1.04 |
| | GCC→GCG | 2418.8 | 1000.2 | -2.94 | * | -1.03 | 359.4 | 163.6 | -0.92 | | -0.44 |
| Gly | GGT→GGC | 3552.4 | 8284.6 | 17.86 | *** | 1.53 | 543.9 | 1132.1 | 10.60 | *** | 2.22 |
| | GGA→GGG | 2797.6 | 1955.4 | -8.94 | *** | -1.15 | 339.9 | 246.1 | -0.71 | | 0.24 |
| | GGT→GGG | 774.3 | 818.7 | -5.38 | *** | -1.63 | 109.2 | 82.8 | 0.83 | | 1.02 |
| | GGA→GGC | 1634.4 | 4333.6 | 16.90 | *** | 2.31 | 200.9 | 504.1 | 5.71 | *** | 2.28 |
| | GGT→GGA | 1595.7 | 2843.3 | 1.39 | | -0.03 | 183.1 | 258.9 | 0.33 | | 0.13 |
| | GGC→GGG | 1766.6 | 563.4 | -9.77 | *** | -3.03 | 260.7 | 87.3 | -4.52 | *** | -2.56 |

(See the next page for footnotes.)

[a] Synonymous family.
[b] Synonymous change in the "forward" direction indicated by the arrows. "Reverse" means the opposite direction.
[c] Numbers of polymorphic changes.
[d] MWU test statistics for SFS comparisons within synonymous families. Positive *z* values indicate SFS of forward changes skewed toward higher values compared with that of reverse changes. The sequential Bonferroni method (Holm 1979) was employed in multiple test corrections within each species. *, ** and *** indicate $p < 0.05$, $< 0.01$ and $< 0.001$, respectively.
[e] Maximum-likelihood estimates of fixation biases, $\gamma$, acting between two codons. Positive values indicate fixation biases that elevate derived allele frequencies for forward mutations. See Fig. S5 for comparisons of $\gamma$ among synonymous families and between species.

**Table S9. SFS comparisons between forward and reverse changes within 4-fold synonymous families in *D. melanogaster*.**

| | | A | | | | X | | | |
|---|---|---|---|---|---|---|---|---|---|
| Syn [a] | Change [b] | # Forward [c] | # Reverse [c] | $z$ [d] | $\gamma$ [e] | # Forward [c] | # Reverse [c] | $z$ [d] | $\gamma$ [e] |
| Val | GTT→GTC | 279.9 | 1376.1 | 0.53 | -0.13 | 45.7 | 296.3 | -1.46 | -0.65 |
| | GTA→GTG | 291.3 | 2614.7 | 3.00 | 0.53 | 39.0 | 560.0 | 0.60 | 0.28 |
| | GTT→GTG | 161.2 | 1558.8 | 1.08 | 0.08 | 18.8 | 322.2 | -1.02 | -1.10 |
| | GTA→GTC | 68.8 | 425.2 | 2.30 | 1.21 | 10.1 | 78.9 | 0.94 | 1.12 |
| | GTT→GTA | 238.8 | 288.2 | -0.64 | -0.24 | 40.3 | 37.7 | -0.94 | -0.09 |
| | GTC→GTG | 333.1 | 704.9 | -1.15 | -0.37 | 62.9 | 158.1 | -3.53 * | -1.84 |
| Ser4 | TCT→TCC | 234.1 | 1936.9 | 1.49 | 0.44 | 26.8 | 395.2 | 1.02 | 0.25 |
| | TCA→TCG | 344.2 | 1513.8 | 3.60 ** | 0.74 | 45.4 | 389.6 | 0.90 | 0.61 |
| | TCT→TCG | 95.1 | 481.9 | 4.17 *** | 1.52 | 19.0 | 94.0 | 1.81 | 2.35 |
| | TCA→TCC | 153.7 | 863.3 | 0.27 | 0.22 | 9.7 | 169.3 | -0.19 | -0.48 |
| | TCT→TCA | 172.2 | 176.8 | 1.49 | 0.61 | 42.2 | 16.8 | 0.95 | 2.45 |
| | TCC→TCG | 633.5 | 376.5 | 3.99 ** | 1.23 | 125.0 | 79.0 | 1.65 | 1.25 |
| Pro | CCT→CCC | 289.4 | 1564.6 | 3.07 | 0.69 | 35.7 | 302.3 | 0.27 | 1.07 |
| | CCA→CCG | 520.8 | 1609.2 | 1.39 | 0.36 | 98.9 | 427.1 | 1.09 | 0.61 |
| | CCT→CCG | 169.9 | 442.1 | 2.89 | 1.07 | 19.7 | 91.3 | 1.27 | 0.77 |
| | CCA→CCC | 227.9 | 1003.1 | 0.16 | -0.07 | 27.1 | 215.9 | -2.67 | -2.35 |
| | CCT→CCA | 334.7 | 261.3 | 5.12 *** | 1.87 | 42.4 | 37.6 | 3.06 | 2.28 |
| | CCC→CCG | 573.0 | 325.0 | 1.40 | 0.44 | 137.6 | 63.4 | 3.11 | 2.21 |

(Continue to the next page)

**Table S9.** (Continue from the last page)

| | | A | | | | | X | | | | |
|---|---|---|---|---|---|---|---|---|---|---|---|
| Syn [a] | Change [b] | # Forward [c] | # Reverse [c] | $z$ [d] | | $\gamma$ [e] | # Forward [c] | # Reverse [c] | $z$ [d] | | $\gamma$ [e] |
| Thr | ACT→ACC | 259.1 | 1792.9 | 0.91 | | 0.34 | 37.3 | 375.7 | 1.28 | | 0.69 |
| | ACA→ACG | 317.7 | 1531.3 | 1.56 | | 0.30 | 55.8 | 365.2 | -0.22 | | 0.28 |
| | ACT→ACG | 149.8 | 521.2 | 4.36 | *** | 1.17 | 20.8 | 83.2 | 1.50 | | 1.49 |
| | ACA→ACC | 169.0 | 1039.0 | 1.26 | | 0.21 | 25.2 | 202.8 | -1.63 | | -0.51 |
| | ACT→ACA | 440.7 | 261.3 | 3.89 | ** | 1.03 | 59.0 | 44.0 | 2.38 | | 2.67 |
| | ACC→ACG | 514.1 | 265.9 | 1.64 | | 0.51 | 101.0 | 58.0 | 2.20 | | 1.59 |
| Ala | GCT→GCC | 417.9 | 2764.1 | 2.09 | | 0.51 | 68.8 | 608.2 | 2.08 | | 0.99 |
| | GCA→GCG | 357.2 | 1274.8 | 2.03 | | 0.21 | 60.8 | 340.2 | -1.11 | | -0.22 |
| | GCT→GCG | 139.6 | 525.4 | 0.44 | | 0.31 | 19.2 | 109.8 | -2.28 | | -1.53 |
| | GCA→GCC | 219.5 | 1413.5 | 1.64 | | 0.38 | 33.5 | 269.5 | 0.58 | | 0.21 |
| | GCT→GCA | 471.1 | 368.9 | 1.86 | | 0.48 | 66.6 | 47.4 | 0.63 | | 0.62 |
| | GCC→GCG | 638.9 | 268.1 | 0.47 | | 0.28 | 146.6 | 73.4 | 2.47 | | 1.67 |
| Gly | GGT→GGC | 656.3 | 2467.7 | 1.26 | | 0.14 | 145.9 | 693.1 | -0.57 | | 0.22 |
| | GGA→GGG | 597.8 | 586.2 | -4.13 | ** | -0.90 | 77.1 | 148.9 | -3.17 | | -2.19 |
| | GGT→GGG | 161.4 | 249.6 | -5.56 | *** | -2.24 | 22.2 | 49.8 | -1.19 | | -1.02 |
| | GGA→GGC | 290.8 | 1027.2 | 5.81 | *** | 1.36 | 48.4 | 207.6 | 3.25 | * | 2.21 |
| | GGT→GGA | 369.3 | 676.7 | -3.37 | * | -0.89 | 52.2 | 147.8 | -1.23 | | -0.81 |
| | GGC→GGG | 457.0 | 145.0 | -1.86 | | -1.32 | 93.3 | 22.7 | -2.60 | | -2.51 |

(See the next page for footnotes.)

[a] Synonymous family.
[b] Synonymous change in the "forward" direction indicated by the arrows. "Reverse" means the opposite direction.
[c] Numbers of polymorphic changes.
[d] MWU test statistics for SFS comparisons within synonymous families. Positive *z* values indicate SFS of forward changes skewed toward higher values compared with that of reverse changes. The sequential Bonferroni method (Holm 1979) was employed in multiple test corrections within each species. *, ** and *** indicate $p < 0.05$, $< 0.01$ and $< 0.001$, respectively.
[e] Maximum-likelihood estimates of fixation biases, $\gamma$, acting between two codons. Positive values indicate fixation biases that elevate derived allele frequencies for forward mutations. See Fig. S5 for comparisons of $\gamma$ among synonymous families and between species.

#### *GC content evolution in the* D. simulans *and* D. melanogaster *ancestral lineages*

##### W→S fixation skew differences among mutation classes

We tested the independence of the ratios of W→S ($N_{W\rightarrow S}$) and S→W ($N_{S\rightarrow W}$) fixation counts between mutation classes (Table S10). Because these ratios are expected to depend on ancestral GC content (Akashi *et al.* 2007), we constructed separate tables for bins of sequences (CDS or introns) with similar GC content at the *ms* node ($GC_{ms}$). We adjusted the $GC_{ms}$ range for each bin to limit the range while allowing for sufficient sample sizes.

We employed the Mantel-Haenszel (MH) approach (Mantel and Haenszel 1959; Mantel 1963) to test independence of the GC-altering fixation count ratios across $GC_{ms}$ bins. We also employed Wilcoxon signed rank (WSR) tests to detect consistent differences in GC fixation skews between two classes. We applied the tests to each pair among the following mutation classes: SI, NAY, $2f_{non\text{-}NAY}$, and "$4f_5$", synonymous changes at 4-fold redundant sites excluding Gly codons (*i.e.*, Ala, $Ser_4$, Pro, Thr, Val). We excluded Gly codons because our SFS analyses support Gly-specific fixation biases consistently disfavoring G-ending codons (see Fixation biases in 4-fold synonymous families).

The ancestral *Dsim* and *Dmel* lineages show GC content reduction for all mutation classes and almost all ancestral GC ranges (Fig. 4). In *Dsim*, although the overall trends seem similar among mutation classes, $2f_{non\text{-}NAY}$ families show a slight difference in W→S fixation skews from the other mutation classes (Figs. 4a and S6c); $2f_{non\text{-}NAY}$ codons show support for less GC reduction compared to $4f_5$ and NAY codons (Table S11). $4f_5$ is not distinguishable from changes at SI sites and NAY codons in *Dsim* (Table S11). Such patterns are roughly consistent with shared parameter changes at SI, $4f_5$, and NAY classes in the ancestral *Dsim* lineage.

We detected heterogeneity in patterns of GC reduction among mutation classes in the longer *Dmel* ancestral lineage. W→S fixation skews are not distinguishable between SI sites and $2f_{non\text{-}NAY}$ codons (Fig. 4c; Table S11) and we refer to the trend for these classes as the “base” trend for autosomal loci in *Dmel*. $4f_5$ changes show similar slopes to the base pattern but with consistently smaller GC reductions (Fig. S6d; Table S11). The NAY mutation class shows more substantial GC differences (reductions among X-linked loci) compared to the base trend (see main text, Fig. S6d, and Table S11).

We compared W→S fixation skew data to predictions under major codon preference scenarios of non-stationary mutation ratio and/or fixation bias (Akashi *et al.* 2007). Consider equilibrium GC content at an ancestral node; mutation bias is shared, but GC fixation bias varies, among genes. Scaling factors are applied to mutation and GC fixation bias parameters. The fixation bias scaling factor is shared among genes so that all genes experience the same proportionate change ($N_e$ fluctuation scenarios fit this model). Evolution proceeds under the new parameters which remain constant within the lineage. The mutation ratio parameter determines the y-intercept and the fixation bias parameter largely determines the slope of the trends and these parameters can be adjusted to fit the *Dmel* and *Dsim* GC fixation bias patterns (Figs. S6a and S6b).

A scenario of mutation bias change toward high AT and reduced GC fixation bias seems to fit W→S fixation skews for SI, NAY, $2f_{non\text{-}NAY}$, and $4f_5$ classes for the *Dsim* ancestral lineage (Fig. S6c). LI shows a distinct trend compared to other mutation classes (Fig. S6c); W→S fixation skews remain roughly constant across $GC_{ms}$ bins. A proportionate $N_e$ change cannot explain the difference in W→S fixation skews between mutation classes within the genome. The result suggests that selection coefficient or biased-gene conversion parameter may be different for GC-altering mutations between SI and LI sites. This scenario is consistent with SFS-based fixation bias estimates that supported stronger GC fixation biases at LI than at SI sites in *Dsim* (Fig. S4a). However, the factors underlying the fixation bias heterogeneity between SI and LI remain unclear.

In the *Dmel* ancestral lineage, a similar mutation bias scaling with more severe GC fixation bias reduction seems to fit W→S fixation skew trends for SI, $4f_5$, and $2f_{non\text{-}NAY}$ mutation classes (Fig. S6d). In contrast to *Dsim*, the LI pattern is similar to the trend from these mutation classes in *Dmel* (Fig. S6d). *Dmel* NAY shows greater reduction of GC compared to the trends from the other mutation classes (Fig. S6d, Table S11). We do not attempt to specify the timing or magnitude of the preference reversal because these parameters can only be estimated jointly and both are largely unconstrained.

**Table S10. General design of 2 x 2 tables comparing GC-altering fixations.**

| | class 1 | class 2 |
|---|---|---|
| W→S | $N_{W \to S,1}$ | $N_{W \to S,2}$ |
| S→W | $N_{S \to W,1}$ | $N_{S \to W,2}$ |

**Table S11. W→S fixation skew differences among mutation classes.**

| Species | Mutation class | Total $N_{W→S}$ | Total $N_{S→W}$ | MH $z$ | | WSR | | OR mean | Bin num |
|---|---|---|---|---|---|---|---|---|---|
| *Dsim* | SI | 743.1 | 938.8 | | | | | | |
| | $2f_{non\text{-}NAY}$ | 801.0 | 966.1 | 2.3 | | 6 | | 1.27 | 12 |
| | SI | 596.8 | 788.1 | | | | | | |
| | $4f_5$ | 1253.9 | 1807.0 | 0.7 | | 24 | | 1.10 | 11 |
| | SI | 1007.0 | 1175.5 | | | | | | |
| | NAY | 1361.9 | 1898.7 | 0.6 | | 85 | | 1.10 | 20 |
| | $2f_{non\text{-}NAY}$ | 3338.7 | 5629.2 | | | | | | |
| | $4f_5$ | 4989.0 | 9214.1 | -4.1 | *** | 327 | *** | 0.91 | 61 |
| | $2f_{non\text{-}NAY}$ | 2990.3 | 4794.8 | | | | | | |
| | NAY | 1854.2 | 2934.0 | -3.9 | *** | 18 | *** | 0.85 | 24 |
| | $4f_5$ | 4259.6 | 7549.8 | | | | | | |
| | NAY | 1819.9 | 2867.0 | -0.3 | | 140 | | 1.01 | 25 |
| *Dmel* | SI | 1321.5 | 2407.8 | | | | | | |
| | $2f_{non\text{-}NAY}$ | 1550.2 | 3973.3 | -1.0 | | 44 | | 0.96 | 15 |
| | SI | 1176.0 | 2197.5 | | | | | | |
| | $4f_5$ | 2504.4 | 5735.3 | 2.7 | * | 22 | | 1.19 | 15 |
| | SI | 1791.7 | 2993.2 | | | | | | |
| | NAY | 2344.8 | 7712.2 | -8.4 | *** | 8 | *** | 0.72 | 23 |
| | $2f_{non\text{-}NAY}$ | 4540.6 | 18200.6 | | | | | | |
| | $4f_5$ | 7053.5 | 24777.8 | 4.5 | *** | 626 | *** | 1.12 | 73 |
| | $2f_{non\text{-}NAY}$ | 4160.9 | 15327.3 | | | | | | |
| | NAY | 2758.9 | 10561.5 | -10.7 | *** | 0 | *** | 0.73 | 33 |
| | $4f_5$ | 6502.2 | 21011.7 | | | | | | |
| | NAY | 2709.1 | 10469.9 | -15.7 | *** | 0 | *** | 0.65 | 34 |

(See the next page for footnotes.)

Note: Each bin is set to contain at least 25 sequences (introns and CDS). The minimum site counts per bin were as follows: 4500 for short intron (SI) vs $2f_{non\text{-}NAY}$, SI vs $4f_5$, and SI vs NAY in *Dsim*, 9000 for non-NAY vs $4f_5$ in *Dsim*, 11000 for $2f_{non\text{-}NAY}$ vs NAY and $4f_5$ vs NAY in *Dsim*, 3500 for SI vs $2f_{non\text{-}NAY}$, SI vs $4f_5$, and SI vs NAY in *Dmel*, 6000 for $2f_{non\text{-}NAY}$ vs $4f_5$ in *Dmel*, and 8000 for $2f_{non\text{-}NAY}$ vs NAY and $4f_5$ vs NAY in *Dmel*. For the statistical analysis, we filtered bins with the lowest and highest $GC_{ms}$ because mean $GC_{ms}$ can vary between mutation classes in these bins. “Bin num” indicates the number of bins employed for statistical tests. Total fixation counts across the $GC_{ms}$ bins are indicated as “Total $N_{W \to S}$” and “Total $N_{S \to W}$” for GC-increasing (W→S) and GC-decreasing (S→W) changes, respectively. OR (odds ratio) indicates the departure from equal ratios of W→S fixation count to S→W fixation count between two mutation classes. OR greater 1 and positive MH *z* values indicate a high W→S fixation skew for a mutation class at the lower row compared to the other mutation class. OR less than 1 and negative MH *z* values indicate a W→S fixation skew difference in the opposite direction. OR values are averages across bins. Multiple tests were corrected among mutation class pairs for each species. We used the sequential Bonferroni method (Holm 1979) for the multiple test correction. *, **, and *** indicate $p < 0.05$, $< 0.01$, and $< 0.001$, respectively.

**Fig. S6. Non-stationary base composition: model-based and observed GC fixation skew in the *D. simulans* and *D. melanogaster* ancestral lineages.**

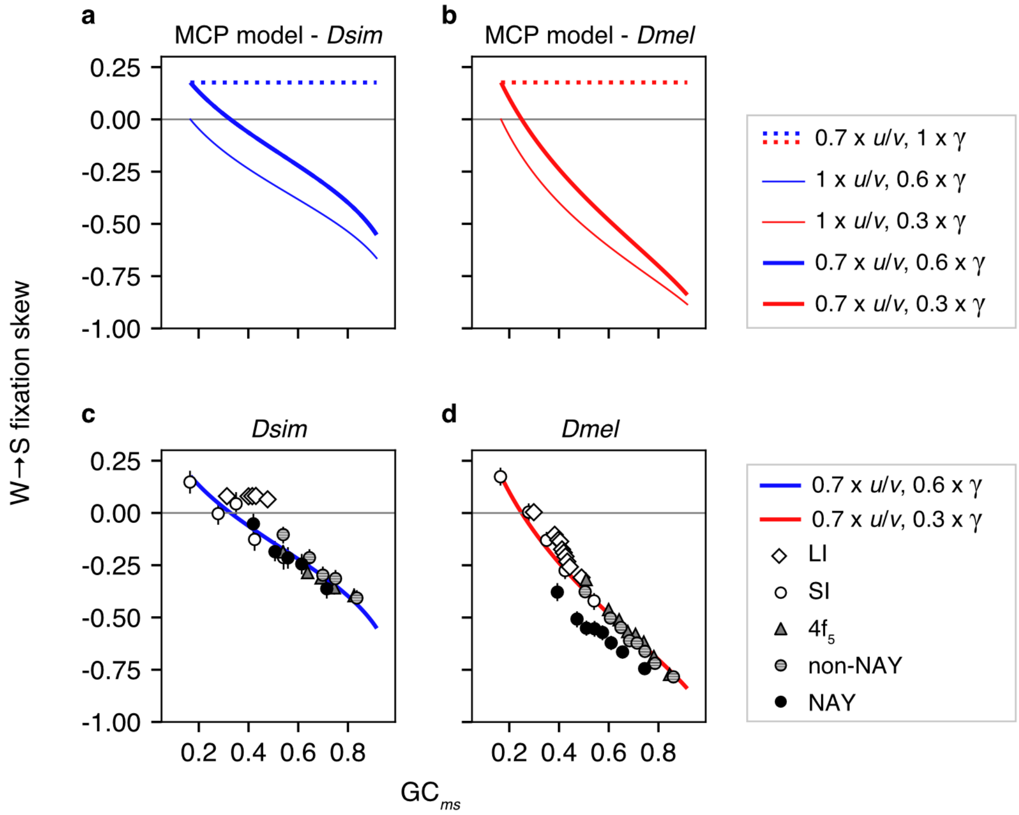

The lines depict W→S fixation skew predictions under MCP scenarios of non-stationary mutation ratio and/or fixation bias (Akashi *et al.* 2007). *u*/*v* is the mutation ratio (S→W mutation rate / W→S mutation rate) and γ is the GC fixation bias at the ancestral node, *ms*. Scaling factors were determined by eye to roughly fit features of W→S fixation skew observations in (a) *D. simulans* ("*Dsim*") and (b) *D. melanogaster* ("*Dmel*"). *u*/*v* was scaled to fit empirical W→S fixation skew for low GC short introns (SI). The solid lines show W→S γ scaling to fit the slopes of W→S fixation skews for SI, , $4f_5$, and $2f_{non\text{-}NAY}$ classes. Scaling factor values are shown in the legends. (c) and (d) show inferred W→S fixation skew for autosomal genes in the *Dmel* and *Dsim* lineages, respectively, compared to the predictions. "LI" refers to long introns. Error bars indicate 95% CIs among 1000 bootstrap replicates (some error bars are hidden by symbols). Ancestral reconstructions were resampled among replicates.

**W→S fixation skew differences between autosomal and X-linked genes**

Our SFS analysis supported stronger fixation biases at X-linked, compared to autosomal, loci in the recent histories (*i.e.*, among segregating mutations) of both *Dsim* and *Dmel* (Fig. 3; see main text) and we tested for signals of such differences in fixation bias in their ancestral lineages. We compared ratios of $N_{W\rightarrow S}$ to $N_{S\rightarrow W}$ between X-linked and autosomal loci in 2 x 2 contingency tables (Table S10). To control for magnitudes of ancestral fixation bias, we assigned autosomal and X-linked genes to $GC_{ms}$ bins and employed MH tests to evaluate the null hypothesis of the independence of GC fixation counts between X vs A. We also employed WSR tests to assess differences in GC fixation skews between these chromosomal classes.

In *Dsim*, low fixation counts from X-linked loci limit our ability to evaluate X-effects (Fig. S7a-d). The NAY class shows weak support for lower NAC loss at X-linked than at autosomal loci but this result is not supported by WSR tests (Table S12). Jackson and co-workers (2017) previously reported *less* GC content reduction at 4-fold synonymous sites at X-linked loci than at autosomal loci. However, their analysis examined single alleles from each species (*i.e.*, the examined lineage was a fusion of *ms*-*s*' and *s*'-$s_i$ where $s_i$ indicates a particular *Dsim* allele) and they examined pooled 4-fold synonymous sites (including Gly codons).

In *Dmel*, X-linked loci show greater GC loss than autosomal loci in all four mutation classes (Fig. S7e-h; Table S12). Some of these trends are similar to previously noted patterns (Singh *et al.* 2007, 2009; Jackson *et al.* 2017) but these studies employed single alleles (*i.e.*, the examined lineage was *ms*-*m*' plus *m*'-$m_i$) and pooled 4-fold redundant (Singh *et al.* 2007; Jackson *et al.* 2017) or all (Singh *et al.* 2009) synonymous families. Our analyses examined both a more specified lineage (*ms*-*m*') and more refined synonymous mutation classes.

We focus on a previously unreported X effect in GC content evolution in the *Dmel* lineage: greater GC shift differences between X and A for NAY than for the other mutation classes (Fig. S7e-h). We employed odds ratios ($GC\_OR_{X,A}$) as summary statistics for the degree of X-vs-A difference in GC fixations. We calculated the ratio of $N_{W\rightarrow S}$ to $N_{S\rightarrow W}$ for X-linked loci (class 1 in Table S10) divided by that for autosomal loci (class 2 in Table S10) for each 2 x 2 table (corresponding to each bin). We compared $GC\_OR_{X,A}$ for NAY with that for the other mutation classes using MWU tests. In *Dmel*, NAY shows greater X effects for AT shifts than the other mutation classes (Table S13). Significant differences in $GC\_OR_{X,A}$ values in pairwise comparison of NAY to SI, $4f_5$, and $2f_{non\text{-}NAY}$ (Table S13) strongly support NAY-specific factor(s) that magnify X-effects favoring NAT codon usage in the ancestral *Dmel* lineage.

**Fig. S7. W→S fixation skews at autosomal and X-linked loci in the *D. simulans* and *D. melanogaster* ancestral lineages**

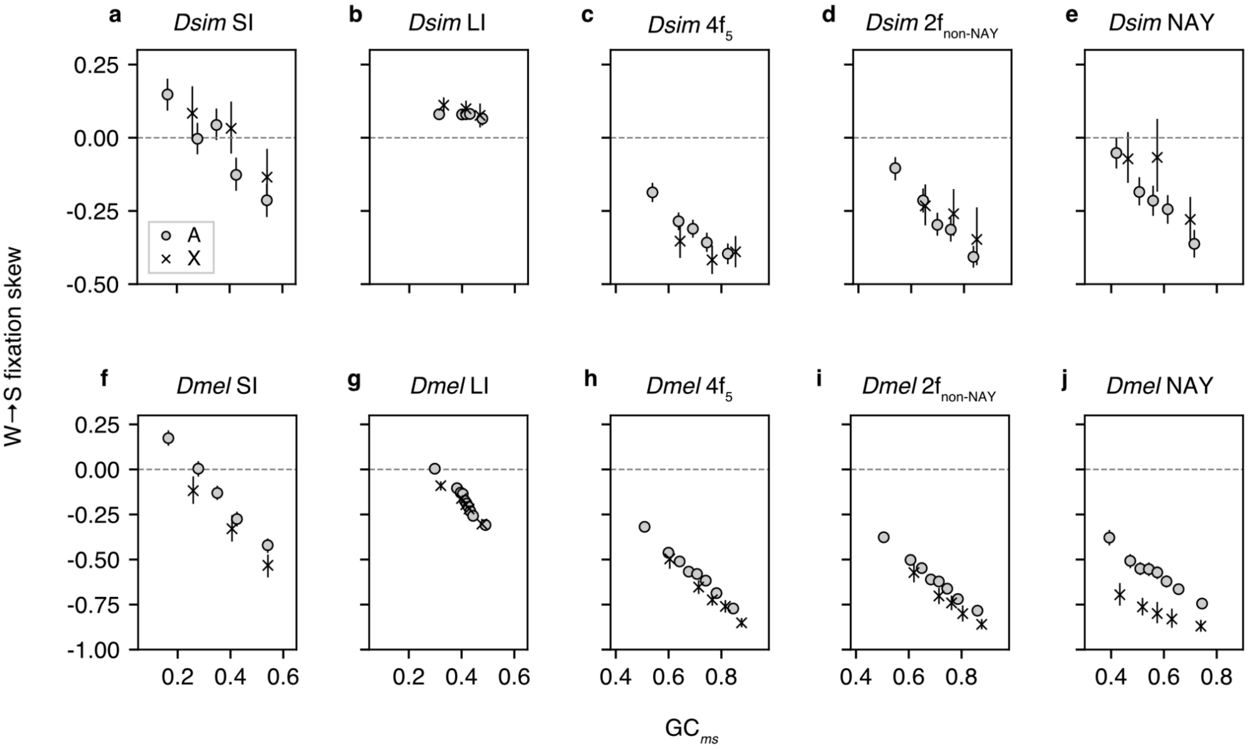

W→S fixation skews at autosomal ("A") and X-linked ("X") loci. A legend in (a) applies to all panels. (a) Short introns (SI) in *D. simulans* (*Dsim*). (b) Long introns (LI) in *Dsim*. (c) 4-fold synonymous families except for Gly ($4f_5$) in *Dsim*. (d) Non-NAY 2-fold synonymous families ($2f_{non\text{-}NAY}$) in *Dsim*. (e) NAY synonymous families in *Dsim*. (f) SI in *D. melanogaster* (*Dmel*). (g) LI in *Dmel*. (h) $4f_5$in *Dmel*. (i) $2f_{non\text{-}NAY}$ in *Dmel*. (j) NAY synonymous families in *Dmel*. CDS are ranked by GC content of sites within intron sites or at synonymous positions at *ms* node ("$GC_{ms}$") and assigned to bins with roughly similar numbers of intron sites or codons. Error bars indicate 95% CIs among 1000 bootstrap replicates. Ancestral reconstructions were sampled in units of introns or CDS for each bin.

**Table S12. Testing X-effects for GC-altering fixations.**

| Species | Mutation class | Chr | Total $N_{W\to S}$ | Total $N_{S\to W}$ | MH $z$ | | WSR | | Bin num |
|---|---|---|---|---|---|---|---|---|---|
| *Dsim* | SI | A | 1409.5 | 1547.9 | | | | | |
| | | X | 381.2 | 383.8 | 1.62 | | 9 | | 9 |
| | LI | A | 33017.6 | 28273.1 | | | | | |
| | | X | 9182.9 | 7558.5 | 2.58 | | 363 | | 46 |
| | $4f_5$ | A | 4237.4 | 8204.4 | | | | | |
| | | X | 1036.2 | 2315.1 | -1.82 | | 71 | | 21 |
| | $2f_{non\text{-}NAY}$ | A | 2706.1 | 4879.7 | | | | | |
| | | X | 723.1 | 1289.6 | 1.19 | | 64 | | 20 |
| | NAY | A | 1748.2 | 2777.7 | | | | | |
| | | X | 567.9 | 733.3 | 3.18 | * | 13 | | 13 |
| *Dmel* | SI | A | 2830.6 | 3980.2 | | | | | |
| | | X | 404.8 | 788.8 | -3.46 | ** | 13 | * | 15 |
| | LI | A | 52159.8 | 72029.8 | | | | | |
| | | X | 9826.0 | 14395.3 | -2.78 | * | 408 | | 51 |
| | $4f_5$ | A | 5930.9 | 22539.4 | | | | | |
| | | X | 776.8 | 4321.6 | -4.22 | *** | 24 | ** | 23 |
| | $2f_{non\text{-}NAY}$ | A | 3838.7 | 16434.0 | | | | | |
| | | X | 521.5 | 3365.9 | -4.51 | *** | 41 | ** | 24 |
| | NAY | A | 2596.3 | 10001.1 | | | | | |
| | | X | 292.9 | 2554.2 | -12.25 | *** | 0 | *** | 15 |

Note: Each bin is set to contain at least 25 sequences (intron or CDS). “SI” and “LI” mean short introns and long introns, respectively. “Chr” indicates chromosomal classes: autosomal (A) and X-linked. The minimum site counts per bin were as follows: 3000, 3000, 7000, 6000, and 6000 for SI, LI, $4f_5$ (4-fold synonymous families except for Gly), $2f_{non\text{-}NAY}$ (2-fold synonymous families excluding NAY codons) and NAY, respectively, in *D. simulans* (*Dsim*), and 2000, 2000, 6000, 5000, 5000 for SI, LI, $4f_5$, $2f_{non\text{-}NAY}$, and NAY, respectively, in *D. melanogaster* (*Dmel*). We filtered bins with the lowest and highest $GC_{ms}$ from analyses of each mutation class because these bins contained sequences with very different $GC_{ms}$. “Bin num” indicates the number of bins employed for statistical tests. Total fixation counts across the $GC_{ms}$ bins are shown as “Total $N_{W\to S}$” and “Total $N_{S\to W}$” for GC-increasing (W→S) and GC-decreasing (S→W) changes, respectively. Multiple tests correction was performed for each species using the sequential Bonferroni method (Holm 1979). *, **, and *** indicate $p$ < 0.05, < 0.01, and < 0.001, respectively.

**Table S13. Comparisons of the magnitude of X-effects for GC fixations for NAY and other mutation classes.**

| Species | Mutation class 1 | Mutation class 2 | Bin num 1 | Bin num 2 | OR mean 1 | OR mean 2 | MWU *z* | |
|---|---|---|---|---|---|---|---|---|
| *Dsim* | SI | NAY | 9 | 13 | 1.18 | 1.26 | 0.27 | |
| | $4f_5$ | NAY | 21 | 13 | 0.97 | 1.26 | 2.87 | * |
| | $2f_{non\text{-}NAY}$ | NAY | 20 | 13 | 1.11 | 1.26 | 1.23 | |
| | Combined | NAY | 54 | 13 | 1.03 | 1.26 | 2.12 | |
| *Dmel* | SI | NAY | 15 | 15 | 0.82 | 0.46 | -4.19 | *** |
| | $4f_5$ | NAY | 23 | 15 | 0.84 | 0.46 | -5.08 | *** |
| | $2f_{non\text{-}NAY}$ | NAY | 24 | 15 | 0.80 | 0.46 | -4.29 | *** |
| | Combined | NAY | 66 | 15 | 0.82 | 0.46 | -5.44 | *** |

Note: Intron and CDS bins are the same as Table S12. OR indicates the odds ratio calculated for GC-altering fixation counts between X-linked and autosomal loci (*i.e.*, the ratio of $N_{W\rightarrow S}$ to $N_{S\rightarrow W}$ for X-linked loci divided that for autosomal loci). The OR values are averaged across $GC_{ms}$ bins. We excluded bins with the lowest and highest $GC_{ms}$ from analyses of each mutation class. The distributions of OR are compared by the MWU test between mutation classes 1 and 2 (NAY). Positive $z$ values indicate that OR for NAY is higher than that for the other mutation class and negative $z$ values indicate the opposite relationship. "SI", "$4f_5$", and "$2f_{non\text{-}NAY}$" mean short introns, 4-fold synonymous families excluding Gly, and 2-fold synonymous families excluding NAY families. "Combined" class includes SI, $4f_5$, and $2f_{non\text{-}NAY}$. Multiple tests were corrected for each species using the sequential Bonferroni method (Holm 1979). *, **, and *** indicate $p < 0.05$, $< 0.01$, and $< 0.001$, respectively.

#### *ApT over ApC dinucleotide preference*

##### Context-dependent W→S aDAF at synonymous sites

Fixation biases for TTR→CTR are markedly shifted toward preference for TTR in the 5′ A context (A|TTR→A|CTR) in *Dmel* (Fig. 5a; Table S14; $p = 0.0043$ from a permutation approach, $10^4$ replicates) and in *Dsim* (Fig. 5b; Table S15; $p = 0.0001$ from a permutation approach, $10^4$ replicates). If the ApT over ApC dinucleotide preference is acting on transcribed and non-transcribed strands, a similar trend in fixation bias estimates is expected for the reverse complement of these changes, A|T→G|T. We employed A↔G synonymous changes at codon 3rd positions for each synonymous family for this test.

In *Dmel*, the direction and magnitude of GC fixation biases are heterogeneous among synonymous families (Fig. S8a) but W→S aDAF skews are consistently lower for A|T→G|T than for A|V→G|V (Fig. S8a; WSR test $p = 0.0007$). *Dsim* shows similar contrasts in W→S aDAF skew with Dmel. W→S aDAF skew is heterogeneous among synonymous families for both A|T→G|T and A|V→G|V changes but is consistently lower for A|T→G|T than for A|V→G|V (Fig. S8b; WSR test $p = 0.0002$). ApT over ApC dinucleotide preference appears to be a prevalent force in both *Dmel* and *Dsim* that can act in opposition to GC fixation biases including MCP and gBGC. Reduced signals for GC fixation biases at *Dmel* synonymous sites do not necessarily imply absence of translational selection.

##### Context-dependent W→S aDAF at SI and LI sites

Consistent with synonymous sites, LI sites show a reduced W→S aDAF skews for ApT→ApC than BpT→BpC changes on the both transcribed (tr) and non-transcribed (rc, reverse complement) strands in both *Dmel* and *Dsim* (Figs. 5c and 5d; Table S16; *Dmel* tr, $p = 0.00007$; *Dmel* rc, $p =$

0.00032; *Dsim* tr, $p = 0.0001$; *Dsim* rc, $p = 0.0001$; $p$ values are from permutation approaches with $10^5$ replicates). Polymorphism data from SI sites are limited for *Dmel*, but we could assess W→S aDAF skew differences at SI sites in *Dsim*. Similarly to LI sites, SI sites show a reduced W→S aDAF skew for ApT→ApC than for BpT→BpC on both transcribed and non-transcribed strands (Table S16; *Dsim* tr, $p = 0.00038$; *Dsim* rc, $p = 0.0001$; $p$ values are from the permutation approach with $10^5$ replicates).

We took advantage of large polymorphism counts from LI sites to test for ApT vs ApC dinucleotide preference at X-linked loci. In both *Dmel* and *Dsim*, LI shows significantly lower W→S aDAF skew for ApT→ApC than BpT→BpC on both transcribed and untranscribed strands (Table S17; *Dmel* tr, $p = 0.013$; *Dmel* rc, $p = 0.0044$; *Dsim* tr, $p = 0.00005$; *Dsim* rc, $p = 0.0001$; $p$ values are from the permutation approach with $10^5$ replicates).

**Fig. S8. ApT vs GpT dinucleotide preference in coding regions: inference from synonymous polymorphisms in *D. melanogaster* and *D. simulans*.**

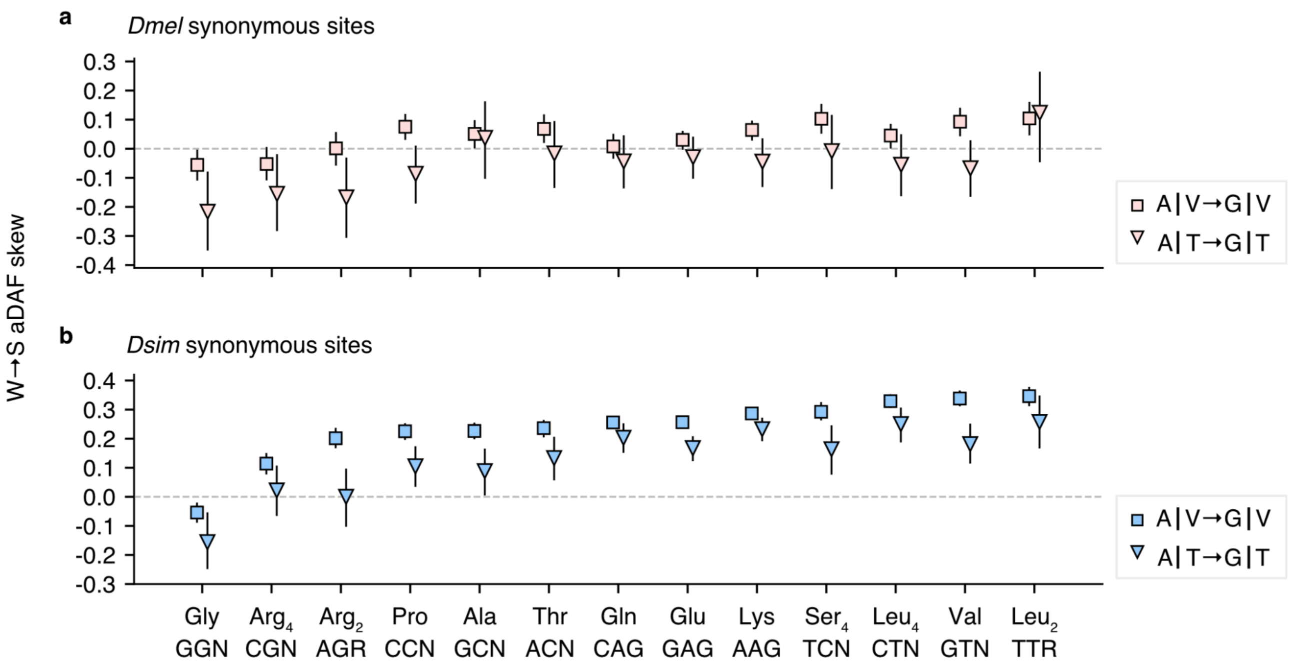

W→S aDAF skew measures SFS difference between "forward" and "reverse" dinucleotide changes. Arrows indicate the forward direction and reverse refers to the opposite direction. Positive W→S aDAF skew values indicate higher frequencies of forward changes than reverse changes. We show results for ApT→GpT and ApV→GpV dinucleotides where "V" indicates C, A, or G nucleotides. These are the reverse complement cases for the results in Fig. 5a and b. W→S aDAF skews are examined within synonymous families for (a) *D. melanogaster* (*Dmel*) polymorphisms and (b) *D. simulans* (*Dsim*) polymorphisms. Arginine-coding and leucine-coding codons are split into 2-fold ($Arg_2$: AGA and AGR, $Leu_2$: TTA and TTG) and 4-fold ($Arg_4$: CGT, CGC, CGA, and CGG, $Leu_4$: CTT, CTC, CTA, and CTG) synonymous families. Synonymous changes at codon 3rd positions in autosomal loci are examined. Pipes ("|") indicate codon boundaries. Synonymous families are arranged in order of W→S aDAF skew values for A|V↔G|V changes in *Dsim*. Error bars indicate 95% CIs among 1000 bootstrap replicates. Ancestral reconstructions are resampled in units of CDS. WSR tests are employed to test consistent differences in W→S aDAF skews for A|T→G|T compared to A|V→G|V ($p$ = 0.00073 for *Dmel* and $p$ = 0.00024 for *Dsim*). See Tables S14 and S15 for details of statistical tests in *Dmel* and *Dsim*, respectively.

**Table S14. ApT vs GpT preference at synonymous sites in autosomal loci in *D. melanogaster*.**

| Syn [a] | Change [b] | # Forward [c] | # Reverse [c] | *z* [d] | | W→S aDAF skew [e] | W→S aDAF skew diff [f] |
|---|---|---|---|---|---|---|---|
| Leu | A\|T→A\|C | 109.2 | 626.8 | -2.51 | | -0.144 | |
| | B\|T→B\|C | 427.6 | 3081.4 | 1.82 | | 0.034 | -0.178 |
| Gly | A\|T→G\|T | 88.4 | 48.6 | -2.87 | | -0.217 | |
| | A\|V→G\|V | 511.7 | 532.3 | -2.30 | | -0.056 | -0.161 |
| Arg$_4$ | A\|T→G\|T | 65.2 | 124.8 | -2.43 | | -0.155 | |
| | A\|V→G\|V | 261.2 | 1022.8 | -2.39 | | -0.053 | -0.103 |
| Arg$_2$ | A\|T→G\|T | 43.3 | 92.7 | -1.86 | | -0.168 | |
| | A\|V→G\|V | 205.9 | 912.1 | -0.30 | | 0.001 | -0.169 |
| Pro | A\|T→G\|T | 99.7 | 185.3 | -2.05 | | -0.087 | |
| | A\|V→G\|V | 424.5 | 1426.5 | 2.38 | | 0.075 | -0.162 |
| Ala | A\|T→G\|T | 53.4 | 134.6 | 0.54 | | 0.037 | |
| | A\|V→G\|V | 341.7 | 1222.3 | 2.04 | | 0.051 | -0.014 |
| Thr | A\|T→G\|T | 65.6 | 176.4 | 0.16 | | -0.016 | |
| | A\|V→G\|V | 291.2 | 1425.8 | 2.15 | | 0.068 | -0.084 |
| Gln | A\|T→G\|T | 92.8 | 565.2 | -0.87 | | -0.044 | |
| | A\|V→G\|V | 412.8 | 3072.2 | -0.37 | | 0.008 | -0.051 |
| Glu | A\|T→G\|T | 137.8 | 712.2 | -0.65 | | -0.029 | |
| | A\|V→G\|V | 645.2 | 4083.8 | 1.29 | | 0.030 | -0.059 |
| Lys | A\|T→G\|T | 145.0 | 702.0 | -0.72 | | -0.045 | |
| | A\|V→G\|V | 605.9 | 3469.1 | 3.45 | * | 0.064 | -0.109 |
| Ser$_4$ | A\|T→G\|T | 62.5 | 170.5 | 0.21 | | -0.009 | |
| | A\|V→G\|V | 294.5 | 1372.5 | 3.49 | * | 0.103 | -0.111 |
| Leu$_4$ | A\|T→G\|T | 75.4 | 429.6 | -1.24 | | -0.055 | |
| | A\|V→G\|V | 337.2 | 3386.8 | 1.21 | | 0.045 | -0.100 |
| Val | A\|T→G\|T | 80.9 | 288.1 | -1.29 | | -0.067 | |
| | A\|V→G\|V | 233.0 | 2394.0 | 3.28 | * | 0.093 | -0.160 |
| Leu$_2$ | A\|T→G\|T | 25.5 | 159.5 | 1.58 | | 0.123 | |
| | A\|V→G\|V | 194.7 | 1369.3 | 2.61 | | 0.104 | 0.019 |

(See the next page for footnotes.)

[a] Synonymous families. “Leu” in the top row refers to YTR codons where ambiguity characters “Y” and “R” indicate T or C and A or G, respectively. SFS for synonymous changes at 1st positons (*i.e.*, TTA↔CTA and TTG↔CTG) are examined. For other synonymous families, SFS for changes at the 3rd positions are examined. Subscript digits indicate coding redundancy at codon 3rd positions.

[b] Dinucleotide change in the “forward” direction are indicated by arrows. “Reverse” refers to the opposite direction. Pipes “|” indicate codon boundaries. Pooled dinucleotide changes are indicated by ambiguity characters (“B” for T, C, or G and “V” for C, A, or G).

[c] Numbers of polymorphic changes.

[d] MWU test statistic for SFS comparisons between forward and reverse changes. Positive $z$ values indicate higher allele frequencies for forward changes compared to reverse changes. Multiple test corrections were conducted using the sequential Bonferroni method (Holm 1979). *, ** and *** indicate $p < 0.05$, $< 0.01$ and $< 0.001$, respectively.

[e] W→S aDAF skew = $(f - r) / (f + r)$, where $f$ is W→S aDAF and $r$ is S→W aDAF. This is an index of the direction and magnitude of a SFS difference between forward (W→S) and reverse (S→W) changes. Positive values indicate higher aDAF for W→S change and negative values indicate lower aDAF for W→S change.

[f] Difference in W→S aDAF skew between the upper and lower dinucleotide change classes for each synonymous family. The null hypothesis of no difference in W→S aDAF skew is tested using a permutation approach for Leu (see text) and WSR tests for other synonymous families. $P$ values from the permutation approach and WSR tests are 0.0043 and 0.00073, respectively. These $p$ values are combined using Fisher’s method (1954): $\chi^2 = 25.3$ and $p = 0.000043$.

**Table S15. ApT vs GpT preference at synonymous sites in autosomal loci in *D. simulans*.**

| Syn [a] | Change [b] | # Forward [c] | # Reverse [c] | *z* [d] | | W→S aDAF skew [e] | W→S aDAF skew diff [f] |
|---|---|---|---|---|---|---|---|
| Leu | A\|T→A\|C | 734.8 | 1518.2 | 5.44 | *** | 0.126 | |
| | B\|T→B\|C | 2563.0 | 9128.0 | 19.53 | *** | 0.251 | -0.125 |
| Gly | A\|T→G\|T | 480.7 | 159.3 | -3.23 | ** | -0.156 | |
| | A\|V→G\|V | 2308.8 | 1734.2 | -4.71 | *** | -0.054 | -0.102 |
| $Arg_4$ | A\|T→G\|T | 327.0 | 275.0 | 0.84 | | 0.022 | |
| | A\|V→G\|V | 1317.9 | 2930.1 | 5.51 | *** | 0.114 | -0.092 |
| $Arg_2$ | A\|T→G\|T | 221.0 | 273.0 | -0.82 | | 0.000 | |
| | A\|V→G\|V | 1125.2 | 3099.8 | 9.04 | *** | 0.201 | -0.201 |
| Pro | A\|T→G\|T | 394.9 | 613.1 | 3.16 | ** | 0.105 | |
| | A\|V→G\|V | 2063.1 | 5127.9 | 12.71 | *** | 0.225 | -0.121 |
| Ala | A\|T→G\|T | 283.6 | 356.4 | 0.82 | | 0.088 | |
| | A\|V→G\|V | 1578.8 | 4291.2 | 11.56 | *** | 0.226 | -0.138 |
| Thr | A\|T→G\|T | 307.5 | 461.5 | 3.05 | ** | 0.133 | |
| | A\|V→G\|V | 1329.0 | 4398.0 | 12.09 | *** | 0.236 | -0.103 |
| Gln | A\|T→G\|T | 614.0 | 1649.0 | 6.06 | *** | 0.203 | |
| | A\|V→G\|V | 2310.3 | 10362.7 | 17.97 | *** | 0.256 | -0.053 |
| Glu | A\|T→G\|T | 871.6 | 2034.4 | 6.61 | *** | 0.167 | |
| | A\|V→G\|V | 3736.9 | 14209.1 | 23.85 | *** | 0.257 | -0.089 |
| Lys | A\|T→G\|T | 872.8 | 1973.2 | 9.91 | *** | 0.232 | |
| | A\|V→G\|V | 3713.1 | 12179.9 | 26.99 | *** | 0.287 | -0.054 |
| $Ser_4$ | A\|T→G\|T | 209.9 | 506.1 | 3.30 | ** | 0.164 | |
| | A\|V→G\|V | 1131.5 | 4404.5 | 13.70 | *** | 0.292 | -0.129 |
| $Leu_4$ | A\|T→G\|T | 368.8 | 1012.2 | 6.68 | *** | 0.250 | |
| | A\|V→G\|V | 1715.2 | 10004.8 | 21.30 | *** | 0.329 | -0.079 |
| Val | A\|T→G\|T | 339.2 | 818.8 | 4.18 | *** | 0.181 | |
| | A\|V→G\|V | 1136.6 | 7487.4 | 18.32 | *** | 0.338 | -0.157 |
| $Leu_2$ | A\|T→G\|T | 158.6 | 468.4 | 4.77 | *** | 0.258 | |
| | A\|V→G\|V | 865.2 | 4335.8 | 16.09 | *** | 0.346 | -0.088 |

(See the next page for footnotes.)

[a] Synonymous families.
[b] Dinucleotide change in the “forward” direction are indicated by arrows. “Reverse” refers to the the opposite direction. Pipes “|” indicate codon boundaries. Pooled dinucleotide changes are indicated by ambiguity characters (“B” for T, C, or G and “V” for C, A, or G).
[c] Numbers of polymorphic changes.
[d] MWU test statistic for SFS comparisons between forward and reverse changes. Positive *z* values indicate higher allele frequencies for forward changes compared to reverse changes. Multiple test corrections were conducted using the sequential Bonferroni method (Holm 1979). *, ** and *** indicate $p < 0.05$, $< 0.01$ and $< 0.001$, respectively.
[e] W→S aDAF skew = $(f - r) / (f + r)$, where *f* is W→S aDAF and *r* is S→W aDAF. This is an index of the direction and magnitude of a SFS difference between forward (W→S) and reverse (S→W) changes. Positive values indicate higher aDAF for W→S change and negative values indicate lower aDAF for W→S change.
[f] Difference in W→S aDAF skew between the upper and lower dinucleotide change classes for each synonymous family. The null hypothesis of no difference in W→S aDAF skew is tested using a permutation approach for Leu (see text) and WSR tests for other synonymous families. *P* values from the permutation approach and WSR tests are 0.0001 and 0.00024, respectively. These *p* values are combined using Fisher’s method (1954): $\chi^2 = 35.1$ and $p = 0.00000045$.

**Table S16. ApT vs ApC dinucleotide preference at autosomal SI and LI in *D. melanogaster* and *D. simulans*.**

| Species [a] | Length [b] | Strand [c] | Change [d] | # Forward [e] | # Reverse [e] | *z* [f] | | W→S aDAF skew [g] | W→S aDAF skew diff [h] | |
|---|---|---|---|---|---|---|---|---|---|---|
| *Dmel* | SI | tr | ApT→ApC | 187.4 | 195.6 | -0.83 | | -0.015 | - | |
| | | | BpT→BpC | 291.3 | 516.7 | 1.14 | | 0.063 | -0.08 | |
| | | rc | ApT→GpT | 201.4 | 206.6 | -1.14 | | -0.051 | - | |
| | | | ApV→GpV | 335.9 | 592.1 | 1.49 | | 0.049 | -0.10 | * |
| | LI | tr | ApT→ApC | 3585.8 | 5381.2 | -2.32 | | -0.034 | - | |
| | | | BpT→BpC | 10110.3 | 21379.7 | 1.63 | | 0.010 | -0.04 | *** |
| | | rc | ApT→GpT | 3865.8 | 5627.2 | -1.05 | | -0.016 | - | |
| | | | ApV→GpV | 10104.2 | 20888.8 | 3.78 | ** | 0.023 | -0.04 | *** |
| *Dsim* | SI | tr | ApT→ApC | 964.0 | 663.0 | 1.55 | | 0.016 | - | |
| | | | BpT→BpC | 1638.8 | 2179.2 | 6.64 | *** | 0.129 | -0.11 | *** |
| | | rc | ApT→GpT | 1116.8 | 686.2 | -0.93 | | -0.074 | - | |
| | | | ApV→GpV | 1769.6 | 2207.4 | 4.29 | *** | 0.067 | -0.14 | *** |
| | LI | tr | ApT→ApC | 14526.5 | 12510.5 | 13.35 | *** | 0.077 | - | |
| | | | BpT→BpC | 40020.0 | 56359.0 | 41.37 | *** | 0.184 | -0.11 | *** |
| | | rc | ApT→GpT | 15906.1 | 13249.9 | 13.80 | *** | 0.075 | - | |
| | | | ApV→GpV | 40998.9 | 56759.1 | 46.08 | *** | 0.194 | -0.12 | *** |

[a] "*Dmel*" and "*Dsim*" indicate *D. melanogaster* and *D. simulans*, respectively.

[b] Intron length classes. "SI" and "LI" indicate short introns and long introns, respectively.

[c] Strand class indicating whether ApT→ApC change is on the transcribed (tr) strand or the non-transcribed strand (rc, reverse complement).

[d] Dinucleotide change in the "forward" direction are indicated by arrows. "Reverse" refers to the the opposite direction. Pooled dinucleotide changes are indicated using ambiguity characters ("B" for T, C, or G and "V" for C, A, or G).

[e] Numbers of polymorphic changes.

[f] MWU test statistics for SFS comparisons between W→S and S→W changes. The sequential Bonferroni method (Holm 1979) was employed in multiple test corrections within each species. ** and *** indicate $p < 0.01$ and $< 0.001$, respectively.

[g] W→S aDAF skew = $(f - r) / (f + r)$, where $f$ is W→S aDAF and $r$ is S→W aDAF. This is an index of the direction and magnitude of a SFS difference between forward (W→S) and reverse (S→W) changes. Positive values indicate higher aDAF for W→S change and negative values indicate lower aDAF for W→S change.

[h] Difference in W→S aDAF skew between the upper and lower dinucleotide change classes for each strand class. *P* values are estimated using a permutation approach. The sequential Bonferroni method is employed in the multiple test corrections within each species. *** indicates $p < 0.001$.

**Table S17. ApT vs ApC dinucleotide preference at X-linked SI and LI in *D. melanogaster* and *D. simulans*.**

| Species [a] | Length [b] | Strand [c] | Change [d] | # Forward [e] | # Reverse [e] | *z* [f] | | W→S aDAF skew [g] | W→S aDAF skew diff [h] | |
|---|---|---|---|---|---|---|---|---|---|---|
| *Dmel* | SI | tr | ApT→ApC | 30.3 | 42.7 | -2.70 | | -0.279 | - | |
| | | | BpT→BpC | 63.0 | 155.0 | -0.50 | | -0.068 | -0.211 | |
| | | rc | ApT→GpT | 44.2 | 39.8 | 0.03 | | -0.068 | - | |
| | | | ApV→GpV | 61.4 | 126.6 | 0.92 | | 0.029 | -0.097 | |
| | LI | tr | ApT→ApC | 613.4 | 1000.6 | -0.56 | | -0.021 | - | |
| | | | BpT→BpC | 1834.3 | 3838.7 | 2.50 | | 0.048 | -0.068 | * |
| | | rc | ApT→GpT | 747.8 | 1068.2 | -1.66 | | -0.024 | - | |
| | | | ApV→GpV | 1861.6 | 3654.4 | 2.94 | * | 0.052 | -0.075 | * |
| *Dsim* | SI | tr | ApT→ApC | 111.7 | 66.3 | -0.12 | | 0.059 | - | |
| | | | BpT→BpC | 224.1 | 314.9 | 2.21 | | 0.137 | -0.078 | |
| | | rc | ApT→GpT | 146.4 | 88.6 | 0.89 | | 0.079 | - | |
| | | | ApV→GpV | 254.5 | 266.5 | 4.33 | *** | 0.211 | -0.132 | |
| | LI | tr | ApT→ApC | 2139.4 | 1564.6 | 7.30 | *** | 0.140 | - | |
| | | | BpT→BpC | 5781.9 | 6929.1 | 20.47 | *** | 0.229 | -0.089 | *** |
| | | rc | ApT→GpT | 2405.2 | 1693.8 | 7.14 | *** | 0.109 | - | |
| | | | ApV→GpV | 6059.3 | 6939.7 | 21.98 | *** | 0.256 | -0.148 | *** |

[a] "*Dmel*" and "*Dsim*" indicate *D. melanogaster* and *D. simulans*, respectively.

[b] Intron length classes. "SI" and "LI" indicate short introns and long introns, respectively.

[c] Strand class indicating whether ApT→ApC change is on the transcribed (tr) strand or on the non-transcribed strand (rc, reverse complement).

[d] Dinucleotide change in the "forward" direction are indicated by arrows. "Reverse" refers to the the opposite direction. Pooled dinucleotide changes are indicated using ambiguity characters ("B" for T, C, or G and "V" for C, A, or G).

[e] Numbers of polymorphic changes.

[f] MWU test statistics for SFS comparisons between W→S and S→W changes. The sequential Bonferroni method (Holm 1979) was employed in multiple test corrections within each species. * and *** indicate $p < 0.05$ and $< 0.001$, respectively.

[g] W→S aDAF = $(f - r) / (f + r)$, where $f$ is W→S aDAF and $r$ is S→W aDAF. This is an index of the direction and magnitude of a SFS difference between forward (W→S) and reverse (S→W) changes. Positive values indicate higher aDAF for W→S change and negative values indicate lower aDAF for W→S change.

[h] Difference in W→S aDAF skew between the upper and lower dinucleotide change classes for each strand class. The null hypothesis of no difference in W→S aDAF skew is tested using a permutation approach. The sequential Bonferroni method is employed in multiple test corrections within each species. * and *** indicates $p < 0.05$ and 0.001.

**Context-dependent W→S fixation skews at synonymous sites**

Our fixation analysis above revealed accelerated NAC→NAT fixations in the ancestral *Dmel* lineage (Figs. 4c and 4d; Table S11). We examined whether ApT over ApC dinucleotide preference can explain this pattern. We employed W→S fixation skew statistics to assess the degree and direction of GC content change. To distinguish codon and dinucleotide preference, we compared W→S fixation skews for A|TTR→A|CTR vs B|TTR→B|CTR. *Dmel* has undergone accelerated ApC→ApT fixations. A|T→A|C changes show marginally significant differences toward lower W→S fixation skew than B|T→B|C according to WSR test (Fig. S9b; Table S18; MH test $p = 0.11$; WSR test $p = 0.016$) but A|T→G|T changes show strong support for more negative W→S fixation skew than A|V→G|V (Fig. S9d; Table S18; MH test $p < 10^{-10}$; WSR test $p < 10^{-5}$). These patterns strongly support ApT over ApC dinucleotide preference contributing to AT shifts at synonymous sites in the ancestral *Dmel* lineage. Such shifts are expected to be greater within NAY synonymous families where all S→W synonymous changes are also ApT to ApC dinucleotide changes.

In contrast, *Dsim* does not show support for accelerated fixations for A|C→A|T or G|T→A|T. W→S fixation skews do not differ for A|C→A|T vs B|C→B|T (Fig. S9a; Table S18; MH test $p = 0.39$, WSR test $p = 0.25$). Although W→S fixation skew values for A|T→G|T fixations at codon 3rd positions are generally more negative compared to skews for A|V→G|V in *Dsim* (Fig. S9c), these differences in W→S fixation skew are not statistically significant (Table S18; MH test $p = 0.58$; WSR test $p = 0.80$). Similarly to the *Dmel* fixation analysis, we combined $p$ values from the A|T→A|C vs B|T→B|C and A|T→G|T vs A|V→G|V tests and found no support for a significant difference in W→S fixation biases ($p = 0.63$). Limited counts for fixations in the *Dsim* lineage, especially for A|T→A|C changes (Table S18), likely limit statistical power to detect ApT vs ApC dinucleotide preference.

Dinucleotide preferences could underlie excess NAT fixations at NAY codons in the *D. melanogaster* ancestral lineage as well as X effects for such fixations. X effects for accelerated AT shifts for other mutation classes may reflect dinucleotide pressure. However, more specified mutation matrices may be necessary to distinguish among the causes of these fixation patterns.

**Fig. S9. A|T→A|C fixation skews for synonymous changes in the ancestral *D. melanogaster* and *D. simulans* lineages.**

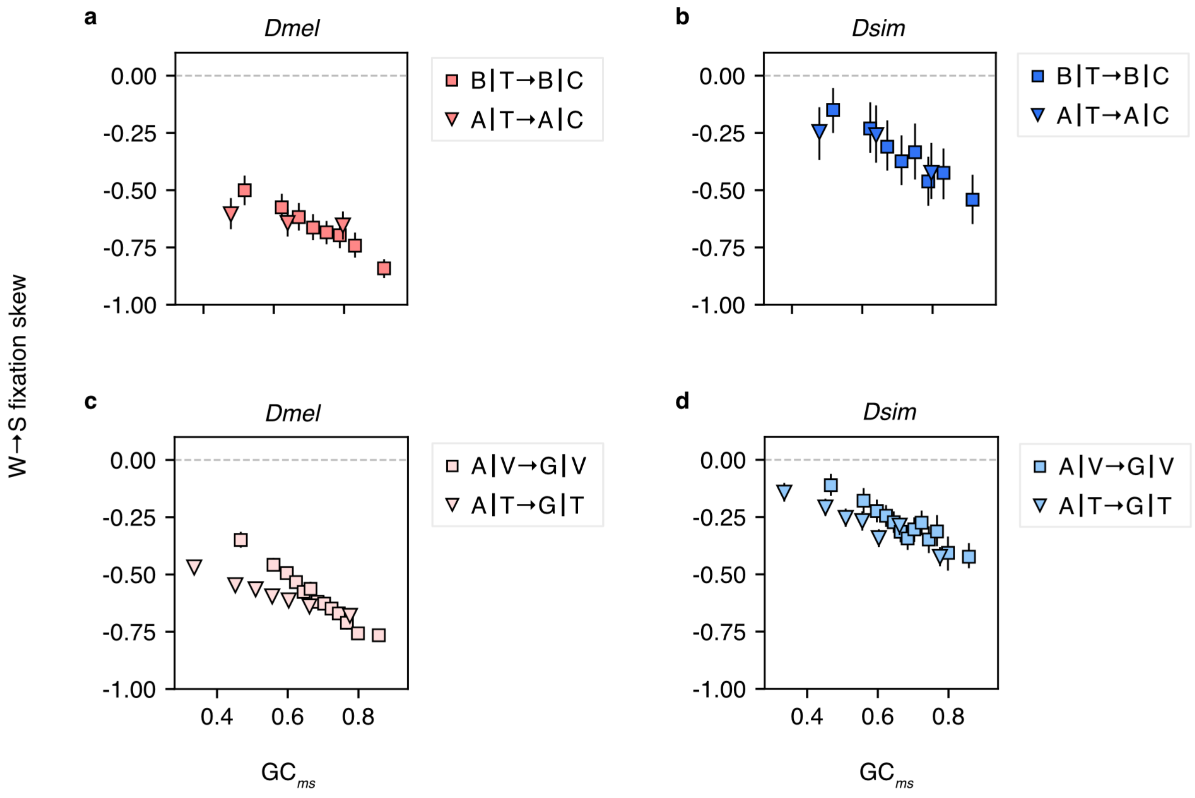

W→S fixation skew = ($N_{W→S}$ – $N_{S→W}$) / ($N_{W→S}$ + $N_{S→W}$), where $N_{W→S}$ is W→S fixation count and $N_{S→W}$ is S→W fixation count. This is an index of the direction and magnitude of the departure from GC content equilibrium. X-axis values indicate GC content at *ms* node, $GC_{ms}$, the proportion of “S” state (A|C, B|C, G|T, and G|V dinucleotides) among S + “W” states (A|Y, B|Y, R|T, and R|V dinucleotides) for A|T→A|C, B|T→B|C, A|T→G|T and A|V→G|V changes, respectively. CDS are ranked by $GC_{ms}$ and assigned to bins with similar numbers of codons. Autosomal loci are employed. Fixations are inferred changes within internal branches: (a) and (c) *ms-m'* for *D. melanogaster* (*Dmel*) fixation data and (b) and (d) *ms-s*' for *D. simulans* (*Dsim*) fixation data. (a) and (b) show A|T→A|C vs B|T→B|C changes at Leu codon 1st positions. (c) and (d) show A|T→G|T vs A|V→G|V at codon 3rd positions. Arrows indicate “forward” direction (W→S) and “reverse” indicates the opposite direction. GGA→GGG changes are excluded because SFS analyses support GGA codon preference over GGG in both *Dsim* and *Dmel*. All other A↔G synonymous changes at codon 3rd positions are included. Error bars indicate 95% CIs among 1000 bootstrap replicates. Ancestral reconstructions are resampled in units of CDS. Statistical analyses are presented in Table S18.

**Table S18. A|T→A|C fixation skews for synonymous changes in the ancestral *D. melanogaster* and *D. simulans* lineages.**

| Species | Dinuc change [a] | Total $N_{W\to S}$ [b] | Total $N_{S\to W}$ [b] | MH $z$ [c] | | WSR [d] | | Bin num [e] |
|---|---|---|---|---|---|---|---|---|
| *Dmel* | B\|T→B\|C | 388.1 | 1549.6 | | | | | |
| | A\|T→A\|C | 36.6 | 170.4 | -1.61 | | 0 | * | 7 |
| | A\|V→G\|V | 4843.6 | 19051.8 | | | | | |
| | A\|T→G\|T | 771.6 | 3125.9 | -6.99 | *** | 14 | *** | 29 |
| *Dsim* | B\|T→B\|C | 208.8 | 335.0 | | | | | |
| | A\|T→A\|C | 18.7 | 35.1 | -0.85 | | 0 | | 3 |
| | A\|V→G\|V | 3283.8 | 5686.0 | | | | | |
| | A\|T→G\|T | 529.4 | 833.4 | -0.55 | | 55 | | 15 |

[a] Context-dependent nucleotide changes. The "forward" direction is indicated by arrows. A|T→A|C and B|T→B|C are synonymous fixations at Leu codon 1st positions and A|T→G|T and A|V→G|V are synonymous fixations at codon 3rd positions. Pipes "|" indicate codon boundaries. GGA→GGG and GGG→GGA changes are excluded. Pooled context sites are indicated by ambiguity characters ("B" for T, C, or G and "V" for C, A, or G). We tested the null hypothesis of independence in a 2 x 2 table shown in Table S10r, where class 1 is B or V context class and class 2 is A or T context class for N|T↔N|C and A|N↔G|N changes, respectively. GC content at *ms* node ($GC_{ms}$) was calculated for each context class for each CDS and employed for binning. The minimum site counts per bin were as follows: 1000 for B|TTR→B|CTR vs A|TTR→A|CTR, 20000 for NNA|V→NNG|V vs NNA|T→NNG|T in *Dsim*, 10000 for B|TTR→B|CTR vs A|TTR→A|CTR, 1100 for NNA|V→NNG|V vs NNA|T→NNG|T in *Dmel*.

[b] Fixation counts employed for statistical tests. Fixation counts are summed across $GC_{ms}$ bins for W→S and S→W. Autosomal loci are employed.

[c] Mantel-Haenszel (MH) test statistic, $z$, to assess overall associations. Positive MH $z$ values indicate a high W→S fixation skew for a mutation class at the lower row compared to the other mutation class. Negative MH $z$ values indicate a W→S fixation skew difference in the opposite direction. Each cell of 2 x 2 tables included an expected frequency >= 3.5 (no table was excluded). We excluded bins with the lowest and highest $GC_{ms}$ because mean $GC_{ms}$ can vary considerably between mutation classes in these bins. *** indicates statistical significance at $p$ = 0.001 after multiple test corrections.

[d] Wilcoxon's signed-rank (WSR) test statistic. Multiple test corrections were conducted across MH tests and WSR tests for each species using the sequential Bonferroni method (Holm 1979). * and *** indicate statistical significance at $p$ = 0.05 and 0.001, respectively.

[e] The number of bins employed for statistical tests.

#### *Codon compositional trends among distantly related Drosophila: no indications of major codon transitions among 2-fold synonymous families*

Previous studies proposed NAT codon preference in species outside of the *Dmel* subgroup (Powell *et al.* 2003). We examined whether NAT codon preference is detectable by codon compositional trend. Under MCP, the fitness benefits of major codons increase with the number of translation events at a given codon. Because this number is shared among codons within a gene, putative major codons within each synonymous family can be identified as those with elevated representation in genes under stronger translational selection (*i.e.*, those that have strong codon usage bias and/or that show high estimates of translation rate or associated measures such as transcript abundance). However, potential contributions of mutational variation (*e.g.*, transcription associated mutation) and/or fixation biases other than translational selection (*e.g.*, gBGC) to such trends need to be considered carefully (e.g., Kanaya *et al.* 1999; Wright *et al.* 2004; Fig. S3).

Compositional trend analyses have consistently supported G- and C-ending codon preference within $2f_{non\text{-}NAY}$ synonymous families in a wide range of Drosophila species (Akashi 1997; Heger and Ponting 2007b; Vicario *et al.* 2007) but have yielded heterogeneous and, in some cases, ambiguous results for major codons in NAY families. We examined support for NAT preference in four distantly related Drosophila species, *D. willistoni* (*Dwil*), *D. mojavensis* (*Dmoj*), *D. grimshawi* (*Dgri*), and *D. virilis* (*Dvir*), for which major codon shifts within NAY families have been reported (Heger and Ponting 2007b; Vicario *et al.* 2007; Zaborske *et al.* 2014). We also include *D. pseudoobscura* as a distant relative of *Dmel* with strong support for MCP from population genetic studies (Akashi 1997; Kliman 2014; Fuller *et al.* 2014). We employed G/C preference at $2f_{non\text{-}NAY}$ codons as a predictor of intensity for translational selection; this notion is supported by elevated usage of G- or C-ending codons at putatively highly expressed genes (*i.e.*, *Dmel* high transcript abundance genes and their

1-to-1 OrthoFinder orthologs; Fig. S18). NAC codon usage (*i.e.*, $CUB_{Chi/L}$ NAY; see Supplementary Methods) increases as a function of this proxy for the strength of MCP (Fig. S10). The associations differ considerably among genomes with steep slopes in *Dmel*, *Dpse*, and *Dmoj* and more moderate slopes in *Dgri*, *Dwil*, and *Dvir*. However, all correlations are statistically significant and we did not find any support for NAT preference in these analyses of pooled synonymous families (Fig. S10). We found similar patterns for individual synonymous families (Figs. S11-S16).

If GC-increasing forces that are unrelated to MCP (*e.g.*, mutation bias and gBGC) are associated with the $CUB_{Chi/L}$ $2f_{non\text{-}NAY}$, such forces may hide the pattern of elevated NAT codon usage. In fact, we found that SI GC content is correlated with (Fig. S3). To control for variation of the "background" GC content among groups of genes, we use base counts from SI within a set ("bin") of genes to calculate expected values and refer to the resulting statistic as "$CUB_{Chi/L\_binexp}$". Within each bin, we summed codon frequencies across CDS and site counts across introns, respectively.

Distributions of $CUB_{Chi/L\_binexp}$ are narrower than that of $CUB_{Chi/L}$ (Fig. S10). This pattern reflects the positive correlation between $CUB_{Chi/L}$ $2f_{non\text{-}NAY}$ and SI GC (Fig. S3). However, Fig. S10 shows that GC-biasing forces that may be shared between intron and synonymous changes are not sufficient to explain the variation of $CUB_{Chi/L\_binexp}$ $2f_{non\text{-}NAY}$. In addition, all data points show positive $CUB_{Chi/L\_binexp}$ $2f_{non\text{-}NAY}$ values. GC-favoring selective forces in coding regions may reflect fitness benefits of GC-ending codons for translation efficiency and/or other functions such as mRNA stability and co-translational protein folding (reviewed in Hanson and Coller 2018; Liu *et al.* 2021). The basic pattern of elevated $CUB_{Chi/L}$ in *Dmel* highly expressed genes and their 1-to-1 OrthoFinder orthologs (Fig. S18) do not distinguish among such proposed functions.

We did not find support for NAT major codons in any of the examined Drosophila species. $CUB_{Chi/L}$ NAY is elevated in highly expressed genes and their 1-to-1 OrthoFinder orthologs in all examined species (Fig. S19). $CUB_{Chi/L\_binexp}$ NAY shows a similar trend, suggesting that intron GC variation among genes does not explain elevated NAC usage in candidates for genes under stronger MCP. Previous studies suggested NAT preference in *Drosophila willistoni* (Anderson *et al.* 1993; Powell *et al.* 2003; Vicario *et al.* 2007), but used codon usage statistics such as relative synonymous codon usage, "RSCU" (Sharp and Li 1987), codon adaptation index, "CAI" (Sharp and Li 1987), and effective number of codons, "ENC" (Wright 1990) that can give misleading results depending on background base composition (RSCU, ENC) and/or the choice of reference genes (CAI). *Dwil* codon bias ($CUB_{Chi/L}$) is weakly, but statistically significantly, positively correlated between $2f_{non\text{-}NAY}$ and NAY (Figs. S10, S13 and S17). Genes with low $CUB_{Chi/L}$ $2f_{non\text{-}NAY}$ show negative values of $CUB_{Chi/L}$ NAY (Fig. S10), indicating that NAC usage is lower than the overall SI GC content. However, the difference between SI GC content and NAC usage became negligible when we employed bin-specific SI GC content to calculate $CUB_{Chi/L}$ (Fig. S10). Associations of $CUB_{Chi/L\_binexp}$ between $2f_{non\text{-}NAY}$ and each of the NAY synonymous families are not statistically significant (Figs. S13 and S17). Evolutionary forces shared between intron and NAY, such as mutation bias and GC-biased gene conversion, appear to be sufficient to explain $CUB_{Chi/L}$ NAY; these analyses fail to detect any signal of translational selection at NAY codons in *Dwil*.

**Fig. S10. Sensitivity of codon compositional trends to intron GC scaling.**

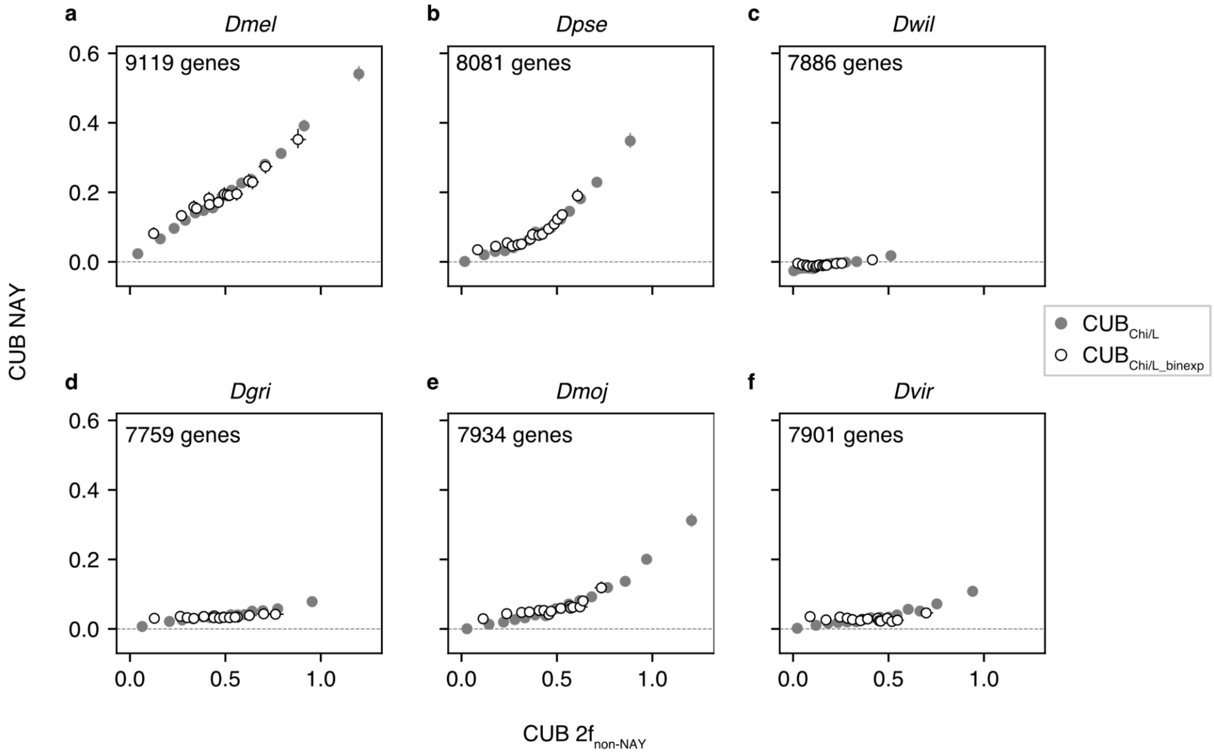

For calculations of the $CUB_{Chi/L}$ statistic, we employed GC content for short intron (SI) sites pooled across the genes in a given analysis. Here, we compare this statistic to $CUB_{Chi/L\_binexp}$ where we employ GC content for bin-specific SI. Positive $CUB_{Chi/L}$ and $CUB_{Chi/L\_binexp}$ values indicate that G-ending or C-ending codon usage is higher than the expected GC content. The statistics are compared between NAY and $2f_{non\text{-}NAY}$ synonymous families for distantly related Drosophila species: (a) *D. melanogaster*, (b) *D. pseudoobscura*, (c) *D. willistoni*, (d) *D. grimshawi*, (e) *D. mojavensis*, (f) *D. virilis*. $CUB_{Chi/L}$ (gray filled) and $CUB_{Chi/L\_binexp}$ (open). Autosomal loci are used. Genes are ranked by $CUB_{Chi/L}$ $2f_{non\text{-}NAY}$ and classified into 15 bins with similar sample size. $CUB_{Chi/L}$ and $CUB_{Chi/L\_binexp}$ were calculated using codon frequencies pooled among CDS for each bin. To calculate bin-specific SI GC content, we pooled intron sites for each of the $CUB_{Chi/L}$ $2f_{non\text{-}NAY}$ bins. Error bars indicate 95% CIs among 1000 bootstrap replicates.

**Fig. S11. Compositional trends among 2-fold synonymous families in *D. melanogaster*.**

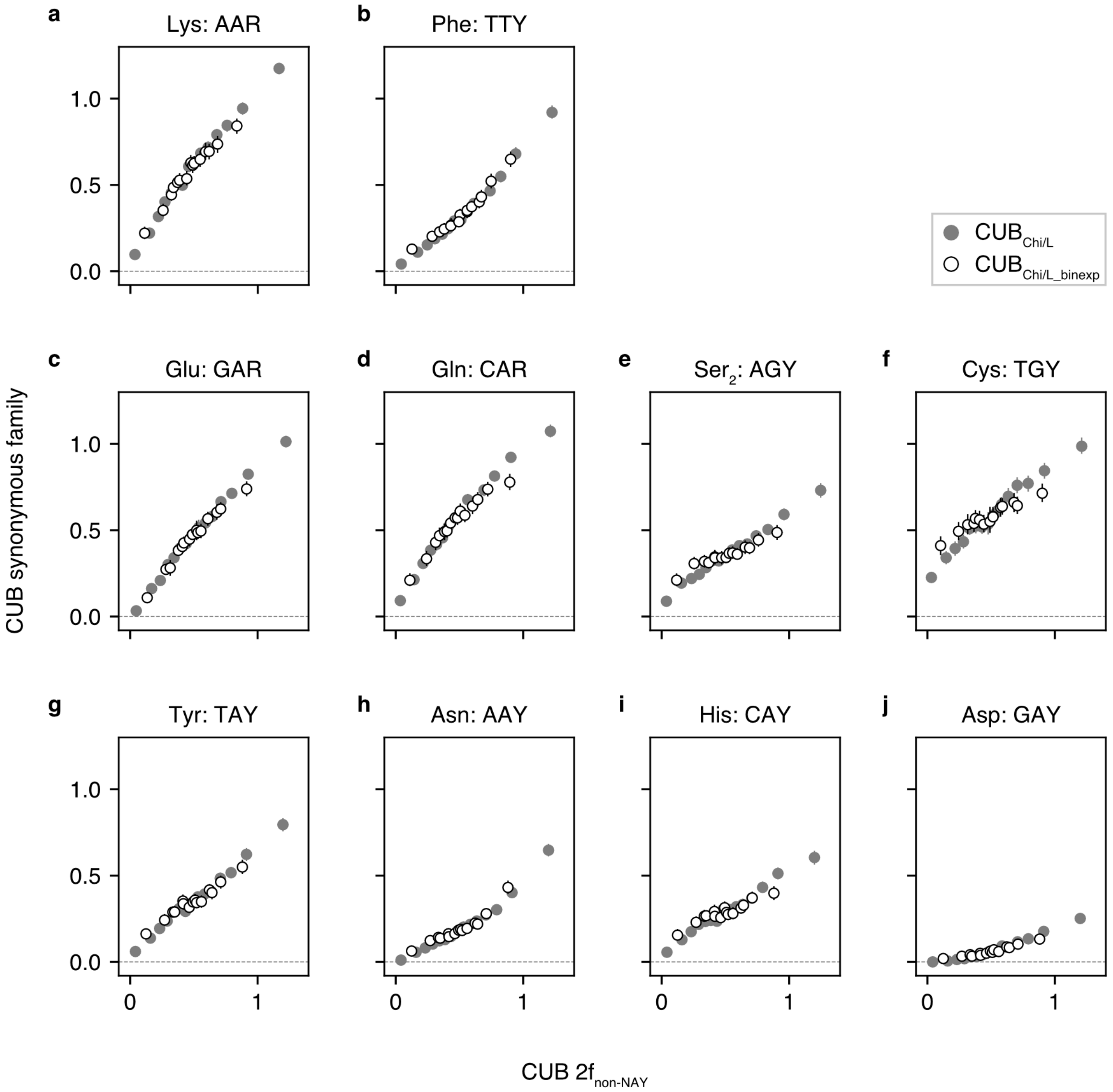

Compositional bias of individual 2-fold synonymous families is compared to a proxy for the magnitude of translational selection, CUB $2f_{non\text{-}NAY}$. Codons for a synonymous family being analyzed are excluded from predictor codons. Positive CUB values indicate that G-ending or C-ending codon usage is greater than SI GC content. Panels are ordered using W→S $\gamma$ values for *D. simulans* autosomal data (see Fig. 3a). Error bars indicate 95% CIs among 1000 bootstrap replicates.

**Fig. S12. Compositional trends among 2-fold synonymous families in *D. pseudoobscura*.**

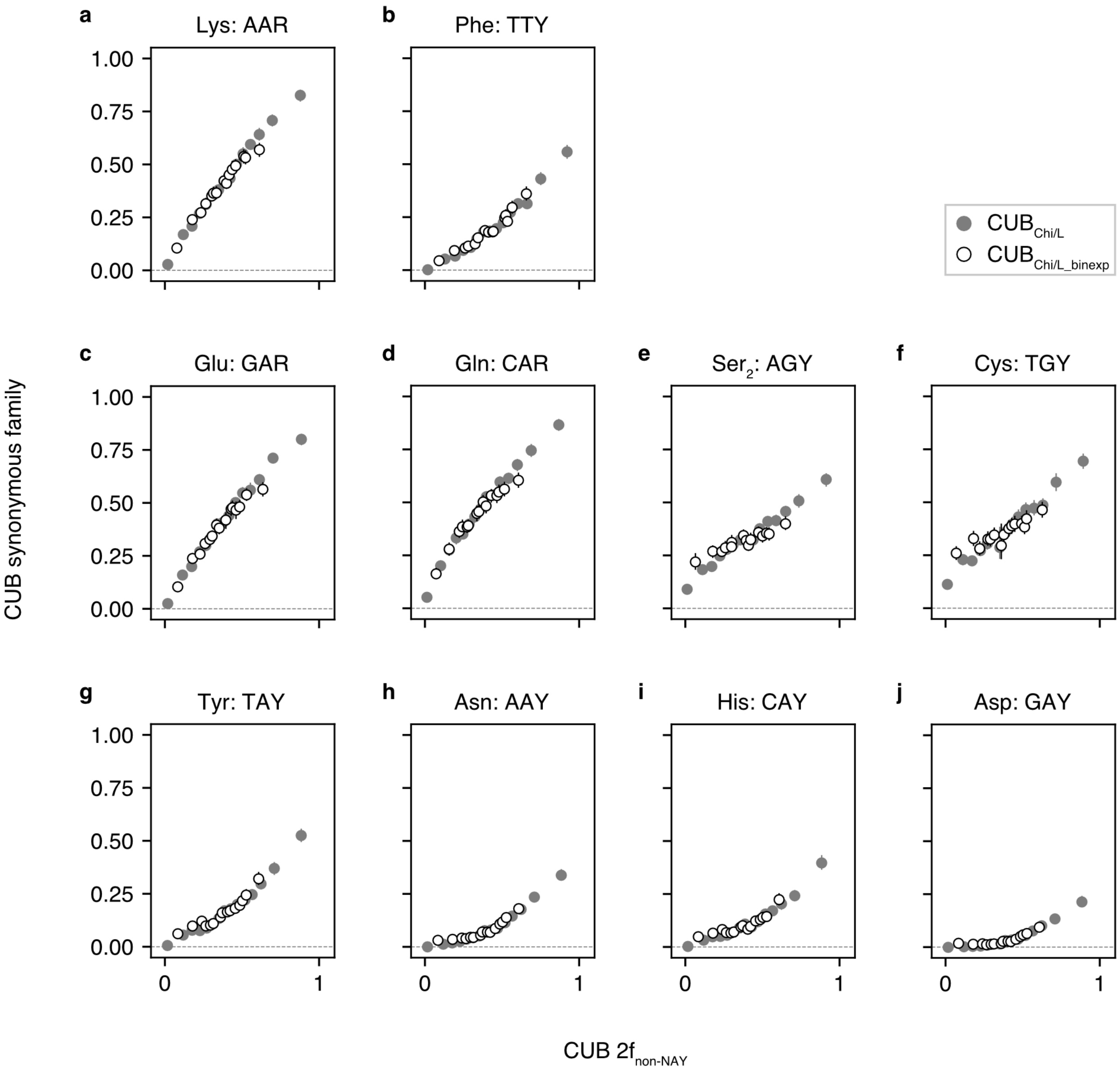

CUB of individual 2-fold synonymous families is compared to a proxy for the magnitude of translational selection, CUB $2f_{non\text{-}NAY}$. Codons for a synonymous family being analyzed are excluded from the predictor calculation. Positive CUB indicate that G-ending or C-ending codon usage is greater than SI GC content. Panels are ordered using W→S $\gamma$ values for *D. simulans* autosomal data (see Fig. 3a). Error bars indicate 95% CIs among 1000 bootstrap replicates.

**Fig. S13. Compositional trends among 2-fold synonymous families in *D. willistoni*.**

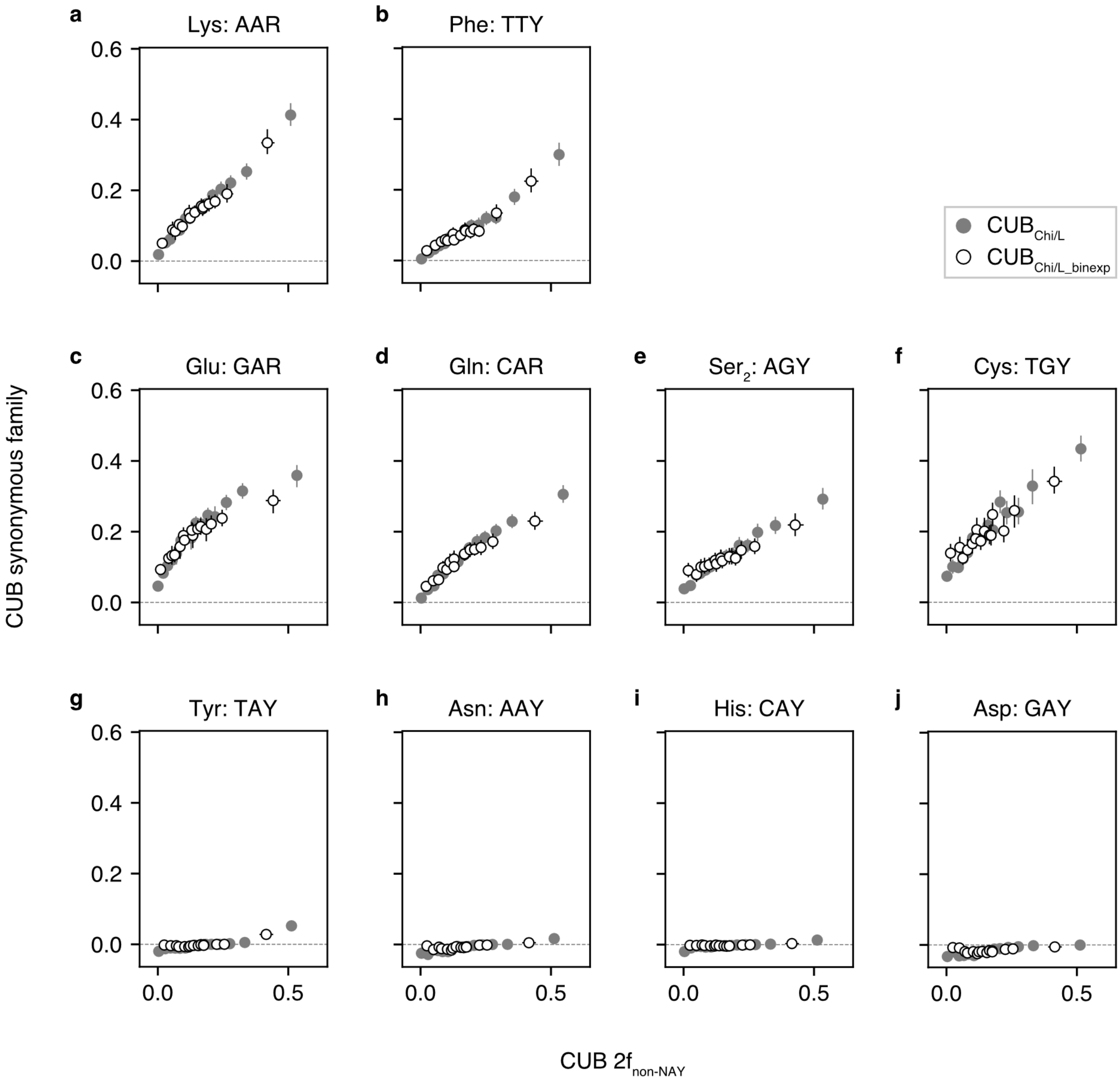

CUB of individual 2-fold synonymous families is compared to a proxy for the magnitude of translational selection, CUB $2f_{non\text{-}NAY}$. Codons for a synonymous family being analyzed are excluded from the predictor calculation. Positive CUB indicate that G-ending or C-ending codon usage is greater than SI GC content. Panels are ordered using W→S γ values for *D. simulans* autosomal data (see Fig. 3a). Error bars indicate 95% CIs among 1000 bootstrap replicates.

**Fig. S14. Compositional trends among 2-fold synonymous families in *D. grimshawi*.**

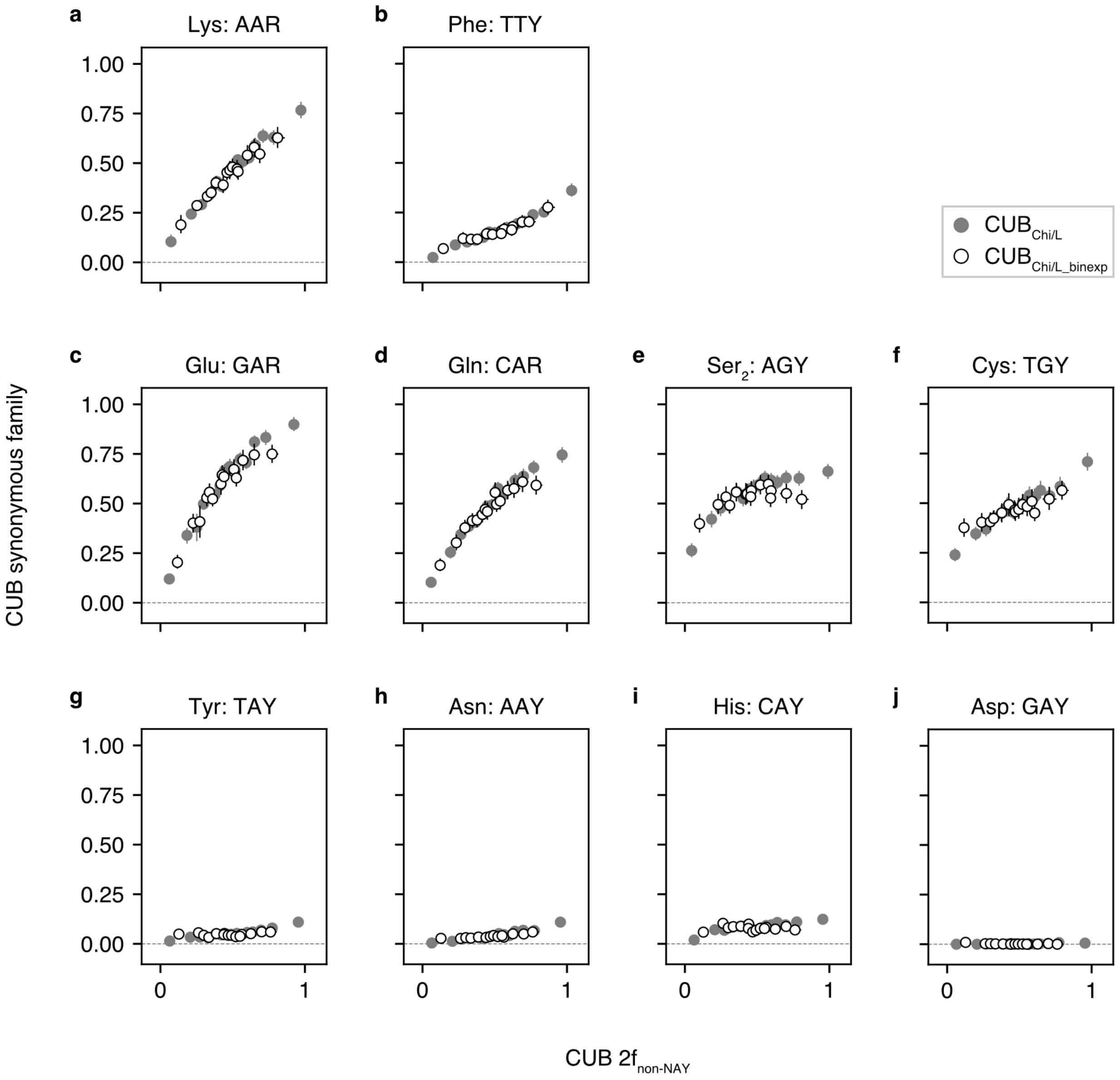

CUB of individual 2-fold synonymous families is compared to a proxy for the magnitude of translational selection, CUB 2f$_{non\text{-}NAY}$. Codons for a synonymous family being analyzed are excluded from the predictor calculation. Positive CUB indicate that G-ending or C-ending codon usage is greater than SI GC content. Panels are ordered using W→S $\gamma$ values for *D. simulans* autosomal data (see Fig. 3a). Error bars indicate 95% CIs among 1000 bootstrap replicates.

**Fig. S15. Compositional trends among 2-fold synonymous families in *D. mojavensis*.**

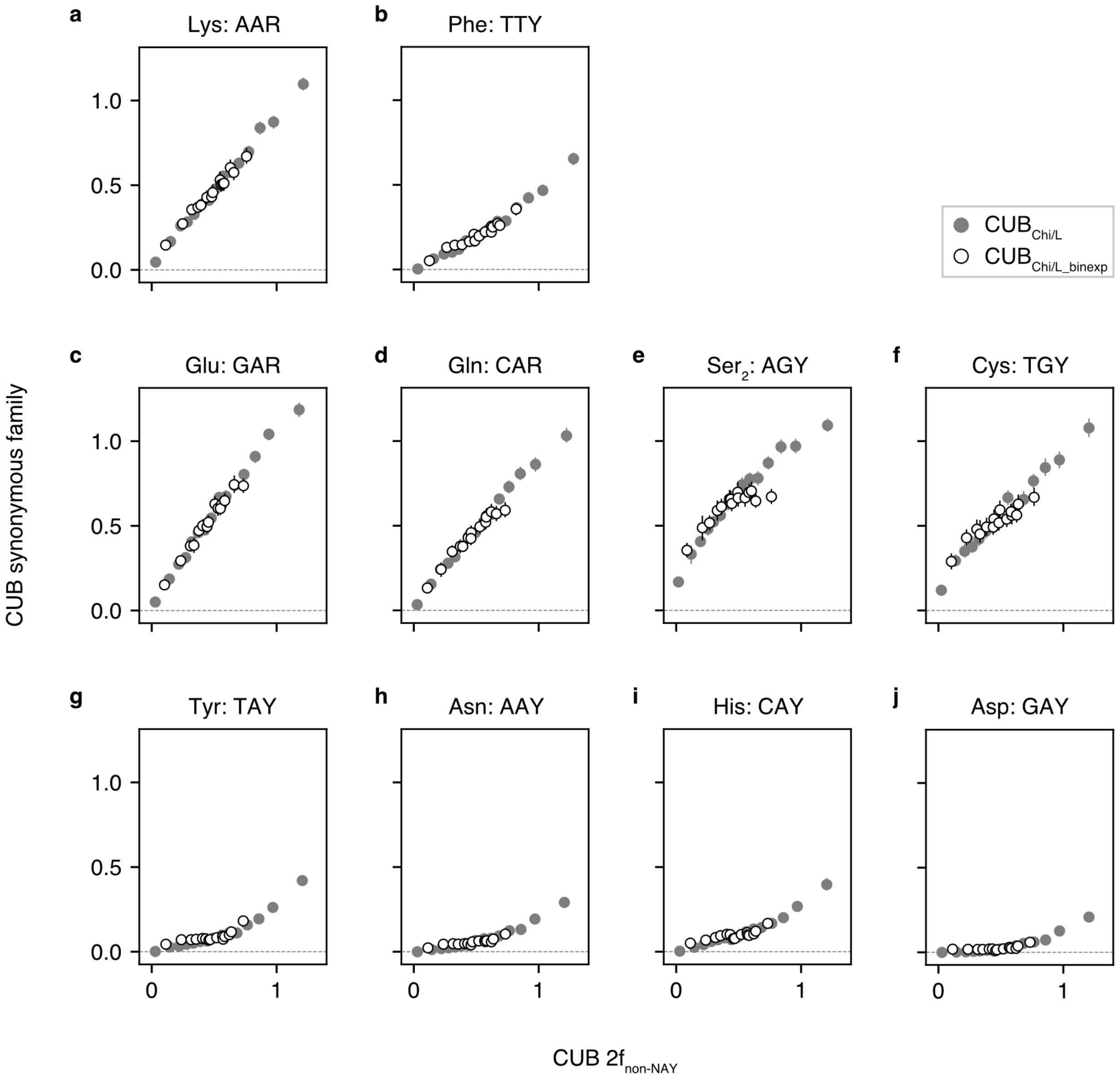

CUB of individual 2-fold synonymous families is compared to a proxy for the magnitude of translational selection, CUB $2f_{non\text{-}NAY}$. Codons for a synonymous family being analyzed are excluded from the predictor calculation. Positive CUB indicate that G-ending or C-ending codon usage is greater than SI GC content. Panels are ordered using W→S γ values for *D. simulans* autosomal data (see Fig. 3a). Error bars indicate 95% CIs among 1000 bootstrap replicates.

**Fig. S16. Compositional trends among 2-fold synonymous families in *D. virilis*.**

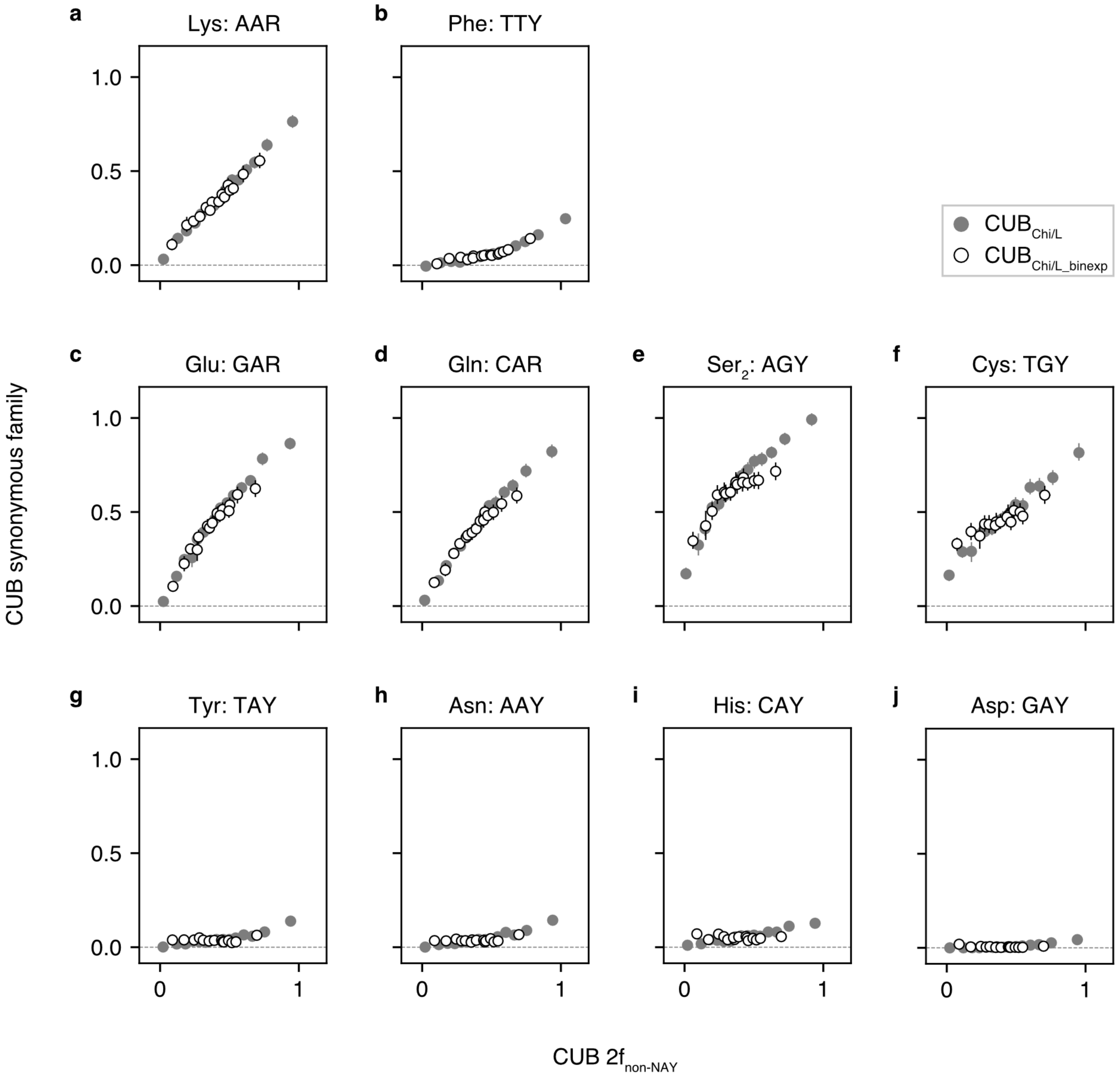

CUB of individual 2-fold synonymous families is compared to a proxy for the magnitude of translational selection, CUB 2f$_{non\text{-}NAY}$. Codons for a synonymous family being analyzed are excluded from the predictor calculation. Positive CUB indicate that G-ending or C-ending codon usage is greater than SI GC content. Panels are ordered using W→S γ values for *D. simulans* autosomal data (see Fig. 3a). Error bars indicate 95% CIs among 1000 bootstrap replicates.

**Fig. S17. Synonymous family-specific compositional trends among distantly related Drosophila species.**

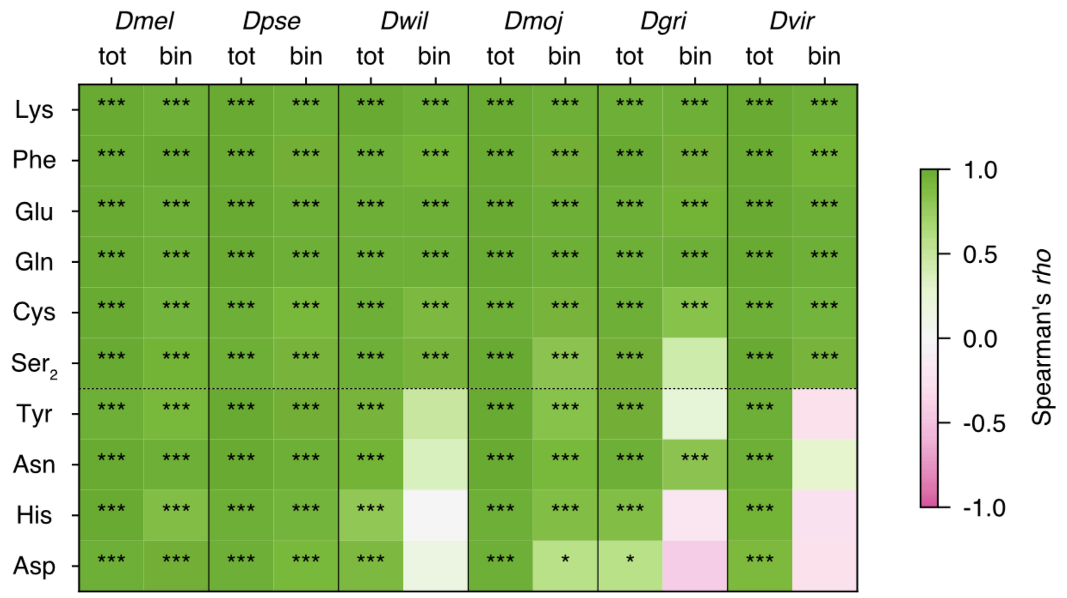

CUB of individual synonymous families were plotted against a proxy for the magnitude of translational selection, CUB $2f_{non\text{-}NAY}$ (excluding a synonymous family being tested). Previous analyses (Heger and Ponting 2007b; Vicario *et al.* 2007) support GC preference at $2f_{non\text{-}NAY}$ codons among these *Drosophila* species. The top six rows show $2f_{non\text{-}NAY}$ families and the bottom four rows show NAY families. “tot” and “bin” indicate analysis of $CUB_{Chi/L}$ and $CUB_{Chi/L\_binexp}$, respectively. Autosomal loci are employed. CDS with < 10 $2f_{non\text{-}NAY}$ codons are excluded. CDS are ranked by $CUB_{Chi/L}$ $2f_{non\text{-}NAY}$ and classified into 15 bins with roughly similar numbers of $2f_{non\text{-}NAY}$ codons. $CUB_{Chi/L}$ and $CUB_{Chi/L\_binexp}$ were calculated using codon frequencies pooled among CDS within a bin. Rank correlation of CUB between $2f_{non\text{-}NAY}$ and individual synonymous family was examined across 15 bins. Green and magenta indicate positive and negative relationships between CUB $2f_{non\text{-}NAY}$ and CUB, respectively, for a synonymous family of interest. Color intensity indicates Spearman’s rank correlation coefficient. The sequential Bonferroni method was employed for multiple test corrections among synonymous families for each species (*i.e.*, 10 tests are corrected). *, **, and *** indicate $p < 0.05$, $< 0.01$, and $< 0.001$, respectively. See Figs. S11 to S16 for data used to examine correlations.

**Fig. S18. Elevated GC-ending codon usage in predicted highly expressed genes: $CUB_{Chi/L}$ $2f_{non\text{-}NAY}$ and transcript abundance**

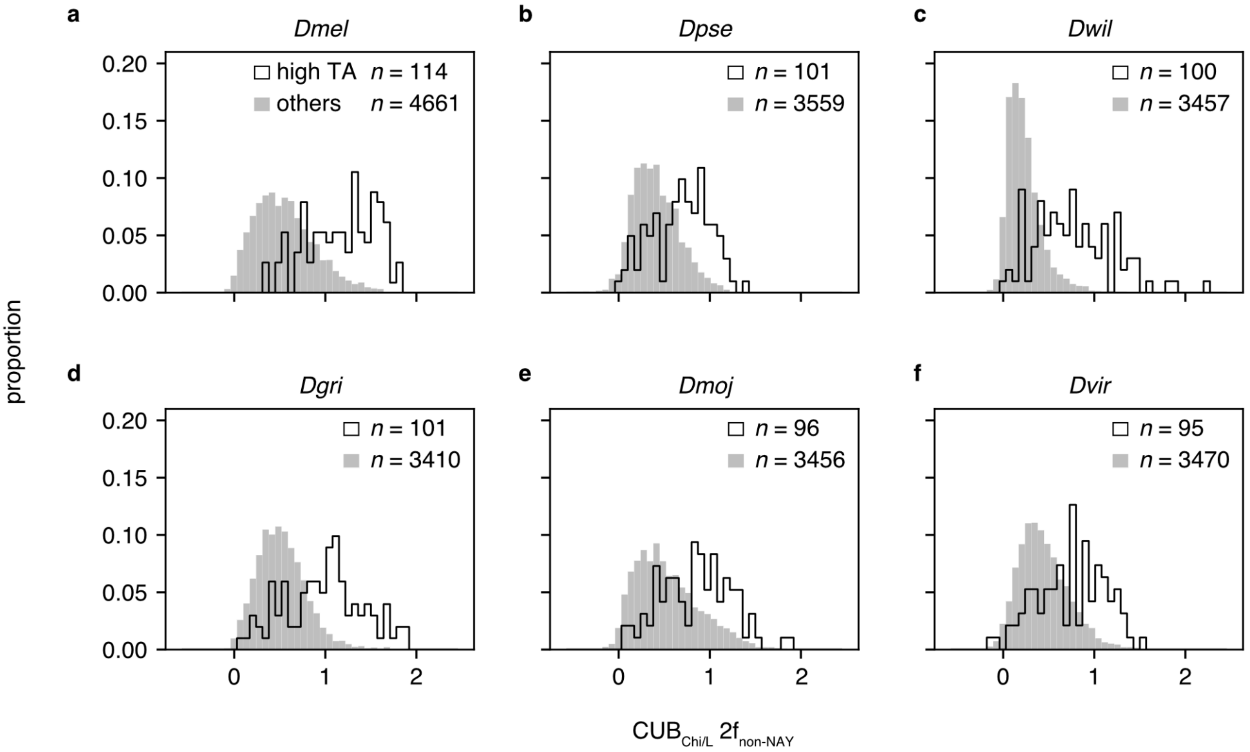

$CUB_{Chi/L}$ $2f_{non\text{-}NAY}$ are compared between predicted highly expressed genes or their 1-to-1 OrthoFinder -orthologs (black unfilled) and other genes (gray filled). (a) *D. melanogaster*, (b) *D. pseudoobscura*, (c) *D. willistoni*, (d) *D. grimshawi*, (e) *D. mojavensis*, and (f) *D. virilis*. Highly expressed genes are defined as genes for which transcript abundance (TA) is among the top 100 and TA dispersion (mean / variance) is < 2.5-percentile among eligible genes in *D. melanogaster*. TA estimates are from microarray data from 22 tissues: adult (ad) hindgut, ad midgut, ad male accessory gland, ad brain, ad crop, ad ovary, ad testis, ad salivary gland, ad carcass, ad fat body, ad eye, ad heart, ad male ejaculatory duct, larva (lv) feeding (fd) hindgut, lv fd midgut, lv fd salivary gland, lv fd malpighian tubule, lv fd fat body, lv fd carcass, lv fd central nervous system, lv fd trachea, and embryo (4h to 10h) (Chintapalli *et al.* 2007; Matsumoto *et al.* 2016). Positive $CUB_{Chi/L}$ values indicate G- or C-ending codon usage higher than GC content of short introns. CDS with > 600 codons are filtered. CDS with < 20 $2f_{non\text{-}NAY}$ codons are filtered.

**Fig. S19. Elevated NAC codon usage in predicted highly expressed genes: $CUB_{Chi/L}$ NAY and transcript abundance.**

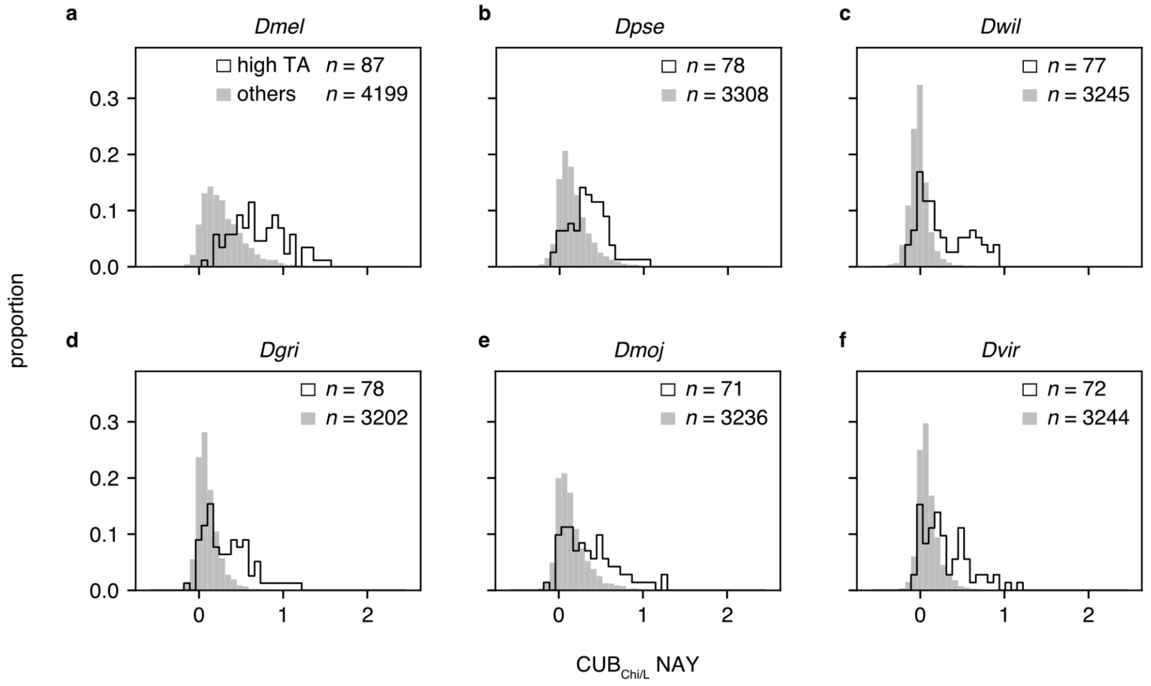

$CUB_{Chi/L}$ NAY are compared between predicted highly expressed genes or their 1-to-1 OrthoFinder-orthologs (black unfilled) and other genes (gray filled). The definition of highly expressed genes is the same as Fig. S17. Positive $CUB_{Chi/L}$ values indicate NAC usage higher than GC content of short introns. (a) *D. melanogaster*, (b) *D. pseudoobscura*, (c) *D. willistoni*, (d) *D. grimshawi*, (e) *D. mojavensis*, and (f) *D. virilis*. CDS with > 600 codons are filtered. CDS with < 20 NAY codons are filtered.

**Fig. S20. Testing major codon preference: fixation bias vs major codon usage.**

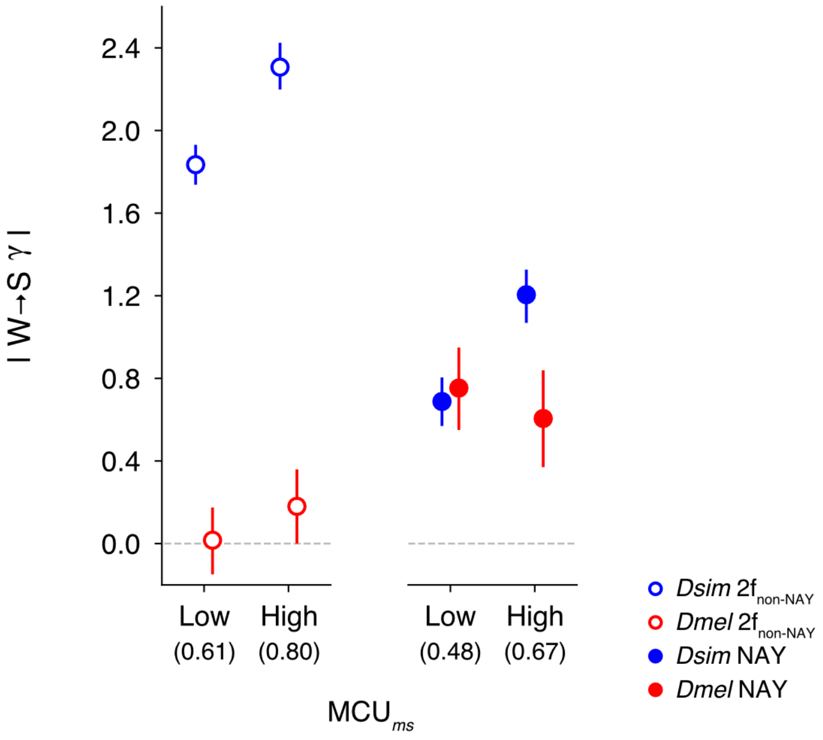

Ancestral major codon usage (MCU) was assessed as a predictor of fixation bias intensity ($\gamma$) at autosomal loci. Ancestral reconstructions were pooled into two classes of NAY (Asp, His, Asn, and Tyr; filled circles) and $2f_{non\text{-}NAY}$ synonymous families (Phe, Cys, $Ser_2$, Lys, Gln, and Glu; open circles) for each gene. MCU at the *ms* node ($MCU_{ms}$) is used for categorizing genes into low and high MCU classes. $MCU_{ms}$ for NAY and $2f_{non\text{-}NAY}$ codons are used for binning in the analysis of NAY and non-NAY, respectively. Values in parenthesis indicate the average $MCU_{ms}$. Genes with 15 or fewer codons are filtered. For each MCU class, fixation biases acting between GC-increasing and AT-increasing synonymous changes are estimated using polymorphisms in the *D. simulans* ("*Dsim*"; blue) and *D. melanogaster* ("*Dmel*"; red) populations. For the required neutral reference for $\gamma$ estimation, we employed GC-conservative changes within short introns within a bin. Note that absolute values of fixation biases (regardless of GC or AT preference) are plotted. Dotted horizontal lines indicate $\gamma$ = 0. Error bars indicate 95% CI among 1000 bootstrap replicates. Ancestral reconstructions are resampled in units of genes within a bin.

### Supplementary Methods

#### *Identifying orthologs among the D. melanogaster subgroup species*

##### Genome sequences

We identified orthologs for protein-coding genes among four species, *Dmel*, *Dsim*, *Drosophila yakuba* (*Dyak*), and *Drosophila erecta* (*Dere*) from the *D. melanogaster* subgroup. We obtained genomic sequences and annotations from FlyBase (ftp://ftp.flybase.net/genomes/) for *Dmel* (r6.24 FB2018_05; last downloaded on 12th July 2019), *Dsim* (r2.02 FB2017_04; last downloaded on 9th March 2020), *Dyak* (r1.05 FB2016_05; last downloaded on 12th July 2019) and *Dere* (r1.05 FB2016_05; last downloaded on 12th July 2019). For genes with multiple protein isoforms, we employed the longest CDS. We filtered genes for which CDS lengths were not multiples of three. We employed *Drosophila ananassae* (*Dana*; r1.06 FB2018_04; last downloaded on 16th October 2019) and *Drosophila pseudoobscura* (*Dpse*; r3.04 FB2018_05; last downloaded on 16th October 2019) as outgroups.

##### Identifying putative ortholog groups

We combined two approaches to identify putative orthologs among the annotated genes. One approach employed the FlyBase (Thurmond *et al.* 2018) ortholog annotations for *Dmel* genes across 12 Drosophila species (ftp://ftp.flybase.net/releases/FB2018_05/precomputed_files/orthologs/dmel_orthologs_in_drosophila_species_fb_2018_05.tsv.gz; last downloaded on 26th December 2019). We obtained 13,493 putative ortholog groups that include at least two representatives among *Dmel*, *Dsim*, *Dyak*, *Dere*, *Dana*, and *Dpse*. The second approach employed protein sequence similarity searches using

OrthoFinder (Emms and Kelly 2015; version 2.3.3; downloaded on 13th September 2019) and yielded 12,789 putative ortholog groups.

We fused the putative ortholog groups from FlyBase and OrthoFinder as a first step to identify sets of *Dmel*, *Dsim*, *Dyak* and *Dere* protein-coding genes that are consistent with the assumed species tree topology and that appear to be evolving independently of other sets (*msye* ortholog sets). FlyBase and OrthoFinder groups were fused if groups shared one or more members. We obtained 16,193 groups among which 10,320 included single representatives in each species (*msye* orthogroups) and 1,384 had at least one representative each from *Dmel*, *Dsim*, *Dyak* and *Dere* but more than one representative for at least one species (multiple candidate *msye* orthogroups). Other groups were missing representatives from one or more species of the *D. melanogaster* subgroup and were not included in further analyses.

**Orthogroups with single representatives from each *D. melanogaster* subgroup species**

We employed phylogenetic and DNA distance approaches to exclude potentially misassigned orthologs and/or questionable alignments from *msye* orthogroups. *Dana* and *Dpse* genes were included from FlyBase and OrthoFinder groups when available. We aligned predicted protein sequences within orthogroups using the E-INS-I method within the MAFFT software package (Katoh *et al.* 2002) and replaced amino acids with codons in the corresponding positions. We removed codons at which any of the aligned codons included gaps and/or non-ATGC characters. Nine groups were eliminated because no codons remained after this process. We estimated gene trees using maximum parsimony (bootstrap resampling of nucleotide sites, $n$ = 1,000) and determined supported clades among gene trees for the bootstrap replicates for each orthogroup using the “majority rule extended method” (implemented in the *consense* program of Phylip). Maximum parsimony was employed to allow bootstrap analyses for each orthogroup. Bootstrap resampling, parsimony tree estimation and consensus tree estimation were conducted using *seqboot*, *dnapars* and *consense* programs respectively in Phylip (Felsenstein 2004; version 3.697; last downloaded on 25th September 2019). We filtered orthogroups in which *Dana* and *Dpse* genes were placed (bootstrap support ≥ 50%) within a *D. melanogaster* subgroup clade because *Dana* and *Dpse* are established as distantly related to this subgroup (Kopp and True 2002; Ko *et al.* 2003; Akashi *et al.* 2006; Pollard *et al.* 2006). Among the 10,311 phylogenetic trees, 26 groups were rejected by this criteria. Because *Dmel*, *Dsim*, *Dyak* and *Dere* are closely related species (Ko *et al.* 2003; Akashi *et al.* 2006; Heger and Ponting 2007a; Wong *et al.* 2007), we do not expect relationships among these species to be resolvable for most single genes and did not evaluate topologies within the *D. melanogaster* subgroup clade.

To reduce the proportion of potentially misaligned/misannotated data, we applied additional filtering based on sequence distance. We tested levels of synonymous divergence ($d_S$) for all CDS pairs within orthogroups. For each alignment, we generated sliding windows of 50 codons with a step size of one codon. For each window, pairwise $d_S$ was estimated based on the Nei-Gojobori method (Nei and Gojobori 1986) implemented in CODEML (Yang 2007; version 4.9). The values were tested against thresholds of 0.7 for *Dmel/Dsim* pairs and 1.0 for other species pairs. We excluded a group if more than 75% of codons were found in high $d_S$ (> threshold for at least one pair) windows or if less than 40 codons remained after removing codons in high $d_S$ windows. We filtered 37 *msye* orthogroups according to these criteria. For this filtering and other steps described below, arbitrary thresholds were chosen to exclude groups with extreme values (usually a few percent).

**Extracting ortholog sets from multiple candidate *msye* orthogroups**

We examined phylogenetic relationships, DNA distances, synteny, and alignment lengths to extract *msye* ortholog sets from the 1,384 multiple candidate *msye* orthogroups. We considered candidate *msye* ortholog set-containing clades that can be explained by simple scenarios of gene duplications on lineages prior to, and within, the *D. melanogaster* subgroup (Fig. S21). The clade support requirements shown in Fig. S21 were designed to filter cases of gene conversion among paralogs following gene duplications. We found 1,788 clades that may contain one or more *msye* ortholog sets.

We filtered *msye* sets within the extracted clades to obtain a single set from each clade. We considered all possible *msye* sets within the clades as candidates. We filtered these sets based on synonymous divergence as described above. In addition, we tested synteny conservation (*i.e.*, sharing of neighboring ortholog genes). For each *msye* set, we examined up to 20 neighboring genes (10 genes 5′ and 10 genes 3′) from the *msye* ortholog sets that we obtained among *msye* orthogroups with single representatives from each species. If fewer than 10 genes were available, we employed as many genes as possible. In each species pair, we counted the numbers of the neighboring genes that belong to the same *msye* set and summed the counts across all pairs of species. We retained candidate *msye* sets that had the highest overall counts for a given clade. For the remaining clades with more than one *msye* set possibility, we selected one set having the greatest number of aligned codons across the four species (and randomly chose among sets showing the same number of the aligned codons). Overall, we obtained 1,774 *msye* ortholog sets after the filtering process to give a total of 12,022 *msye* ortholog sets for the analysis.

**Fig. S21. Phylogenetic approach for ortholog assignment.**

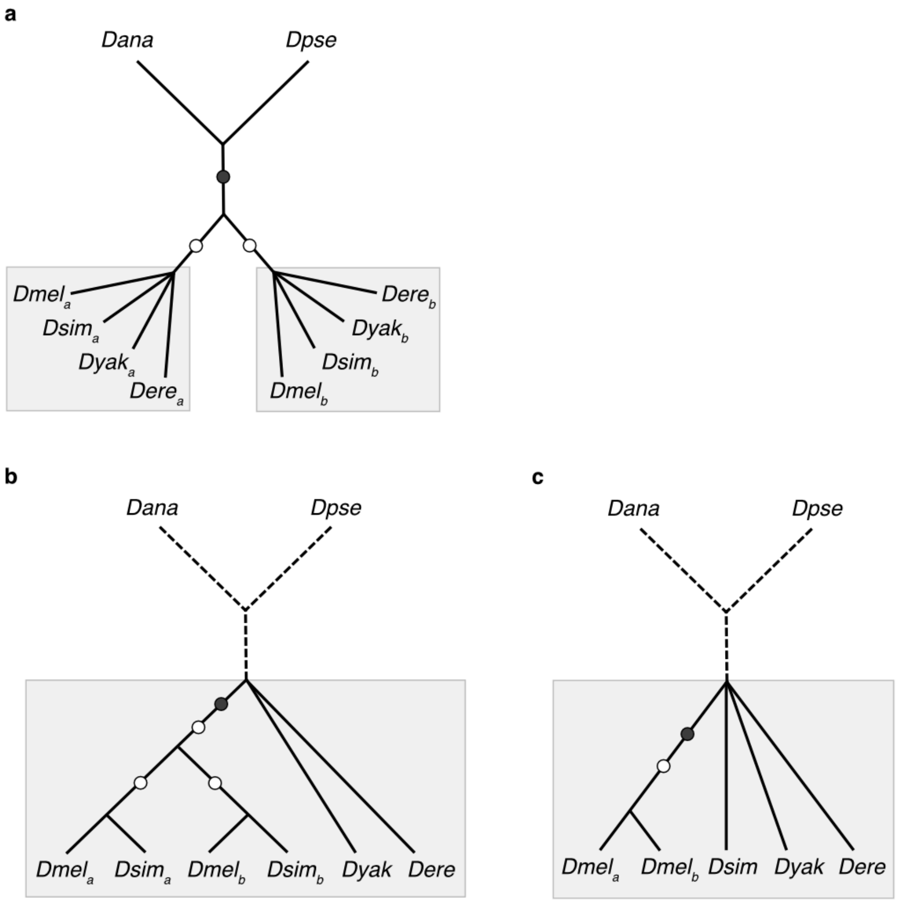

Scenarios of clades that contain candidate *msye* ortholog sets (gray boxes) are shown. Three gene duplication scenarios were considered: (a) gene duplication prior to the ancestor of the *D. melanogaster* subgroup (*msye* node), (b) gene duplication on an internal branch within the *D. melanogaster* subgroup (prior to the *Dmel*, *Dsim* or the *Dyak*, *Dere* split) and (c) gene duplication on a terminal branch within the *D. melanogaster* subgroup. Locations of inferred gene duplications are shown as filled circles. Clades requiring bootstrap support (≥50%, 1000 replicates) are indicated with open circles. Relationships among the *D. melanogaster* subgroup members are shown as star trees when the topology is not considered for inclusion. *Dana*/*Dpse* outgroup lineages are shown as dotted lines when outgroup support is not required for identification of duplication lineages. We included cases with mixtures of (a), (b) and (c) duplication types if the extant gene configuration can be explained by a single most parsimonious scenario that combines such gene duplications.

**Obtaining intron ortholog sets**

We examined gene structure consistency within *msye* ortholog sets to identify putative intron orthologs. We used the *Dmel* genome annotation as a reference and considered ortholog candidates in *Dsim*, *Dyak*, and *Dere* that showed consistent splice positions (*i.e.*, predicted splice sites occur within the same codon positions within CDS alignments). Among introns that did not qualify under this criteria, those located between intron pairs showing consistent splice positions within genes (*i.e.*, consistent *order* within orthologous genes) were also included as candidates. We employed pairs in which *Dmel* introns have lengths less than or equal to 100 bp for the further analyses of 22,209 candidates for orthogroups that include a single representative each from *Dmel*, *Dsim*, *Dyak* and *Dere* (*msye* intron orthogroups).

We filtered some of the candidates to insure non-overlapping sets of *msye* orthogroups and to remove misannotated introns or questionable alignments. We examined pairwise sequence alignments between *Dmel* and *Dsim*, *Dyak*, and *Dere*. We retained orthogroups in which all pairs passed cutoffs for numbers of aligned nucleotides (40 bp) and sequence similarity (for pairs showing consistent splice positions, 0.50 for *Dmel*/*Dsim* and 0.40 for *Dmel*/*Dyak* and *Dmel*/*Dere*, for pairs showing consistent order within genes, 0.712, 0.552 or 0.554, respectively). We filtered 85 orthogroups (0.38%) by these filters. In addition, we examined lengths/numbers of gaps for the latter pairs. We filtered pairs that included extensive gap regions within alignments (proportion of aligned nucleotides lower values than 0.833, 0.738 or 0.730 for the three species pairs, respectively). We also filtered cases with large *numbers* of gaps (number of gaps scaled to alignment length greater than 0.0317, 0.0405 or 0.0408 for the larger value in each pair). These filters excluded 126 orthogroups (0.59%). We excluded an additional four *msye* intron orthogroups in which members overlapped. Overall, we obtained 21,994 *msye* intron ortholog sets. For the analyses, we employed short introns, those with length 100 bp or less in both *Dmel* and *Dsim*.

##### *Within-species DNA sequences*

For lines from *Dmel* populations, we extracted CDS and intron sequences after converting FlyBase r5.28 genome annotations to r6.24 versions using an annotation mapping table (https://github.com/FlyBase/bulkfile-scripts/blob/master/dmel_r5_to_r6/dmel_r5_to_r6_mapping.tsv ; last downloaded on 30th July 2019). Genes with different numbers of exons between r5.28 and r6.24 were excluded from the data set.

We mapped FlyBase reference (r2.02) to a reference sequence (Hu *et al.* 2013) that was used for the reconstruction of genome sequences for within-species samples of *Dsim* (Rogers *et al.* 2014; Jackson *et al.* 2017). We compared the main scaffolds (2L and Scf_2L, 2R and Scf_2R, 3L and Scf_3L, 3R and Scf_3R, 4 and Scf_4, and X and Scf_X) using MUMmer (Kurtz *et al.* 2004; version 3.23; last downloaded on 6th March 2020) and obtained coordinates of 1-to-1 matching regions using the *delta-filter* command with *-1* option. FlyBase sequences showed high sequence identity with the Hu *et al.* (2013) sequences (minimum 99.99%). Using the annotation mapping table, we converted FlyBase gene coordinates to the Hu *et al.* (2013) genome.

##### *Sequence alignments*

We aligned amino acid translations for the ortholog CDS's using the E-INS-i algorithm implemented in the MAFFT program (Katoh *et al.* 2002) and back-translated to nucleotides. We used the reference sequence alignments of orthologs as a "backbone" to align within-species data. We inserted sequences for the within-species lines (14 for the *Dmel* population and 21 for the *Dsim* population) to the alignments by mapping codon or nucleotide positions to the corresponding positions of the reference sequences. The *Dmel* and *Dsim* reference sequences were removed from the sequence alignments prior to analyses.

We filtered data from heterochromatic/low crossover regions defined in (Kliman and Hey 1993). We assigned alignment data in such regions using chromosome map positions of *Dmel* genes (ftp://ftp.flybase.net/releases/FB2018_05/precomputed_files/genes/gene_map_table_fb_2018_05.tsv.gz; last downloaded on 29th December 2020). We also filtered some regions within the remaining CDS and intron sequence alignments. We filtered codons/sites that include any gaps among the aligned sequences. We used the *Dmel* genome annotation to filter codons/sites that overlap with predicted transposable elements and/or of transcripts from other genes. In addition, we restricted the analysis to sites that are included in all predicted CDS isoforms for a given gene. Finally, we filtered some putatively functionally constrained regions within introns (Halligan *et al.* 2004; Halligan and Keightley 2006; Parsch *et al.* 2010): 10 bases at the 5′ splice junctions and 30 bases at the 3′ splice junctions in the sequence alignments. Filtering statistics are shown in Table S19. We used these alignments as the starting point for inferring polymorphic and fixed changes.

**Table. S19. Data filtering statistics.**

| Site class [a] | Chr [b] | # alignments [c] | KH [d] | TE overlap [e] | Alternatively spliced | Transcript overlap [f] | # alignments [g] |
|---|---|---|---|---|---|---|---|
| CDS | A | 10,122 | 761 | 491 | 334,393 | 446,299 | 8,166 |
| | X | 1,746 | 133 | 304 | 67,528 | 100,456 | 1,382 |
| Intron | A | 18,719 | 1,475 | - | 8,765 | 77,852 | 15,927 |
| | X | 2,705 | 141 | - | 5,370 | 13,584 | 2,350 |

[a] "Intron" indicates short introns.
[b] Autosomal (A) and X-linked loci.
[c] Numbers prior to filtering.
[d] Numbers of CDS or introns located in heterochromatic/lowly recombining regions defined by Kliman and Hey (1993). Data in these regions are filtered because such regions may experience different mutational spectra (Takano-Shimizu 2001; Marais *et al.* 2001, 2003; Singh *et al.* 2005) as well as reduced efficacy of natural selection (Fisher 1930; Muller 1932; Crow and Kimura 1965; Felsenstein 1974) compared to euchromatic regions.
[e] Numbers of codons overlapping with transposable elements.
[f] Numbers of codons/ intronic sites overlapping with transcripts for other genes.
[g] Numbers after filtering

#### *Statistical methods*

##### Ancestral inference

We employed a likelihood-based method that incorporates non-stationary and biased base composition evolution in the gene tree. Previously, we developed an approximate method for data that includes within-species variation in recombining regions where gene trees may not be shared across sites. We first convert sequence alignments to a format suitable for ancestral inference and then estimate the probabilities of ancestral nucleotides by fitting a nucleotide substitution model to input data. Finally, we weight the probabilities using site frequencies. Computer simulations showed high reliability of this approach under a range of scenarios that attempt to emulate base composition evolution in the *Drosophila melanogaster* subgroup species (Matsumoto *et al.* 2015; Matsumoto and Akashi 2018).

We implemented the BASEML-BTW method in a software package (https://github.com/nigevogen). The package takes sequence alignments and an assumed topology as input for the following three processes: 1) convert sequence alignments to bifurcating tree input (Figure 2 in Matsumoto and Akashi 2018), 2) perform ancestral inference using BASEML (Yang 2007), and 3) weight the ancestral inference using observed and expected site frequencies. BASEML estimates evolutionary parameters (branch lengths and transition parameters in a nucleotide substitution model) from terminal node nucleotide configurations (TNNC). Based on these parameters, sets of internal node nucleotide configurations (INNC) and their probabilities (joint reconstructions) are generated for each observed TNNC. The weighted probabilities of INNC for each site in the input data (weighted joint reconstructions) are the final product of the BTW method.

In our analyses, we generated the BASEML input sequences from alignments as described above and assumed the tree topology of Fig. 1b. We obtained the weighted joint reconstructions at ancestral nodes, *ms*, *ye*, *m*' and *s*', for each site. For ancestral inference by BASEML, we employed the GTR nucleotide substitution model (Tavaré 1986). Our approach attempts to account for both biased base composition evolution and lineage-specific transition parameters [*a~f* and $\pi_T$, $\pi_C$, $\pi_A$, $\pi_G$ in GTR-$NH_b$ substitution model, Matsumoto *et al.* (2015)]. In addition, we employed a newly implemented option to assign the same transition parameters to user-defined branches ($m'$-$m_{c1}/m_{c2}$ and $s'$-$s_{c1}/s_{c2}$ in Fig. 1b). For the weighting process, we employed an iterative approach [$BTW_{est}$ in (Matsumoto and Akashi 2018)]. The first round weights the BASEML probabilities of INNC at each site using observed allele frequencies and the expected SFS under neutral equilibrium (Fisher 1930; Wright 1969). The weighted probabilities are employed to estimate the SFS (see below) which is then used for weighting in the next round. We conducted five iterations to allow the probabilities to converge.

We employed concatenated alignments for a given mutation class as input for the ancestral inference process (described below). For inference of synonymous mutations, we extracted 3rd codon positions from aligned codons that belong to the same synonymous family and concatenated the data across genes. Pooled family analysis included codons from all 2-fold or all 4-fold synonymous families. For inference of intronic mutations, we concatenated sites across introns.

**Inferring counts of mutations in the gene tree**

We inferred numbers of changes found polymorphic or fixed within species samples following Matsumoto and Akashi (2018). We assumed a minimal change model (no multiple or reverse changes) and probabilities were used as counts of nucleotide changes. Nucleotide changes inferred between nodes *m'* and *s'* and *Dmel* and *Dsim* terminal nodes ($m_{c1}$, $m_{c2}$, $s_{c1}$ and $s_{c2}$ in Fig. 1b) were classified as "polymorphic" mutations. The assumption of minimal evolution within a branch can result in slight underestimation of fixation counts given the estimated branch lengths (Matsumoto and Akashi 2018).

**Ancestral inference for context-dependence analysis**

We inferred nucleotide changes with specified 5′ and 3′ neighboring sites to test for dinucleotide preferences. We refer to a given site as a "subject" site and the neighboring site as the "context" site (context sites are either the 5′ or 3′ nearest neighbor of the subject site depending on the analysis). A given subject site was considered for dinucleotide analysis if none of the terminal nodes (*Dmel*, *Dsim*, *Dyak* and *Dere*) have gaps at the required context site. Furthermore, we required a high probability that the nucleotide state at the context site has not changed after the split of *Dmel* and *Dsim*. We retained cases where context sites satisfy the following two criteria: (1) *Dmel* and *Dsim* terminal nodes show identical nucleotides and (2) the sum of probabilities is high (>=0.99) for INNCs/INCCs in which *ms*, *m'* and *s'* nodes show the identical nucleotides to the *Dmel* and *Dsim* terminal nodes. Accepted dinucleotides were treated similarly to nucleotides (above) to infer locations/counts of changes within the population samples and on ancestral branches.

**Permutation approach**

We employed a permutation approach to test the null hypothesis that W→S aDAF skews for two classes are drawn from the same distribution. In each replicate, we randomly permuted classes (reassigned labels) and re-calculated the W→S aDAF skew difference. We estimated *p* values as the proportions of replicates that show lower or higher W→S aDAF skew differences than the value from the actual data. For cases of zero replicates that satisfy this criteria, we set $p = 0.001$ (10,000 replicates) or $p = 0.0001$ (100,000 replicates). The number of replicates is indicated for each analysis. The classes depend on the analysis: autosomal vs X-linked, ApT↔ApC vs BpT↔BpC, or ApT↔GpT vs ApV↔GpV.

**Bootstrap estimates of confidence intervals**

Our bootstrap replicates resampled units of CDS or introns. In population genetics analysis, we performed independent ancestral inference as described above for each replicate unless noted otherwise. 95% confidence intervals (CI) were estimated as the range from 2.5th- to 97.5th-percentile of observed statistics among replicates (Efron 1979). The number replicates are shown in figure legends (300 or 1000).

#### *Measure of codon usage bias*

We employed a variant of the "scaled $\chi^2$ statistic (Shields *et al.* 1988; Akashi 1995) as a measure of the deviation of synonymous codon usage from a putatively neutral expectation. Goodness-of-fit tests are designed to compare G+C counts and A+T counts at synonymous positions to expectations based on observed GC content at SI. The $\chi^2$ statistics from such tests are signed to indicate whether the proportion of G- or C-ending codons among synonymous codons is greater (positive) or smaller (negative) than expected GC. $\chi^2$ values are summed across synonymous families and the sum is divided by the number of codons to give a measure that is not dependent on sample size (*i.e.*, numbers of codons), codon usage bias ($CUB_{Chi/L}$).

#### *Compositional trend analysis*

We employed six distantly related Drosophila species: *Dmel*, *Dpse*, *Dwil*, *Dgri*, *Dmoj*, and *Dvir*. For *Dmel* and *Dpse*, we used CDS and intron sequences to construct the *Dmel* subgroup data set. For other species, we extracted predicted CDS and intron sequences from the genome-scale DNA sequences using annotations. We downloaded DNA sequences and annotations from NCBI (https://www.ncbi.nlm.nih.gov/assembly/) for *Dwil* (101, GCF_000005925.1), for *Dgri* (103, GCF_018153295.1), for *Dmoj* (102, GCF_018153725.1), and for *Dvir* (103, GCF_003285735.1). These data from NCBI were downloaded on 13th October 2021. For genes with multiple CDS isoforms, we used the longest CDS. See below for details of filtering and a method to infer 1-to-1 OrthoFinder orthologs among these species.

We obtained SI sequences using transcript annotations. We excluded SI in untranslated regions and intron sites that are included in coding regions in one or more of transcript isoforms. Within introns, we filtered 10 bases at 5′ and 30 bases at 3′ splice junctions to reduce the contribution of functional constraint (Halligan *et al.* 2004; Halligan and Keightley 2006; Parsch *et al.* 2010). Only introns with 10 or more nucleotides remaining after these filtering steps were included in the analyses.

Major codons increase in usage within their synonymous family in genes experiencing higher MCP-related fixation biases. We tested associations between synonymous family-specific $CUB_{Chi/L}$ and $CUB_{Chi/L}$ $2f_{non\text{-}NAY}$(excluding the synonymous family being tested). Our proxy for MCP-related fixation biases is supported by previous findings of G- or C-ending major codons for $2f_{non\text{-}NAY}$ synonymous families in the six Drosophila species under analysis (Heger and Ponting 2007a; Vicario *et al.* 2007). We ranked genes by $CUB_{Chi/L}$ $2f_{non\text{-}NAY}$ and assigned genes into 15 bins (minimum 70,000 $2f_{non\text{-}NAY}$ codons per bin). $CUB_{Chi/L}$ for each bin was calculated from combined codon frequencies among CDS within the bin. We employed Spearman's rank correlation to assess the associations.

##### *Identifying OrthoFinder-orthologs among distantly related Drosophila species*

We identified potential 1-to-1 orthologs for pairs between *Dmel* and each of distantly related Drosophila species: *Dpse*, *Dwil*, *D. grimshawi* (*Dgri*), *D. mojavensis* (*Dmoj*), and *D. virilis* (*Dvir*). We ran OrthoFinder (Emms and Kelly 2015; version 2.5.2; downloaded on 15th March 2021) to obtain putative ortholog groups for 30 species from the genus Drosophila: *melanogaster* (*mel*), *simulans* (*sim*), *sechellia* (*sec*), *mauritiana* (*mau*), *teissieri* (*tei*), *yakuba* (*yak*), *santomea* (*san*), *erecta* (*ere*), *eugracilis* (*eug*), *suzukii* (*suz*), *subpullchrella* (*sup*), *biarmipes* (*bia*), *takahashii* (*tak*), *elegans* (*ele*), *rhopaloa* (*rho*), *ficusphila* (*fic*), *kikkawai* (*kik*), *ananassae* (*ana*), *bipectinata* (*bip*), *pseudoobscura* (*pse*), *persimilis* (*per*), *miranda* (*mir*), *obscura* (*obs*), *guanche* (*gua*), *subobscura* (*sub*), *willistoni* (*wil*), *grimshawi* (*gri*), *mojavensis* (*moj*), *virilis* (*vir*), and *serrata* (*ser*). Genome sequences, annotations, and download date are as follows: *mel*, FlyBase, 6.24, FB2018_05, 12 July 2019; *sim*, FlyBase, 2.02, FB2017_04, 9 March 2020; *sec*, NCBI, 101 (GCF_004382195.1), 13 October 2021; *mau*, NCBI, 100 (GCF_004382145.1), 11 March 2021; *tei*, NCBI, 100 (GCF_016746235.2), 15 October 2021; *yak*, FlyBase, 1.05, FB2016_05, 12 July 2019; *san*, NCBI, 101 (GCF_016746245.2), 13 October 2021; *ere*, FlyBase, 1.05, FB2016_05, 12 July 2019; *eug*, NCBI, 101 (GCF_000236325.1), 11 March 2021; *suz*, NCBI, 102 (GCF_013340165.1), 11 March 2021; *sup*, NCBI, 100 (GCF_014743375.2), 11 March 2021; *bia*, NCBI, 101 (GCF_000233415.1), 11 March 2021; *tak*, NCBI, 101 (GCF_000224235.1), 11 March 2021; *ele*, NCBI, 101

(GCF_000224195.1), 11 March 2021; *rho*, NCBI, 101 (GCF_000236305.1), 11 March 2021; *fic*, NCBI, 101 (GCF_000220665.1), 11 March 2021; *kik*, NCBI, 101 (GCF_000224215.1), 11 March 2021; *ana*, FlyBase, 1.06, FB2018_04, 16 October 2019; *bip*, NCBI, 101 (GCF_000236285.1), 11 March 2021; *pse*, FlyBase, 3.04, FB2018_05, 16 October 2019; *per*, NCBI, 101 (GCF_003286085.1), 13 October 2021; *mir*, NCBI, 102 (GCF_003369915.1), 11 March 2021; *obs*, NCBI, 100 (GCF_002217835.1), 11 March 2021; *gua*, NCBI, 100 (GCF_900245975.1), 11 March 2021; *sub*, NCBI, 100 (GCF_008121235.1), 11 March 2021; *wil*, NCBI, 101 (GCF_000005925.1), 13 October 2021; *gri*, NCBI, 103 (GCF_018153295.1), 13 October 2021; *moj*, NCBI, 102 (GCF_018153725.1), 13 October 2021; *vir*, NCBI, 103 (GCF_003285735.1), 13 October 2021; and *ser*, NCBI, 100 (GCF_002093755.1), 11 March 2021.

We extracted predicted CDS from genome sequences based on the annotations. For genes with multiple CDS isoforms, we used the longest CDS. We filtered CDS from trans-spliced genes and mitochondrial DNA-encoded genes. For data from NCBI, we filtered CDS marked as “LOW QUALITY PROTEIN”. We filtered CDS whose lengths are not divisible by three. However, in cases where NCBI provides coding frame information for partial CDS, we included CDS in the latter class after trimming one or two bases to eliminate incomplete codons. We extracted gene pairs that were the sole representatives of a given species pair within an OrthoFinder group (1-to-1 OrthoFinder orthologs). We did not process OrthoFinder groups containing multiple representatives from at least one of the six species.

We filtered some CDS and SI sequences using 1-to-1 OrthoFinder ortholog pairs of *Dmel* and each of non-*Dmel* species. We employed CDS and SI sequences from autosomal genes in the *Dmel* reference genome and those from 1-to-1 OrthoFinder orthologs in non-*Dmel* species. We excluded CDS with < 10 $2f_{non\text{-}NAY}$ codons as well as *Dmel* genes located in heterochromatic/low crossover regions and their 1-to-1 OrthoFinder orthologs in non-*Dmel* species. In addition, for *Dmel*, we filtered codons that include sites overlapping with transcribed regions of other genes and codons that are annotated as intron sites among at least one of transcript isoforms. We obtained 9119, 8081, 7886, 7759, 7934, and 7901 CDS for *Dmel*, *Dpse*, *Dwil*, *Dgri*, *Dmoj*, and *Dvir*, respectively.

### References

Akashi H., 1995 Inferring weak selection from patterns of polymorphism and divergence at “silent” sites in Drosophila DNA. Genetics 139: 1067–1076.

Akashi H., 1997 Distinguishing the effects of mutational biases and natural selection on DNA sequence variation. Genetics 147: 1989–1991.

Akashi H., 1999 Inferring the fitness effects of DNA mutations from polymorphism and divergence data: statistical power to detect directional selection under stationarity and free recombination. Genetics 151: 221–238.

Akashi H., W.-Y. Ko, S. Piao, A. John, P. Goel, *et al.*, 2006 Molecular evolution in the *Drosophila melanogaster* species subgroup: frequent parameter fluctuations on the timescale of molecular divergence. Genetics 172: 1711–1726. https://doi.org/10.1534/genetics.105.049676

Akashi H., P. Goel, and A. John, 2007 Ancestral inference and the study of codon bias evolution: implications for molecular evolutionary analyses of the *Drosophila melanogaster* subgroup. PLoS One 2: e1065. https://doi.org/10.1371/journal.pone.0001065

Anderson C. L., E. A. Carew, and J. R. Powell, 1993 Evolution of the *Adh* locus in the *Drosophila willistoni* group: the loss of an intron, and shift in codon usage. Mol. Biol. Evol. 10: 605–618. https://doi.org/10.1093/oxfordjournals.molbev.a040027

Assaf Z. J., S. Tilk, J. Park, M. L. Siegal, and D. A. Petrov, 2017 Deep sequencing of natural and experimental populations of Drosophila melanogaster reveals biases in the spectrum of new mutations. Genome Res. 27: 1988–2000. https://doi.org/10.1101/gr.219956.116

Bénitìere F., T. Lefébure, and L. Duret, 2024 Variation in the fitness impact of translationally optimal codons among animals. bioRxiv 2024.07.22.604600.

Chintapalli V. R., J. Wang, and J. A. T. Dow, 2007 Using FlyAtlas to identify better Drosophila melanogaster models of human disease. Nat. Genet. 39: 715–720. https://doi.org/10.1038/ng2049

Crow J. F., and M. Kimura, 1965 Evolution in Sexual and Asexual Populations. Am. Nat. 99: 439–450. https://doi.org/10.1086/282389

Efron B., 1979 Bootstrap Methods: Another Look at the Jackknife. The Annals of Statistics 7: 1–26. https://doi.org/10.1214/aos/1176344552

Emms D. M., and S. Kelly, 2015 OrthoFinder: solving fundamental biases in whole genome comparisons dramatically improves orthogroup inference accuracy. Genome Biol. 16: 1–14. https://doi.org/10.1186/s13059-015-0721-2

Felsenstein J., 1974 The evolutionary advantage of recombination. Genetics 78: 737–756.

Felsenstein J., 2004 PHYLIP (Phylogeny Inference Package) version 3.6. Distributed by the author. www. evolution. gs. washington. edu.

Fisher R. A., 1930 *The Genetical Theory of Natural Selection*. Clarendon Press, Oxford.

Fisher R. A., 1954 *Statistical Methods for Research Workers*. Oliver & Boyd, Edinburgh.

Fuller Z. L., G. D. Haynes, D. Zhu, M. Batterton, H. Chao, *et al.*, 2014 Evidence for stabilizing selection on codon usage in chromosomal rearrangements of *Drosophila pseudoobscura*. G3 4: 2433–2449. https://doi.org/10.1534/g3.114.014860

Galtier N., E. Bazin, and N. Bierne, 2006 GC-biased segregation of noncoding polymorphisms in Drosophila. Genetics 172: 221–228. https://doi.org/10.1534/genetics.105.046524

Glémin S., P. F. Arndt, P. W. Messer, D. Petrov, N. Galtier, *et al.*, 2015 Quantification of GC-biased gene conversion in the human genome. Genome Res. 25: 1215–1228. https://doi.org/10.1101/gr.185488.114

Haddrill P. R., and B. Charlesworth, 2008 Non-neutral processes drive the nucleotide composition of non-coding sequences in Drosophila. Biol. Lett. 4: 438–441. https://doi.org/10.1098/rsbl.2008.0174

Halligan D. L., A. Eyre-Walker, P. Andolfatto, and P. D. Keightley, 2004 Patterns of Evolutionary Constraints in Intronic and Intergenic DNA of Drosophila. Genome Res. 14: 273–279. https://doi.org/10.1101/gr.1329204

Halligan D. L., and P. D. Keightley, 2006 Ubiquitous selective constraints in the Drosophila genome revealed by a genome-wide interspecies comparison. Genome Res. 16: 875–884. https://doi.org/10.1101/gr.5022906

Hanson G., and J. Coller, 2018 Codon optimality, bias and usage in translation and mRNA decay. Nat. Rev. Mol. Cell Biol. 19: 20–30. https://doi.org/10.1038/nrm.2017.91

Heger A., and C. P. Ponting, 2007a Evolutionary rate analyses of orthologs and paralogs from 12 Drosophila genomes. Genome Res. 17: 1837–1849. https://doi.org/10.1101/gr.6249707

Heger A., and C. P. Ponting, 2007b Variable strength of translational selection among 12 Drosophila species. Genetics 177: 1337–1348. https://doi.org/10.1534/genetics.107.070466

Holm S., 1979 A Simple Sequentially Rejective Multiple Test Procedure. Scand. Stat. Theory Appl. 6: 65–70.

Hu T. T., M. B. Eisen, K. R. Thornton, and P. Andolfatto, 2013 A second-generation assembly of the *Drosophila simulans* genome provides new insights into patterns of lineage-specific divergence. Genome Res. 23: 89–98. https://doi.org/10.1101/gr.141689.112

Jackson B. C., J. L. Campos, P. R. Haddrill, B. Charlesworth, and K. Zeng, 2017 Variation in the Intensity of Selection on Codon Bias over Time Causes Contrasting Patterns of Base Composition Evolution in Drosophila. Genome Biol. Evol. 9: 102–123. https://doi.org/10.1093/gbe/evw291

Jackson B., and B. Charlesworth, 2021 Evidence for a force favoring GC over AT at short intronic sites in *Drosophila simulans* and *Drosophila melanogaster*. G3 11: jkab240. https://doi.org/10.1093/g3journal/jkab240

Kanaya S., Y. Yamada, Y. Kudo, and T. Ikemura, 1999 Studies of codon usage and tRNA genes of 18 unicellular organisms and quantification of Bacillus subtilis tRNAs: gene expression level and species-specific diversity of codon usage based on multivariate analysis. Gene 238: 143–155. https://doi.org/10.1016/s0378-1119(99)00225-5

Katoh K., K. Misawa, K. Kuma, and T. Miyata, 2002 MAFFT: a novel method for rapid multiple sequence alignment based on fast Fourier transform. Nucleic Acids Res. 30: 3059–3066. https://doi.org/10.1093/nar/gkf436

Kliman R. M., and J. Hey, 1993 Reduced natural selection associated with low recombination in *Drosophila melanogaster*. Mol. Biol. Evol. 10: 1239–1258. https://doi.org/10.1093/oxfordjournals.molbev.a040074

Kliman R. M., and J. Hey, 1994 The effects of mutation and natural selection on codon bias in the genes of Drosophila. Genetics 137: 1049–1056. https://doi.org/10.1093/genetics/137.4.1049

Kliman R. M., 2014 Evidence that natural selection on codon usage in *Drosophila pseudoobscura* varies across codons. G3 4: 681–692. https://doi.org/10.1534/g3.114.010488

Ko W.-Y., R. M. David, and H. Akashi, 2003 Molecular Phylogeny of the *Drosophila melanogaster* Species Subgroup. J. Mol. Evol. 57: 562–573. https://doi.org/10.1007/s00239-003-2510-x

Kopp A., and J. R. True, 2002 Phylogeny of the Oriental *Drosophila melanogaster* Species Group: A Multilocus Reconstruction, (J. Whitfield, Ed.). Syst. Biol. 51: 786–805. https://doi.org/10.1080/10635150290102410

Kurtz S., A. Phillippy, A. L. Delcher, M. Smoot, M. Shumway, *et al.*, 2004 Versatile and open software for comparing large genomes. Genome Biol. 5: 1–9. https://doi.org/10.1186/gb-2004-5-2-r12

Liu Y., Q. Yang, and F. Zhao, 2021 Synonymous but Not Silent: The Codon Usage Code for Gene Expression and Protein Folding. Annu. Rev. Biochem. 90: 375–401. https://doi.org/10.1146/annurev-biochem-071320-112701

Mantel N., and W. Haenszel, 1959 Statistical aspects of the analysis of data from retrospective studies of disease. J. Natl. Cancer Inst. 22: 719–748.

Mantel N., 1963 Chi-Square Tests with One Degree of Freedom; Extensions of the Mantel-Haenszel Procedure. J. Am. Stat. Assoc. 58: 690–700. https://doi.org/10.1080/01621459.1963.10500879

Marais G., D. Mouchiroud, and L. Duret, 2001 Does recombination improve selection on codon usage? Lessons from nematode and fly complete genomes. Proc. Natl. Acad. Sci. U. S. A. 98: 5688–5692. https://doi.org/10.1073/pnas.091427698

Marais G., D. Mouchiroud, and L. Duret, 2003 Neutral effect of recombination on base composition in Drosophila. Genet. Res. 81: 79–87. https://doi.org/10.1017/s0016672302006079

Maside X., A. W. Lee, and B. Charlesworth, 2004 Selection on Codon Usage in *Drosophila americana*. Curr. Biol. 14: 150–154. https://doi.org/10.1016/j.cub.2003.12.055

Matsumoto T., H. Akashi, and Z. Yang, 2015 Evaluation of Ancestral Sequence Reconstruction Methods to Infer Nonstationary Patterns of Nucleotide Substitution. Genetics 200: 873–890. https://doi.org/10.1534/genetics.115.177386

Matsumoto T., A. John, P. Baeza-Centurion, B. Li, and H. Akashi, 2016 Codon Usage Selection Can Bias Estimation of the Fraction of Adaptive Amino Acid Fixations. Mol. Biol. Evol. 33: 1580–1589. https://doi.org/10.1093/molbev/msw027

Matsumoto T., and H. Akashi, 2018 Distinguishing Among Evolutionary Forces Acting on Genome-Wide Base Composition: Computer Simulation Analysis of Approximate Methods for Inferring Site Frequency Spectra of Derived Mutations in Recombining Regions. G3 8: 1755–1769. https://doi.org/10.1534/g3.117.300512

Muller H. J., 1932 Some Genetic Aspects of Sex. Am. Nat. 66: 118–138. https://doi.org/10.1086/280418

Nei M., and T. Gojobori, 1986 Simple methods for estimating the numbers of synonymous and nonsynonymous nucleotide substitutions. Mol. Biol. Evol. 3: 418–426.

https://doi.org/10.1093/oxfordjournals.molbev.a040410

Parsch J., S. Novozhilov, S. S. Saminadin-Peter, K. M. Wong, and P. Andolfatto, 2010 On the Utility of Short Intron Sequences as a Reference for the Detection of Positive and Negative Selection in Drosophila. Mol. Biol. Evol. 27: 1226–1234. https://doi.org/10.1093/molbev/msq046

Poh Y.-P., C.-T. Ting, H.-W. Fu, C. H. Langley, and D. J. Begun, 2012 Population Genomic Analysis of Base Composition Evolution in *Drosophila melanogaster*. Genome Biol. Evol. 4: 1245–1255. https://doi.org/10.1093/gbe/evs097

Pollard D. A., V. N. Iyer, A. M. Moses, and M. B. Eisen, 2006 Widespread Discordance of Gene Trees with Species Tree in Drosophila: Evidence for Incomplete Lineage Sorting, (B. F. McAllister, Ed.). PLoS Genet. 2: e173. https://doi.org/10.1371/journal.pgen.0020173

Powell J. R., E. Sezzi, E. N. Moriyama, J. M. Gleason, and A. Caccone, 2003 Analysis of a Shift in Codon Usage in Drosophila. J. Mol. Evol. 57: S214–S225. https://doi.org/10.1007/s00239-003-0030-3

Rogers R. L., J. M. Cridland, L. Shao, T. T. Hu, P. Andolfatto, *et al.*, 2014 Landscape of standing variation for tandem duplications in *Drosophila yakuba* and *Drosophila simulans*. Mol. Biol. Evol. 31: 1750–1766. https://doi.org/10.1093/molbev/msu124

Sharp P. M., and W. H. Li, 1987 The codon Adaptation Index--a measure of directional synonymous codon usage bias, and its potential applications. Nucleic Acids Res. 15: 1281–1295.

Shields D. C., P. M. Sharp, D. G. Higgins, and F. Wright, 1988 Silent sites in Drosophila genes are not neutral: evidence of selection among synonymous codons. Mol. Biol. Evol. 5: 704–716. https://doi.org/10.1093/oxfordjournals.molbev.a040525

Singh N. D., P. F. Arndt, and D. A. Petrov, 2005 Genomic heterogeneity of background substitutional patterns in *Drosophila melanogaster*. Genetics 169: 709–722. https://doi.org/10.1534/genetics.104.032250

Singh N. D., V. L. Bauer DuMont, M. J. Hubisz, R. Nielsen, and C. F. Aquadro, 2007 Patterns of mutation and selection at synonymous sites in Drosophila. Mol. Biol. Evol. 24: 2687–2697. https://doi.org/10.1093/molbev/msm196

Singh N. D., P. F. Arndt, A. G. Clark, and C. F. Aquadro, 2009 Strong Evidence for Lineage and Sequence Specificity of Substitution Rates and Patterns in Drosophila. Mol. Biol. Evol. 26: 1591–1605. https://doi.org/10.1093/molbev/msp071

Takano-Shimizu T., 2001 Local changes in GC/AT substitution biases and in crossover frequencies on Drosophila chromosomes. Mol. Biol. Evol. 18: 606–619. https://doi.org/10.1093/oxfordjournals.molbev.a003841

Tavaré S., 1986 Some Probabilistic and Statistical Problems in the Analysis of DNA Sequences, pp. 57–86 in *Some Mathematical Questions in Biology: DNA Sequence Analysis*, edited by Miura R. M. Lectures on Mathematics in the Life Sciences.

Thurmond J., J. L. Goodman, V. B. Strelets, H. Attrill, L. S. Gramates, *et al.*, 2018 FlyBase 2.0: the next generation. Nucleic Acids Res. 47: D759–D765. https://doi.org/10.1093/nar/gky1003

Vicario S., E. N. Moriyama, and J. R. Powell, 2007 Codon usage in twelve species of Drosophila. BMC Evol. Biol. 7: 1–17. https://doi.org/10.1186/1471-2148-7-226

Wong A., J. D. Jensen, J. E. Pool, and C. F. Aquadro, 2007 Phylogenetic incongruence in the *Drosophila melanogaster* species group. Mol. Phylogenet. Evol. 43: 1138–1150. https://doi.org/10.1016/j.ympev.2006.09.002

Wright S., 1969 *Evolution and The Genetics of Populations: Vol. 2. The Theory of Gene Frequencies*. University of Chicago Press.

Wright F., 1990 The "effective number of codons" used in a gene. Gene 87: 23–29. https://doi.org/10.1016/0378-1119(90)90491-9

Wright S. I., C. B. K. Yau, M. Looseley, and B. C. Meyers, 2004 Effects of gene expression on molecular evolution in *Arabidopsis thaliana* and *Arabidopsis lyrata*. Mol. Biol. Evol. 21: 1719–1726. https://doi.org/10.1093/molbev/msh191

Yang Z., 2007 PAML 4: phylogenetic analysis by maximum likelihood. Mol. Biol. Evol. 24: 1586–1591. https://doi.org/10.1093/molbev/msm088

Yıldırım B. C., and C. Vogl, 2024 The influence of GC-biased gene conversion on nonadaptive sequence evolution in short introns of *Drosophila melanogaster*. J. Evol. Biol. 37: 383–400. https://doi.org/10.1093/jeb/voae015

Zaborske J. M., V. L. B. DuMont, E. W. J. Wallace, T. Pan, C. F. Aquadro, *et al.*, 2014 A nutrient-driven tRNA modification alters translational fidelity and genome-wide protein coding across an animal genus. PLoS Biol. 12: e1002015. https://doi.org/10.1371/journal.pbio.1002015
